# Salmonids elicit an acute behavioral response to heterothermal environments

**DOI:** 10.1101/2024.05.03.592389

**Authors:** Robert Naudascher, Stefano Brizzolara, Jonasz Slomka, Robert M. Boes, Markus Holzner, Luiz G. M. Silva, Roman Stocker

## Abstract

Most fish species are ectothermic and rely on behavioral strategies to control their body temperature in heterothermal environments. Both thermotaxis and thermokinesis have been suggested as important underlying mechanisms. However, to what extent these behaviors allow fish to respond to rapid (timescales of minutes) and strong thermal disturbances, like those caused by anthropogenic water releases into natural freshwater systems, is poorly understood. Here, we quantify this response for a salmonid species with a novel laboratory approach coupled with image-based tracking. We exposed brown trout parr (*Salmo trutta*), acclimated to 12 °C, to rapidly forming cold- and warm-water interfaces with temperatures ranging from 4 to 20 °C. We found that fish actively avoided colder water (≤8 °C) through a rapid response that combined thermotaxis and thermokinesis. In contrast, fish did not avoid warmer water and frequently crossed interfaces having temperature contrasts of up to 8 °C. This study shows that brown trout parr swiftly deploy multiple behavioral strategies to minimize exposure to cold water and take advantage of warm water, illustrating their capability to cope with rapidly occurring thermal alterations.

## Introduction

Water temperature constitutes a key physical property of riverine ecosystems. It has been described as the ’abiotic master factor’ for fishes^1,2^, strongly affecting their energy balance and behavior^3,4^. In natural habitats, temperature may change on a variety of spatial and temporal scales. Thermal gradients occur at river confluences, at locations where rivers flow into lakes, and within vertically stratified lakes. As most fish species are ectotherms, their body temperature changes according to the temperature of the water they occupy^5^. Because temperature influences the rates of most physiological processes, rapid warming or cooling can affect fish performance traits, including metabolic rates, swimming ability, and thermal tolerance^6^.

Because they lack physiological means to regulate body temperature, fish have two main choices when exposed to heterothermal environments (i.e., environments in which temperature varies over space and/or time): acclimate physiologically to changes in local temperature (physiological plasticity^7,8^) or adopt specific behaviors to regulate their body temperature (behavioral thermoregulation^5,9^). Behavioral thermoregulation allows fish to thrive in environments where temperature fluctuates, by maximizing the time spent in areas where temperature is within their optimum performance range and by avoiding temperature extremes. While fish may generally adjust to gradual temperature changes through physiological plasticity, the risk of stress increases for more rapid temperature changes^10^ and in this situation, behavioral thermoregulation may be of greater importance.

Fish were found to actively seek optimal temperatures in their environment through movement behavior 50 years ago^11^. Recent experiments to determine the mechanisms underlying behavioral thermoregulation have identified both thermokinetic and thermotactic components^12,13^. Thermokinesis refers to a change in the rate of movement of fish in response to temperature. One explanation for thermokinesis is that a change in body temperature impacts the metabolic rate, which in turn affects movement activity. In this scenario, thermal sensing is not required. For instance, zebrafish larvae (*Danio rerio*) increase their turn frequency, turn amplitude and swimming speed when water temperature is increased from 18°C to 32 °C ^14^. This thermokinetic behavior can explain how zebrafish larvae avoid warmer water, but not how they avoid colder water.

Thermotaxis, in contrast, is the directed movement along a temperature gradient, toward warmer or colder water, and requires the ability to sense a temperature gradient^12,13^. Fish may achieve thermotaxis either by sensing a thermal gradient spatially along their body or temporally over a given time window as they swim. The response to a thermal gradient can then be either deterministic, through directed movement along the gradient, or biased, for example through the increase in the turning rate when moving down the gradient, analogous to bacterial chemotaxis^15,16^. Thermotaxis has been observed in a number of vertebrate organisms^17–19^. While the thermal preference of fish is a well-established field of research, very few quantitative studies of behavioral mechanisms allowing fish to seek their preferendum (i.e. thermotaxis) have been conducted in fish. Among these is work showing that larval zebrafish integrate thermal information over a 500-ms time window^5^ and bias the magnitude and direction of their turns to favor movement toward colder regions.

Past studies have investigated the long-term (hours) behavioral response of fish to stationary thermal gradients. In contrast, very little is known about the thermoregulatory behavior in response to the exposure to sharp, rapidly occurring thermal gradients associated for example with anthropogenic disturbances^12^. Here, we report experiments to quantify the thermoregulatory behavior of a salmonid in its early life-stages in response to the rapid formation of sharp vertical temperature gradients generated using a lock-release approach. In particular, we identified a knowledge gap associated with the rapid intrusion of cold and warm water into the fishs habituated surrounding. Thereby we advance our understanding of thermoregulatory behavior in vertically stratified heterothermal environments. Salmonids are among the most common and well-studied families of freshwater fish in the northern hemisphere, yet we know little about their thermoregulatory behavior^20^. All salmonids spawn in the shallow gravel beds of freshwater headstreams and spend most of their juvenile stage in rivers and small creeks, where temperature regimes are often strongly affected by human activities^21^. Due to their small body size, the early life-stages of fish gain or lose heat rapidly, making them particularly sensitive to thermal disturbances^22^.

The natural thermal regime of rivers can be significantly altered by human activities^23^, including the operation of storage hydropower (i.e., induced thermopeaking)^21,24^, the use of water for cooling in thermal power plants^25^, flow regulation at artificial barriers^26,27^, urbanization^28^ and climate change^29,30^. If the local water temperature changes too rapidly, this might trigger an avoidance response, driving fish to abandon habitats^20^. For example, recent observations of the impact of thermopeaking show that a sudden decrease in water temperature (by 2.5 °C in 10 min) led to significantly higher downstream drift of juvenile European grayling (*Thymallus thymallus*) than during events with increasing water temperature (1). However, research on this topic remains scarce^31^ and the mechanisms underlying rapid behavioral thermoregulation have remained unclear.

Human-induced thermal alterations in freshwater systems can be unnaturally rapid and strong^24^. For example, the water temperature downstream of hydropower facilities can decrease by up to 6 °C within minutes in spring and summer, and increase strongly (up to 4 °C) in winter^24^. During spring and summer, cold thermopeaking may cause significant mortality, as it coincides with the presence of early life stages of salmonids and cyprinids (32). Rapid thermal changes (multiple degrees Celsius over minutes) are likely challenging for fish, partly because their natural occurrence over evolutionary timescales might have been rare. For instance, if they are unable to relocate upon being exposed to cold water, fish can become cold-shocked, a lethargic state of slowed metabolism, reduced oxygen uptake and muscle stiffness, which results in reduced swimming activity and increased risk of predation^10^.

The primary aim of our study was to quantify how brown trout parr (*Salmo trutta*), hereafter trout, respond to rapid (i.e., over timescales τ_PHYS_ of minutes) thermal disturbances. To mimic the rapid release of thermal effluent into a freshwater system, we developed a laboratory setup that uses the lock-release of a gravity current to rapidly expose fish acclimated to 12 ° (their assumed thermal preferendum) to a warmer or colder body of water. By continuously tracking individual fish, we quantified their response to incoming water of different temperatures (*T*_incoming_ = 4–20 °C). We hypothesized that after rapid exposure fish would avoid the incoming colder or warmer water across all treatments and that they would do so through a combination of thermokinesis and thermotaxis. To test this hypothesis, we developed a novel imaging approach that allowed simultaneous tracking of the thermal interface and the swimming behavior of individual fish at high spatio-temporal resolution. Our results indicate that fish use both thermokinetic and thermotactic behaviors to rapidly and effectively avoid incoming cold water, whereas they perform frequent excursions into incoming warm water, possibly as a strategy to gradually elevate their body temperature.

## Results

To test the hypothesis that strong thermal gradients trigger an avoidance response in trout, we exposed individuals (in groups of four for each replicate experiment) to heterothermal conditions in a laboratory setup. The setup consisted of a lock-exchange tank initially divided into equal halves containing water at two different temperatures (Fig. 1*A*). To avoid behavioral bias due to visual cues, the tank was shielded with white semi-transparent material (except for the front and the water surface). The experiments were conducted in a dark room and the tank was illuminated with a light source from behind (Supplementary Fig. 1 the left side of the tank, fish were acclimated for 15 min to water at 12 °C, which was assumed to be close their thermal preferendum^32,33^. This water temperature was also equal to the husbandry tank at our experimental facility. The right side of the tank contained water at a different temperature (colder or warmer), dyed blue (Patent blue V sodium salt, Sigma Aldrich) to allow visualization of the thermal interface (Supplementary Fig. 2). Experiments included four cold treatments (TR1, 10 °C; TR2, 8 °C; TR3, 6 °C; TR4, 4 °C) and four warm treatments (TR6, 14 °C; TR7, 16 °C; TR8, 18 °C; TR9, 20 °C) (Supplementary Table 1). After the 15 min acclimation time, the vertical lock separating the two halves of the tank was rapidly lifted, resulting in a gravity current with warmer water intruding along the top of the tank and colder water intruding in the opposite direction along the bottom. The initial gravity current reached the end-wall of the tank after 1–2 min, creating a sharp thermal interface vertically separating the colder water in the bottom part of the tank (*T*_bottom_) from the warmer water in the top part (*T*_top_) (Fig. 1*B*; Supplementary Fig. 2). Owing to the oscillations triggered by the lock release, the vertical location of the thermal interface was not constant in time (Supplementary Movies 1 and 2), but instead oscillated around the half depth of the tank, with oscillations dissipating over time (Supplementary Fig. 3).

**Fig. 1.**
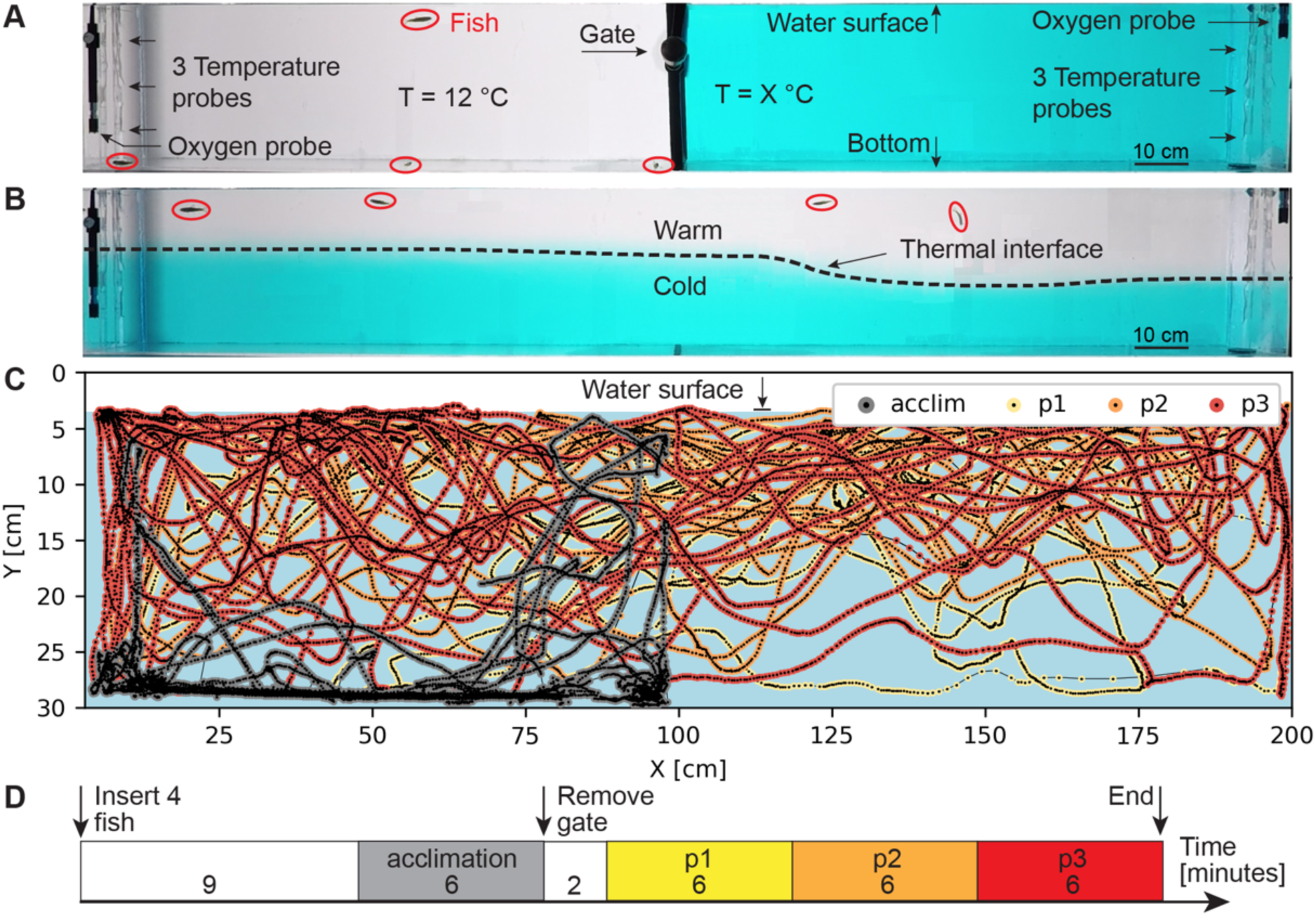
Lock-exchange tank used to expose fish to an acute heterothermal environment. (*A*) Side-view of the experimental tank during the acclimation phase (15 min), with fish in the left compartment exposed to water at their acclimation temperature (12 °C) separated by a closed gate from the dyed water of a different temperature in the right compartment. Red ellipses indicate the locations of four trout parr tested simultaneously. Fish were tracked continuously based on video recordings of the lateral view of the entire tank. (*B*) The treatment phase was initiated by rapidly removing the gate, allowing the cold water to slowly intrude below the warmer water (1–2 min), thereby exposing the fish to a heterothermal environment. A sharp thermal interface (dashed line) separated upper, warmer water from lower, colder water. Within the first two minutes, the thermal interface moved laterally and afterwards spanned the entire lateral extent of the tank. Over time, the interface increasingly stabilized around the central depth of the tank. The movement of the interface was much slower than fish swimming speeds (Supplementary Figs. 3 and 20). The frame used here for illustration was captured in a cold-water treatment, 11 min after gate removal. (*C*) Trajectory of a single fish over 18 min (treatment TR3, *T*_bottom_ = 6 °C, *T*_top_ = 12 °C). The trajectory is color-coded according to the temporal phase of the experiment (see *D*). (*D*) The different phases of the experiments. After 15 min acclimation, the gate was removed and it took approximately 2 min to establish a horizontal thermal interface spanning the entire tank length. To allow the analysis of temporal trends in fish behavior, the dataset was divided into four periods of equal duration: acclimation, and experimental phases p1–p3. For a list of treatments, see Supplementary Table 1.

The lateral view of the entire tank was continuously recorded for 35 min with a camera at 24 frames per second, starting four minutes after releasing the fish. Using image analysis, we obtained the two-dimensional projection of the trajectory of each fish over the entire duration of the experiment (Fig. 1*C*), while simultaneously tracking the instantaneous location of the thermal interface (Supplementary Fig. 4). In this manner, we determined the water temperature experienced by each fish at each point in time. Trajectories were analyzed for the last 6 min of the acclimation period and for the 18-min treatment period, which we divided into three phases of 6 min each (treatment phases p1, p2, and p3; Fig. 1*D*). The depth occupancy was computed from the vertical position of the fish *Y(t)* and normalized to the range *Y*_norm_ = 0–1, with 1 being the maximum depth (28 cm) of the tank. To quantitatively analyze fish movement behavior in relation to the thermal interface, each trajectory was binarized in time based on whether at a given instant a fish was in the warm or the cold water, as determined from the instantaneous vertical distance to the thermal interface, *D(t)* (Fig. 2*A,B;* Supplementary Movies 1 and 3; *Materials and Methods*).

**Fig. 2.**
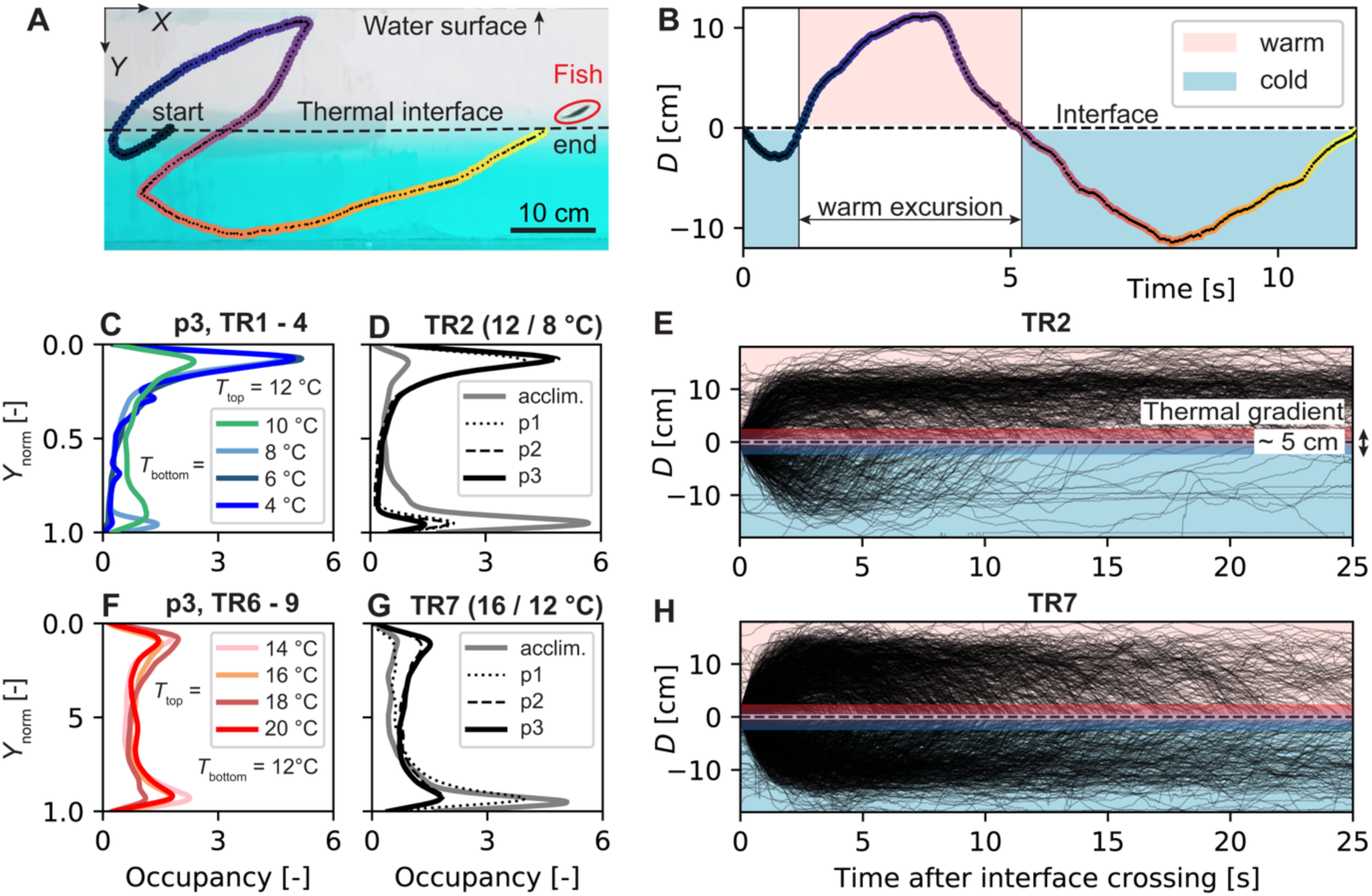
Fish avoid the cold, but not the warm. (*A*) Side-view of the (two-dimensional) swimming trajectory of a fish (colored line) in relation to the thermal interface (dashed black line). Black dots along the trajectory indicate the center of gravity of the fish during an example segment of 12 s. Colors indicate elapsed time, from blue to yellow. (*B*) Definition of warm- and cold-water excursions within a trajectory based on the fish’s instantaneous vertical distance to the thermal interface, *D*(*t*). An excursion starts and ends when the fish crosses the thermal interface. (*C* and *F*) Probability density functions of normalized fish depth *Y*_norm_ during the final 6 min (p3) of the experiments for all cold-water (TR1–TR4, *C*) and all warm-water (TR6–TR9, *F*) treatments. Data was aggregated at the treatment level, hence each line represents data from 20 fish, tested in groups of 4 individuals in 5 separate experiments. *T*_bottom_ and *T*_top_ indicate the water temperature below and above the thermal interface. The depth was normalized so that *Y*_norm_ = 0 is the water surface and *Y*_norm_ = 1 indicates the bottom of the tank. (*D* and *G*) Probability density functions of fish depth for TR2 (*D*) and TR7 (*G*), separated according to the four phases of analysis (acclimation phase and experimental phases p1–p3; see Fig. 1D). Data were aggregated at the treatment level as in *C* and *F*. Data for other treatments are shown in Supplementary Fig. 5. (*E* and *H*) Vertical distance to the interface, *D*, as a function of time for each cold-water (light blue shaded region) and warm-water excursion (light pink shaded region) for TR2 (*N* = 980) (*E*) and TR7 (*N* = 430) (*H*) aggregated for all three experimental phases (p1–p3). *N* is the total number of cold-water excursions performed by the 20 fish. The vertical extent of the thermal gradient thickness is indicated as blue and red (see Methods). shown in red Data for other treatments are shown in Supplementary Figs. 6 and 7. For a list of treatments, see Supplementary Table 1.

### Trout parr avoid cold water by biasing their vertical turns and do not avoid warm water

Fish showed a clear thermotactic avoidance behavior of incoming cold water for *T*_bottom_ ≤ 8 °C. During phase p3 in the three coldest treatments, TR2 (*T*_bottom_ = 8 °C), TR3 (*T*_bottom_ = 6 °C) and TR4 (*T*_bottom_ = 4 °C), fish predominantly occupied the upper half of the tank (*Y*_norm_ = 0–0.5) and avoided the colder water in the bottom half (Fig. 2*C*; see also Supplementary Fig. 9). From phase p1 to p3, fish spent 75–95% of the time above the thermal interface (Supplementary Fig. 8A). This represents a substantial change of the preferential depth occupancy compared to the acclimation phase, in which fish spent 50–90% of the time in the lower half of the tank (*Y*_norm_ = 0.5–1; Supplementary Fig. 9). The response was rapid, with avoidance of the cold water being apparent already in phase p1 and persisting through to the end of the experiment (TR2 in Fig. 2*D*; all cold-water treatments in Supplementary Fig. 5 *A-D*). In contrast, incoming cold water in TR1 (*T*_bottom_ = 10 °C) – i.e. only 2 °C colder than the acclimation temperature – did not trigger an avoidance behavior: the depth occupancy showed a bimodal distribution with peaks in both the warmer and colder water (Fig. 2*C*).

Fish did not avoid warmer water in any of the warm-water treatments (TR6–TR9). Instead, fish transitioned to a bimodal depth occupancy, with comparable time spent above and below the interface (Fig. 2*F*; Supplementary Fig. 8B). This bimodal behavior was especially pronounced in the later phases, p2 and p3 (TR7; Fig. 2*G;* all warm-water treatments in Supplementary Fig. 5 *E*–*H*), whereas in phase p1 the depth occupancy was similar to that exhibited during the acclimation period, with 50– 70% of the time spent in the lower, colder water (Supplementary Fig. 10). Thus, over time fish increasingly occupied warmer water across all warm-water treatments, reducing the fraction of time spent below the interface during phase p3 to 30–50% (Supplementary Figs 8B and 10).

Further analysis of the response to cold water treatments revealed a strong asymmetry between warm-water and cold-water excursions, defined as the periods of time during which a fish was above or below the thermal interface, respectively. This asymmetry was present in treatments with *T*_bottom_ ≤ 8 °C: TR2 (Fig. 2*E*), TR3 and TR4 (Supplementary Fig. 6C,D), but not TR1 (Supplementary Fig. 6A). In cold treatments TR2, TR3 and TR4, cold-water excursions were considerably shorter (91–94% shorter than 10 s) than warm-water excursions (38–49% shorter 10 s; Supplementary Table 3). In contrast, during all warm treatments, warm and cold excursions had similar durations (Fig. 2*H*, Supplementary Fig. 7).

To determine the behavioral mechanism underlying cold-water avoidance, we analyzed the fine-scale features of the fish trajectories as a function of the position of the fish relative to thermal interface. We identified upward and downward turning events within each trajectory, based on when the vertical velocity component *v*_y_ changed sign (Fig. 3*A, B*; *Materials and Methods*; Supplementary Fig. 11). For each turning event, we computed the corresponding vertical distance *D̂* to the thermal interface. We also quantified the median number of cold-water excursions *Ñ* of the four fish within each experimental replicate. We found that fish reduced their tendency to cross the thermal interface and rapidly redirected their swimming when in the colder water. In cold-water treatments TR2-TR4 (*T*_bottom_ ≤ 8 °C) the median number of interface crossings was significantly lower than in TR1 (*T*_bottom_ = 10 °C) (ANOVA and post hoc pairwise *t*-tests – p < 0.01, Fig. 3*C*). In TR2-TR4, the probability of upward turning *ρ*_up_ exceeded the probability of downward turning *ρ*_down_ for the entire region below the interface (-20 cm < *D̂* < 0 cm; Fig. 3*D,F,* Supplementary Fig. 12). This difference in probabilities (*ρ*_up_ - *ρ*_down_) was particularly large in the region immediately above and below the interface (-5 cm < *D̂* < 5 cm; Fig. 3*F*) and is a hallmark of a thermotactic behavior.

**Fig. 3.**
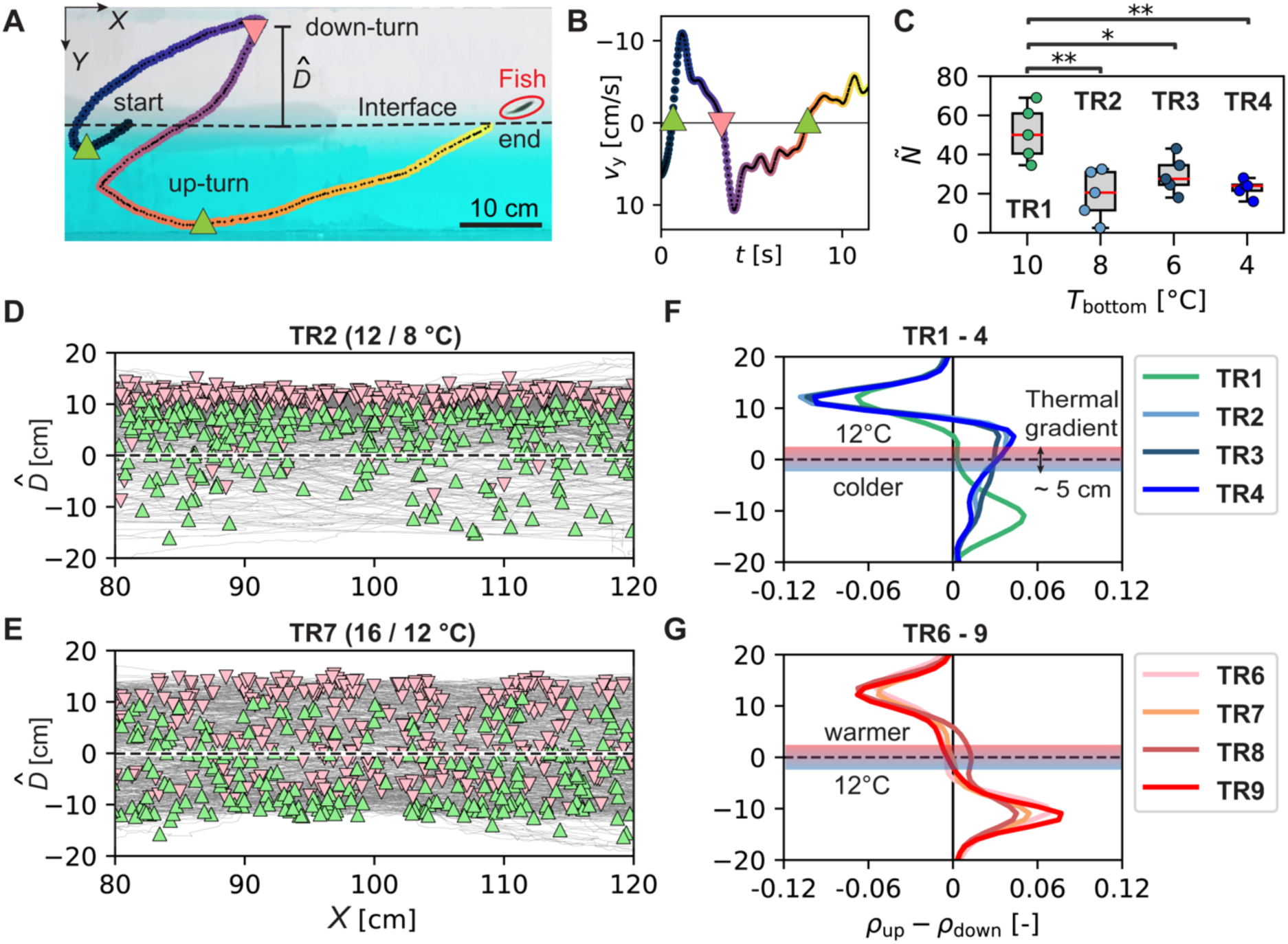
Fish avoid cold water by directional upward turning. (*A*) Example trajectory (12 s) with identified vertical turning points (pink triangles, downward; green triangles, upward). (*B*) Turning points were identified as sign changes of the time series of the vertical velocity component, *v*_y_(*t*). The same trajectory as in A is shown. (*C*) Boxplots showing the number of cold-water excursions for each cold temperature treatment. Each dot represents the median number of cold-water excursions *Ñ* of four individuals tested simultaneously within one experiment. Red lines indicate the median of 5 experiments. Whiskers extent to the full range of observations. The temperature treatments TR2 and TR4 differ significantly (ANOVA, *p* < 0.001), with pairwise *t*-test results as indicated (* 0.01 < *p* ≤ 0.05; ** 0.001 < *p* ≤ 0.01, see also Supplementary Table 18-20). (*D* and *E*) Swimming trajectories (gray lines) and vertical turning points (color-coded as in panel A) plotted with respect to the vertical distance *D̂* to the thermal interface (black dashed line) for cold-water treatment TR2 (*D*) and warm-water treatment TR7 (*E*). The trajectories shown are from the central region (80 < *X* < 120 cm) of the tank, for 20 fish (5 x 4 fish per experiment). Turning points were filtered based on a vertical displacement threshold of d*z* > 3 cm to detect turning activity that resulted in significant vertical displacement (Supplementary Fig. 14). Data for other treatments are shown in Supplementary Figs. 12 and 13. (*F* and *G*) Difference of probability density distributions of upward turns (*ρ*_up_) and downward turns (*ρ*_down_) for the entire width of the tank (0 < *X* < 200 cm) as a function of the distance from the thermal interface (located at *D̂* = 0), for all cold (*F*) and warm treatments (*G*). The same y-axes as in panels *D* and *E* apply. The tank bottom is located at around -10 cm > *D̂* > -20 cm, while the water surface is located at around 10 cm < *D̂* < 20 cm (the variability being related to the initial period of time in which the thermal interface oscillates). The underlying normalized histograms are in Supplementary Figs. 14 *C* and *E*. The vertical extent of the thermal gradient thickness is indicated as blue and red (see Methods).

This thermotactic avoidance behavior was absent in TR1 and all warm treatments, where the probabilities of upward and downward turning were similar (*ρ*_up_ - *ρ*_down_ ≈ 0) close to the interface (-5 cm < *D̂* < 5 cm) (Fig. 3*E,G*). In these treatments, the probability of upward turning peaked below the thermal interface, close to the bottom of the tank (-15 cm < *D̂* < -10 cm; Fig. 3*G*), while the probability of downward turning peaked above the thermal interface, close to the water surface (10 cm < *D̂* < 15 cm; Fig. 3*E, G*, Supplementary Fig. 13 for TR6–TR9).

### Fish modulate the duration and depth of cold-water excursions as a function of cold-water temperature

In cold treatments fish did not exclusively stay within the upper warmer water layer, but engaged in occasional exploratory behavior of the underlying, colder water. In particular, fish did not reduce the frequency of cold-water excursions over time (Supplementary Fig. 15A). During these excursions, they crossed the interface downwards and thus experienced a sudden decrease in temperature. We found that fish modulated these cold-water excursions as a function of the temperature of the colder water. The duration *Ω* (s) and depth *D*_max_ (the maximum vertical distance below the thermal interface) of cold-water excursions both decreased (Fig. 4*A*) with decreasing temperature of the lower layer (*T*_bottom_). This trend was statistically significant for both the median duration *Ω͌* (Fig. 4*B)* and the median depth *D͌*_max_ (Fig. 4*C)* across treatments (Mann–Kendall test, *p* < 0.001, Supplementary Table 6). As *T*_bottom_ decreased from 10 °C to 4 °C (TR1 to TR4), the median excursion duration decreased by 57% (from *Ω͌* = 6 s to *Ω͌* = 2.6 s) and the median excursion depth decreased by 57% (from *D͌*_max_ = 9.9 cm to *D͌*_max_ = 4.3 cm, Supplementary Table 5).

**Fig. 4.**
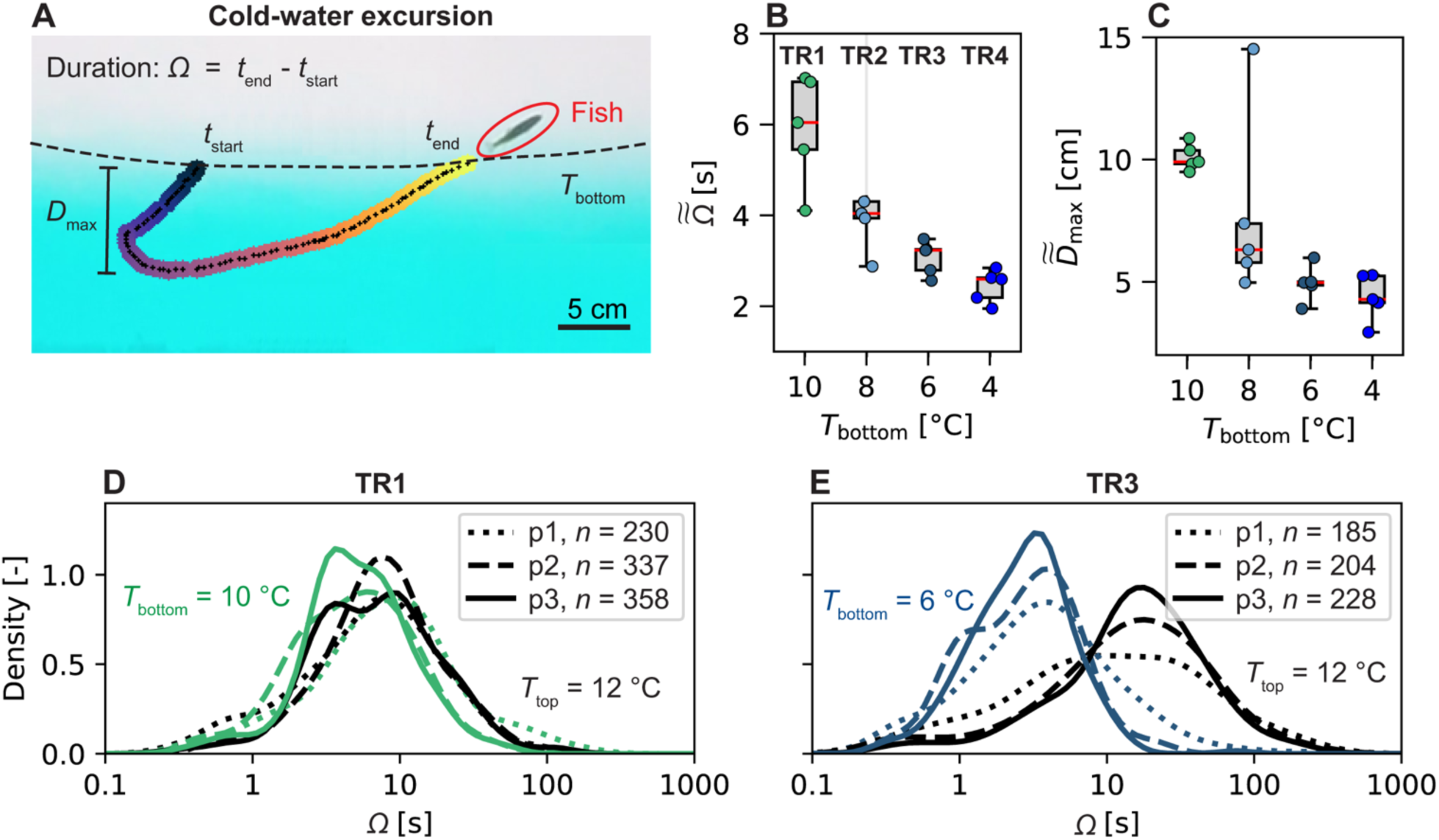
Fish reduce the duration and depth of cold-water excursions in larger measure the colder the water temperature below the thermal interface. (A) Fish perform cold-water excursions in which they cross the thermal interface and swim into colder water, then turn upward to return to the warmer water. Individual cold excursions were characterized by their duration, 𝛺, and the maximum vertical distance to the thermal interface, *D*_max_. (*B*) Fish exhibit shorter durations of cold-water excursions when encountering lower water temperatures below the interface (*T*_bottom_). The decreasing trend was statistically significant (Mann–Kendall test: for *Ω͌*, *p* < 0.001; for *D͌*_max_, *p* < 0.001; Supplementary Table 6). Boxplots show the median duration of cold-water excursions for fish groups across all phases (p1, p2 and p3) within each treatment. Points represent the median of a single experiment. TR2, one outlier not shown (out of range). (*C*) For colder water temperatures, fish penetrate less deeply into the cold lower water. The decreasing trend was statistically significant (Mann–Kendall test: *D͌*_max_, *p* < 0.001; Supplementary Table 6). Underlying distributions and histograms are shown in Supplementary Fig. 18. Equivalent plots for warm treatments are shown in Supplementary Fig. 17A,B. (*D* and *E*) Probability density distributions of the durations *Ω* of warm- and cold-water excursions for TR1 (*D*) and TR3 (*E*), respectively. The distributions of the warm- and cold-water excursion durations become increasingly distinct over exposure time for TR3 but not TR1. Excursions at 12 °C (above the interface) are shown in black, while excursions in the lower, colder water are depicted in the respective color. Treatment phases are indicated by line type (p1–p3). Values of *n* indicate the total number of excursions of each type within that phase for 20 fish (as the excursions alternate, *n* is equal for cold and warm excursions). Underlying histograms of *D* and *E* and other treatments are shown in Supplementary Fig. 19 and 16.

To understand whether this exploratory behavior of the colder water changed over the 18-min duration of the experiments, we analyzed the distribution of the excursion duration *Ω* of warm- and cold-water excursions separately for each phase (p1, p2 and p3). For TR1 (*T*_bottom_ = 10 °C, *T*_top_ = 12 °C), cold and warm water excursions showed almost identical distributions of their durations, and the distributions changed little over time across the three phases (*Ω̃*_warm_ = 6.7 s, 7.5 s, 7.3 s and *Ω̃*_cold_ = 8.4 s, 4.9 s, 5.1 s, for p1, p2, p3 respectively; Fig. 4*D*). The same conclusions apply to all warm treatments (TR6–TR9; Supplementary Fig. 16 *E–H*). In stark contrast, in cold-treatment TR3 the distributions of the durations of cold-water excursions (into *T*_bottom_ = 6 °C) and warm-water excursions (into *T*_top_ = 12 °C) are clearly distinct already in phase p1, and this difference increases in p2 and p3 (*Ω̃*_warm_ = 9.8 s, 15.5 s, 15.8 s and *Ω̃*_cold_ = 3.7 s, 2.8 s, 2.8 s for p1, p2, p3, respectively; Fig. 4*E*). In particular, short warm-water excursions (0.1 s < *Ω* < 10 s) and long cold-water excursions (10 s < *Ω* < 100 s) became less frequent over time. A similar pattern is observed in TR2 and TR4 (Supplementary Fig. 16B, D).

### Fish swimming speed increases during short-term exposure to cold water

In addition to displaying a thermotactic behavior in modulating their turning events upon experiencing temperature gradients, fish further reduced cold-water exposure through a thermokinetic behavior, by increasing their swimming speed when in cold water. For *T*_bottom_ ≤ 6 °C (TR3 and TR4), the swimming speed normalized by the mean speed of each fish during the entire treatment phase, *v̅̅*_n,t_ (*Materials and Methods*), was significantly greater (44% in TR3; 33% in TR4) when fish were located below compared to above the thermal interface (*t*-test, *p* < 0.001 and *p* < 0.05, respectively, Fig. 5*A,* Supplementary Tables 14–17). For TR2 (*T*_bottom_ = 8 °C), a similar increase (22%) was observed, but was not statistically significant due to outliers (likely cold-shocked fish) in one replicate (Fig. 5*A*). In contrast, in TR1 the swimming speed was similar during cold-water (*T*_bottom_ = 10 °C) and warm-water (*T*_top_ = 12 °C) excursions (Fig. 5*A*).

**Fig. 5.**
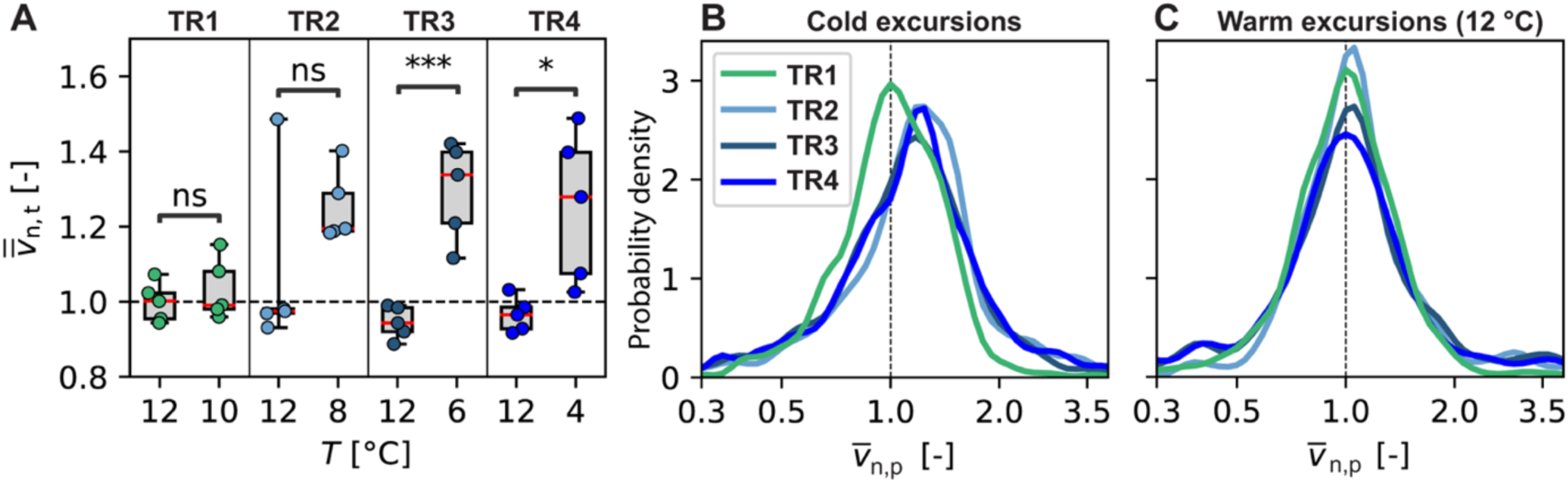
Fish swim more rapidly in the cold. (*A*) Mean swimming speed when above (in warmer water) and below (in colder water) the thermal interface. Speed was standardized by the average swimming speed of each individual during the treatment phases (p1–p3). Data was aggregated at the replicate level; hence *v̅̅*_n,t_ represents the average response of the fish group (four fish) within one replicate. Symbols represent the results of pairwise *t*-tests (ns, *p* > 0.05; *, 0.01 < *p* ≤ 0.05; ***, 0.0001 < *p* ≤ 0.001, see also Supplementary Table 16). Whiskers extent to the full range of observations and red lined indicate the median. The equivalent data for warm treatments are shown in Supplementary Fig. 22 and absolute swimming speeds are shown in Supplementary Figs. 20 and 21. (*B* and *C*) Probability density functions of averaged, swimming speed during cold-water excursions (*B*) and warm-water excursions (*C*) in cold treatments (TR1-TR4). Speed was standardized by the average swimming speed of each individual within each phase and *v̅*_n,p_ represents the average of this quantity over the duration of each individual excursion. The total numbers of cold-water excursions are TR1, *n* = 889; TR2, *n* = 389; TR3, *n* = 611; and TR4, *n* = 452; with identical numbers of warm-water excursions for each treatment. A kernel density estimator was fitted to the normalized histogram after log10-transforming the data. Equivalent figures for warm treatments are shown in Supplementary Fig. 23. Excursion durations were scattered against average normed swimming speed (see Supplementary Fig. 24).

Swimming speed increased over time (from p1 to p3) across all treatments (Supplementary Fig. 20). To account for this temporal trend in the analysis cold-water excursions, we further normalized the swimming speed by the average speed of that fish during the corresponding phase, *v*_n,p_. To give each excursion the same weight, independently of its duration, we further averaged that quantity for each excursion, *v̅*_n,p_. Notably, the distribution of average swimming speeds during cold-water excursions in cold treatments with *T*_bottom_ ≤ 8 °C (TR2-TR4) is clearly skewed toward values larger than *v̅*_n,p_ = 1. This indicates that the swimming speed was above average during the majority of cold-water excursions (Fig. 5*B*), indicating the presence of a thermokinetic response. In contrast, for TR1 (*T*_bottom_ = 10 °C) the speed during cold-water excursions peaks at *v̅*_n,p_ = 1 (Fig. 5*B*), similar to the warm-water excursions across all cold-water treatments (Fig. 5*C*), indicating the absence of a thermokinetic response in these cases. This observation was further supported by more frequently detected high absolute swimming speeds (> 20 cm/s) during cold-water occupation in TR2-TR4 (Supplementary Fig. 21). These findings demonstrate that fish exhibit an increase in swimming speed during excursions in cold water (*T*_bottom_ ≤ 8 °C), indicative of a rapid and transient thermokinetic response, on a timescale of seconds, to cold-water exposure.

## Discussion

Using a novel experimental design, we showed that trout parr acclimated to 12 °C display within minutes a strong avoidance of incoming cold water (*T*_bottom_ = 8, 6, 4 °C) through a combination of thermotactic and thermokinetic behavior, whereas they do not avoid incoming warm water (*T*_top_ = 14, 16, 18, 20 °C). Other fish species have been shown to avoid unfavorable thermal conditions, though often over timescales of hours to days^34–36^. For example, vertical relocation over timescales of hours has been observed for marine sablefish, *Anoplopoma fimbria*, acclimated to 12 °C and exposed to 2–8 °C ^37^.

Our observations revealed that the avoidance of cold water was mediated by two distinct behavioral responses: fish biased their vertical turns, which enabled them to redirect their swimming out of the colder water, and increased their swimming speed, which allowed them to reduce the time spent in the colder water. Cold-water excursions were of shorter duration and shallower the lower the temperature of the cold water was. Larval zebrafish have been found to exhibit thermotactic behavior similar in some respects to that of trout in our experiments, biasing both turn magnitude and turn direction to swim toward warmer water (40). However, in contrast to our observation of increased swimming speeds in colder water, swimming activity of larval zebrafish decreased in cold water^14^. Furthermore, to guide their movement zebrafish have recently been found to integrate temperature information over a time window of 0.4 s preceding the next swimming bout^12^. The duration of cold-water excursions of trout in our experiments were of a similar order of magnitude (0.1 s < *Ω* < 10 s), suggesting that thermal sensing in trout may occur over similar timescales to zebrafish.

Surprisingly, no superficial cold receptor has yet been identified in trout skin^38^ or in fish more broadly^5^, and recent studies suggest that fish may instead respond to temporal changes of their internal body temperature^39^. To investigate how body temperature might trigger behavioral responses, we used a heat transfer model for fish^22^ to estimate the decrease in internal body temperature *T*_body_ over the duration of cold-water excursions (see Methods and Supplementary Fig. 29 *AD*). We found that that the median body temperature *T*_body_ during cold-water excursions decreased by only 0.2 °C in TR1 up to 0.5 °C in TR4 (Supplementary Fig. 25). This suggests that the avoidance response could have been triggered by a small decrease in internal body temperature, in the order of 0.2–0.5 °C. Further, this analysis indicates that by reducing the duration of cold-water excursions, fish maintained body temperatures within the range 11.5–12 °C, ultimately preventing body cooling and the need to acclimate physiologically to the colder temperature.

Our results suggest that the observed thermoregulation behavior in warm-water treatments is a strategy to gradually increase body temperature. When the temperature of the incoming water was higher than 12 °C (TR6–TR9), fish exhibited a bimodal distribution of depth occupancy that was distinctly different from the acclimation phase and the cold-water treatments. This bimodal distribution was not due to variability in the temperature preference among individual fish, but rather to individual fish periodically moving between warmer and colder regions (Supplementary Fig. 8B). Interestingly, fish gradually increased the occupancy of warmer water over time (Supplementary Fig. 26B). In contrast to the cold treatments, no avoidance response was observed at the thermal interface in warm treatments. We propose that this periodic relocation between warmer and colder regions is a behavioral strategy to slowly elevate body temperature, while avoiding rapid and drastic body warming to 14–20°C (TR6–TR9). It has indeed been shown theoretically that frequent transitions between cold- and warm-water regions that are each, per se, thermally suboptimal, could allow fish to maintain body temperatures between the available extremes while avoiding excessive heating or cooling^39^. Our observations resemble this mode of behavioral thermoregulation, in which fish progressively favor warmer regions within a heterothermal environment. However, additional experimental evidence is required to determine the mechanisms underlying this behavior.

We conclude from the behavior of fish when warmer water was available that their acute thermal preferendum exceeded 12 °C, departing from the acclimation temperature we had chosen based on the thermal preferendum for trout reported in literature^33^. Indeed, the thermal biology for any fish species is more complex than a single, static thermal preferendum: Many internal and external factors, such as hypoxia, satiation, time of day, and life stage^5^, can influence the temperature preference of fish. For example, the level of satiation can have an impact because when fish are well fed, their growth rate increases with body temperature as metabolic performance increases^40^. This modifies the preferred temperature, as observed in Bear Lake sculpin (*Cottus extensus*) that ascend into warmer water after feeding to stimulate digestion and thereby achieve a three-fold higher growth rate^41^. In contrast, field studies with adult fish have observed movement from warm to cold water in summer^42,43^, allowing fish to lower their metabolic rate, likely in effort to conserve energy^2,44^. We propose that the behavior of trout parr upon exposure to warmer water in our experiments served to achieve a higher body temperature to ultimately increase growth rate, which is critical for this life stage^45,46^. Indeed, growth experiments on brown trout populations have shown that optimal growth temperatures can range between 15 and 19 °C, depending in the stream of origin^46^.

The behavioral thermoregulation exhibited by trout in our experiments also included a strong thermokinesis response. Fish rapidly increased swimming speed while in colder water, on a timescale of seconds during individual cold-water excursions (in cold treatments TR2 -TR4). This contrasts with the established paradigm for thermokinesis in fish. For a variety of species including trout^47^, swimming speed was found to decrease with decreasing water temperature. This is thought to be a physiological effect, as a lower temperature results in a lower metabolic rate and locomotory capacity due to decreased contractile rates in the red and white swimming muscles and the heart^48^. This type of thermokinetic response cannot serve to avoid cold water.

Our results instead suggest that trout parr employ a very different form of thermokinesis. Our analysis showed that increasing swimming speed during cold-water excursions allowed fish to shorten the exposure to cold water. This conclusion is supported by our model predictions that body temperature during cold-water excursions remained above 11.5 °C. Anecdotal observations from our experiments suggest that prolonged exposure to cold water (in the order of minutes) can nonetheless reduce swimming speed. Two individual fish in TR2 (*T*_bottom_ = 8 °C) appear to have become trapped within the cold water (Supplementary Fig. 26A) and displayed reduced swimming speeds, in line with previously described deactivation upon cold-water exposure^49,50^.

Taken together, these observations suggest that the thermokinetic response to cold water is timescale dependent. If the exposure to cold water is short (seconds), fish accelerate to escape the cold. If exposure is longer (minutes), however, fish slow down, likely due to a reduction in metabolic rate. Importantly, this timescale dependence implies that long-term (hours) measurements of behavior at constant water temperatures cannot be used to model fish movement response to thermal gradients occurring over shorter timescales. Our findings may help explain why recent modeling of fish behaviour in heterothermal environments could not reproduce the observed avoidance of cold water in stationary thermal gradients^14^.

Despite the avoidance response to cold water, fish engaged in repeated cold-water excursions, potentially reflecting a behavioral strategy to map the thermal environment. This pattern may also reflect an inherent tendency to occupy the lower part of the tank, as observed during homogeneous temperature of 12 °C during the acclimation phase. However, over time the frequency of longer cold-excursions (>10 s) diminished, reflecting a modulation of this exploratory behavior. While most studies have associated thermoregulation with sustained relocation from colder to warmer water or vice versa^51^, it has been suggested that short excursions into water of a different temperature might be a thermoregulatory behavior that allows fish to exploit resources in suboptimal thermal regions within a heterothermal environment^39^. For example, by keeping excursions into cold water short, hammerhead sharks can access deep-sea foraging habitats without critically cooling their body temperature^52^.

Our findings show that fish were able to perform upward turns while still located above the thermal interface and that is, before actually sampling the cold water below the interface. In fact, our simulation of the vertical extent of the thermal gradient revealed that a substantial fraction of upward turns occurred before fish encountered the gradient itself — that is, prior to any sensory detection of the temperature change (Supplementary Fig. 39). This finding may be evidence of associative learning, whereby fish used information on the presence of colder water at depth obtained at prior times. While the current data do not provide conclusive evidence in this regard, they prompt the possibility that, rather than responding solely to immediate thermal cues, fish use spatial memory or associative learning to anticipate the location of colder water based on prior experience. Indeed, fish are able to perform associative learning based on non-visual cues^53^, create mental maps of their surroundings^54^ and retain memory for hours^55^, days^56^ and months^57,58^.

Determining the mechanisms underlying fish behavior in heterothermal environments is challenging, because thermokinesis and thermotaxis may co-exist and play out across different spatio-temporal scales. In rivers and lakes, heterothermic environments form naturally and persist over a variety of spatial and temporal scales^59^. Anthropogenic warm- and cold-water releases expose fish to rapid (minutes) thermal changes, known as thermopeaking^21^, which can affect river sections hundreds of kilometers downstream^60^. Here, we mimicked such rapid thermal changes for environmentally relevant thermal contrasts of 2–8 °C using a lock-release setup. We have demonstrated that trout use a combination of thermotaxis (modulation of turning behavior) and thermokinesis (modulation of swimming speed) to avoid cold water, while occasionally still exploring it through brief cold-water excursions. In contrast, we found that for incoming warm water, fish explore both warm- and cold-water regions and do not display any thermotactic avoidance.

While limited to laboratory conditions, these observations provide a framework for interpreting the thermal behavior of fish in nature. For example, a recent study observed a significant increase in downstream displacement for early life stages of a cypriniform fish in response to cold-water thermopeaking^61^. The authors hypothesized that this response was caused by reduced swimming capacity and cold shock. Our findings suggest that the observed downstream displacement could alternatively have been caused by an avoidance response at the interface with cold water. However, a generalization of our observations to horizontally oriented thermal gradients remains elusive. Our results are inherently tied to the vertical stratification created in our experiments. As warm water was always positioned above and cold water below, we could not control for the effect of vertical position (i.e., we could not do cold over warm layer experiments). This limits our ability to directly compare our findings to those obtained from horizontally oriented thermal gradients. On the other hand, the case we addressed is of direct environmental relevance, as natural waters often experience vertical thermal stratification.

By capturing the multifaceted response of fish to thermal change, our results may help to predict the larger-scale distribution of species and their response to unnatural disturbances, and thus inform environmental conservation policies. Our study also highlights the importance of deepening our understanding of the fundamental behavioral mechanisms of fish to alterations in their environment, particularly anthropogenic ones. Ultimately, the large-scale distribution of an organism results from the combined effects of responses to countless stimuli on the individual scale. Our experimental setup provides a platform to investigate these stimuli through a reductionist approach, and a blueprint for how to investigate responses to other stimuli such as chemicals or salinity for diverse aquatic organisms.

## Methods

### Experimental animals

A total of 180 hatchery-sourced trout (standard length 3.6 ± 0.5 cm) were used for experimentation and each individual was used in a single experiment only. Fish were acclimatized to the housing facility (*T*_water_ = 12 °C) for a minimum of 24 h prior to experimentation and kept for a maximum of 5 days.

Fish originated from wild mature trout, artificially spawned in a cantonal hatchery (Flüelen, canton Uri, Switzerland) in December 2020. Fertilized eggs were transported (35 min by car) in 200 mL plastic containers to the Swiss Federal Institute of Aquatic Science and Technology (Eawag, Kastanienbaum), where they were transferred to temperature-controlled incubators. The water temperature during transport and incubation was maintained at 3 ± 0.5 °C. The eggs were incubated for three months until hatching. After hatching, the water temperature in the incubators was incrementally increased to 5 ± 0.5 °C to promote growth of the alevins. After yolk-sac absorption, trout were fed with half a teaspoon of *Artemia* and chironomid larvae per 50 fish once per day for one month and then placed in a covered outdoor housing tank, with water temperature ranging from 8 °C to 11 °C. All incubators and housing tanks were fed by lake water (Lake Lucerne) in a flow-through system.

When fish had developed to the parr stage^62^, they were transported (65 min by car) in an aerated, temperature-controlled (11–12° C) tank (60 L) to ETH Zurich (ETHZ), where they were acclimatized to the housing temperature of 12 ± 0.3 °C. Flow-through tap water continuously replenished the housing tank, water quality parameters (pH = 8–9, NH_4_ < 0.04 ppm, NO_3_ < 1 ppm, NO_2_ < 0.04 ppm) were monitored twice per day, and the fish were fed in the evenings with frozen bloodworms (1.5% of body weight). Experimental procedures and husbandry facility were approved by the Zurich veterinary office (licence no. 30997; housing facility no. 182). All fish were euthanized post-experimentation.

### Experimental setup

Experiments were conducted during daytime (09:00–17:00) in July 2021 at ETH Zurich. We performed experiments using an acrylic tank (dimensions 200 cm × 20 cm × 30 cm, length × width × height; Fig. 1*A*), with a removable opaque vertical gate (polymethyl methacrylate, PMMA, white 60% transparency) separating the tank into two compartments of equal dimensions (Fig. 1*A* and Supplementary Fig. 1). Three temperature probes (resistance thermoelement, type K-3895, ®Adafruit) were installed at three depths (4.5 cm, 14 cm, and 23.5 cm) in each compartment of the tank. A dissolved oxygen (DO) probe (WTW CellOx 325, galvanic, membrane based) was placed on each side wall of the tank (one at a depth of 4.5 cm and one at 23.5 cm, Fig. 1*A*). Probes were synchronized and provided continuous measurements at a sampling rate of 0.2 Hz (Supplementary Figs 27 and 28)). At the beginning of an experiment, each compartment of the tank was filled with water to a depth of 28 cm (56 L per side). The initial water temperature on the left side was always 12 °C; water temperature on the right side varied according to treatment. Water on the right side was dyed blue by the addition of 16.66 mg pre-dissolved Patent Blue V sodium salt (C_27_H_31_N_2_NaO_7_S_2_, Sigma-Aldrich; final concentration in half tank 0.29 mg/L) to provide a visual contrast and allow visualization of the thermal interface during the experiment (Supplementary Fig. 2). Prior to the experiment, the temperature on each side was adjusted by adding ice or stirring with a heater until it reached the desired temperature homogeneously across depth. Throughout the experiments, the DO concentration remained within the range 7–10 mg/L. The tank was sheathed with white diffusive material (PMMA, white 60% transparency) and homogeneously illuminated with white LED light (4000 Kelvin) from all sides except the top and front of the tank. This homogeneous illumination and tank sheathing was necessary to provide sufficient contrast for image-based tracking of fish.

### Experimental procedure

Each experiment consisted of four fish exposed to a temperature treatment. In total, we conducted five experimental replicates per treatment. Treatments were defined based on the initial water temperature in the right compartment of the tank (values of 4–20 °C, in 2 °C increments; Supplementary Table 1). Therefore, for all treatments, fish were initially located and acclimatized to 12 °C (the *thermal preferendum* of adult trout reported in the literature^33^), and exposed to a thermal interface towards warmer water on the top or colder water on the bottom. All water temperatures were within the thermal tolerance range of the species (0–25 °C ^63^) and specifically within the thermal range in which trout show positive somatic growth rates (3.6–19.5 °C in ^45^; approximately 5–23 °C in ^46^). After 15 min acclimation to 12 °C (Fig. 1*A*), the central gate was removed, triggering gravity currents^64^ caused by the density difference resulting from the water temperature difference between the left and right compartments of the tank (Supplementary Fig. 2). During the treatment phase, a sharp thermal gradient (i.e., a thermal interface) spanned the entire width of the tank and gradually stabilized at the half-tank depth such that its temporal vertical movement reduced to ∼5 cm (Supplementary Fig. 3). The interface could be visualized throughout experiments due to the blue dye in the water that was initially well mixed into the right compartment of the tank (see Fig. 1 *A* and *B,* Movies 1–4, and Supplementary Fig. 4).

Each experiment consisted of a 15-min acclimation period followed by a 20-min treatment period (Fig. 1*D*). A group of four fish was released into the left compartment of the tank containing water at 12 °C (identical to the husbandry temperature). After 3 min, cameras were started, initiating the acclimation phase. After 15 min, the lock was gently removed by hand (∼3 s) exposing fish to the gravity current. For the next 20 min, fish were exposed to the heterothermal environment throughout the tank. After each experiment, fish were captured and transferred to a separate compartment of the husbandry facility. To capture temporal trends in the behavioral response, we compared four 6-min periods (acclimation, p1, p2, and p3, Fig. 1*D*). During the treatment period, fish could choose to occupy lower colder water or upper warmer water. The final 6 min of the acclimation phase provided a reference for the absence of thermal contrasts. Phase 1 (p1) started 2 min after the lock was removed and the thermal interface had already occupied the entire width of the tank (Fig. 1*D*). Phase 2 and phase 3 (p2, p3) were the consecutive 6-min periods, during which the vertical movement of the thermal interface line diminished (Supplementary Fig. 2).

### Gravity currents

A gravity current is the flow of one fluid within another caused by the density difference between the fluids^65^. In our setup, this density difference was caused by the different water temperatures which defined our treatments (Supplementary Table 1). The treatment phase of the experiment was initiated with the slow removal (∼3 s) of the vertical lock separating the two compartments of the tank, followed by initial gravity currents^66^ and the formation of a horizontal thermal interface (Supplementary Fig. 2). The symmetric warm and cold water currents traveled at similar speeds but in opposite directions^66^ (Movies 1 and 3). Water from the colder compartment traveled along the tank bottom and remained sharply separated from the warm water above it. For similar thermal contrasts, it has been shown that despite some mixing, the interface between the two counter-flowing layers is stable if the density ratio *ρ*_left_/*ρ*_right_ is close to unity^66^, which was the case for all our treatments (Supplementary Table 1). Once the interface was established, the temperature contrast would smear out due to thermal diffusion. We simulated numerically (in Python) the thermal diffusion for the strongest thermal contrast (d*T* = 8 °C) and found that over the duration of an experiment (20 min), an initially sharp thermal interface (gradient thickness = 0) evolves into a gradient of thickness of ∼4-6 cm (Supplementary Fig. 29C). The density ratio between colder and warmer fluid was close to unity, resulting in slowly creeping currents with maximum velocities of ∼0.8–1.6 cm/s during the initial 2 min after gate removal. To eliminate the effects of flow on fish behavior, this period was excluded from the data analysis. After 1–2 min, the currents impounded at the side walls and the thermal interface extended over the entire width of the tank. Vertical movement of the interface (∼0.1 cm/s) was much slower than fish swimming speeds (4–6 cm/s) (Supplementary Fig. 20, Movies 1–4) and over time, the thermal interface stabilized at the half depth of the tank (Supplementary Fig. 3).

### Imaging and tracking of fish

To allow simultaneous tracking of fish behavior and the position of the thermal interface, experiments were continually filmed using a video camera (GoPro Hero6) mounted at a distance of 1.8 m from the front of the tank, recording color images at a resolution of 9.2 pixel/cm. The camera setting ‘linear field of view’ was selected to eliminate fish-eye effect typically captured by the cameras wide-angle lens, without compromising image quality.

We analyzed 26.3 h of video recordings (45 experimental replicates) and obtained individual movement trajectories at a temporal resolution of 24 fps (Supplementary Figs. 30 and 31) and an average tracking efficiency of 99% across all analyzed videos (Supplementary Fig. 14; Movies 1 and 3). Each video was cropped using custom-made Python scripts, and the start of the treatment phase was set at the time when the vertical gate detached from the water surface. As the blue dye was not visible in the red channel of the RGB image (Supplementary Fig. 33C), fish tracking was performed on the red channel using the open source software TRex^67^. Light conditions and dye concentration were identical for all experiments, allowing the use of an optimized set of tracking parameters (Supplementary Table 2). Due to the low density of fish in the tank, occlusions (in which two fish temporarily merge into one object) rarely occurred; however, the automated tracking procedure was reviewed for all videos. In difficult cases (e.g., when one individual was occluded by another moving in the same direction), individual identities were manually assigned based on simultaneous recording from a camera positioned above the tank.

### Visualization and evolution of the thermal interface

The dye used to visualize the thermal interface, Patent Blue V, is a non-toxic dye^68^ that has previously been injected without side effects into the dorsal fin of experimental fish at concentrations of up to 300 mg/mL^69^. However, to confirm the absence of potential side effects of the dye on fish behavior, five control experiments to TR3 (*T*_bottom_ = 6 °C) were performed without the addition of the dye. In the comparison of results from these control experiments with TR3, no statistically significant difference in vertical occupancy was detected (Supplementary Fig. 34 and Fig. 35). We therefore conclude that the presence of the dye did not affect fish swimming behavior. Furthermore, because the dye concentration and resulting color intensity were consistent across all treatments, the dye did not act as a confounding variable in our statistical comparisons.

Locating the thermal interface was central to relating fish movements to their immediate temperature environment. For each time instance (frame), a line, representing the thermal interface, was derived from the visual signal of the dye. To do this, an in-house image analysis pipeline was developed in Python Opencv^70^. The pipeline consisted of background subtraction, thresholding, dilation, erosion and finally the abstraction of the interface line *y*_I_(*x*,*t*) (python functions: cv2.thresh, cv2. dilate, cv2. erode, cv2.Laplacian) (Supplementary Figs. 33 and 36). Our analysis assumes that over the duration of p1 + p2 + p3 = 18 min, the abstracted color boundary is representative of the location of the thermal contrast. We thus assume that the molecular diffusion driving heat exchange and molecular diffusion of the dye acted on similar time-scales.

### Characterization of the thermal gradient

Due to the combined effects of turbulence and thermal diffusion, the thermal gradient gradually weakens and expands over time. To quantify its vertical extent, we simulate the evolution of the gravity current over a 20-min period following its release. To account for transient mixing during the initial movement of the gravity current (within the first 2 min after removing the gate), we adopt a phenomenological approach to estimate the gradient thickness after the gravity current first hits the side wall of the tank. Then we predict the effect of thermal diffusion in a numerical simulation. This approach accounts for transient mixing in the initial, more turbulent phase and provides a realistic estimation of the subsequent evolution of the thermal gradient by diffusion.

First, we introduce two types of interface thickness: the enstrophy thickness (𝛿_!_) and the temperature thickness (𝛿*_k_*). From the thermal diffusivity (𝜅 = 1.4 · 10^-7^ 𝑚^2^/𝑠) and the momentum diffusivity (i.e., kinematic viscosity) of water (𝜈 = 10^-6^ 𝑚^2^/𝑠), we obtain the Prandtl number, 𝑃𝑟 = 𝜈/𝜅 = 7.14. In turbulence, the enstrophy thickness can be approximated^71^ by 𝛿*_v_*= 10𝜂, where [inline] is the Kolmogorov length scale and 𝜖 is the turbulent energy dissipation rate. Thus, the thickness of the thermal interface is given by [inline], which in our case amounts to 𝛿*_k_* = 3.7𝜂.

The Kolmogorov length can be estimated from large-scale balances as follows. The large-scale velocity can be estimated as [inline], where 𝑔 is the gravitational acceleration, 𝜌 is the mean water density, Δ𝜌 is the density difference, and ℎ, corresponds to half of the total water depth^66^. Note that this scaling assumes that the large-scale fluid motion is independent of viscosity. To verify this assumption, we measured the propagation speed of the gravity current 𝑢_-_ in our experimental videos for each cold-water treatment (TR1-TR4) and plotted it against the theoretical propagation velocity 𝑢, (Supplementary Fig. 29A). The good fit between the theoretical and observed values (𝑅^2^ = 0.96) confirms that the flow is indeed large-scale dominated. Therefore, the Kolmogorov length 𝜂 can be estimated by using the production-dissipation balance, 𝜖 = 𝐶𝑢_p_^3^/ℎ_0_, for each temperature contrast in the cold-water treatments (TR1-TR4), where 𝐶 is a non-dimensional constant of order unity that can vary among different flow conditions.

Assuming 𝐶 = 1 we calculated the turbulent energy dissipation rate and, subsequently, the Kolmogorov length for each cold-water treatment (Supplementary Fig. 29B). Across our treatments, we found a maximum estimated Kolmogorov length of 𝜂 ≈ 0.76 mm in TR1 (Fig. S29 B), indicating that turbulence caused the thermal interface thickness to reach 𝛿*_k_* = 3.7 𝜂 = 2.8 mm. To account for turbulence, we initialized thermal diffusion simulations by using this value of 𝛿*_k_* as the initial interface thickness and we ran a numerical simulation of thermal diffusion for t = 20 min. After 20 min the interface thickness reached a maximum of ∼4 cm in TR1 and ∼6 cm in TR4, depending on the thermal contrast Δ𝑇 (Supplementary Fig. 29C).

### Identification of time spent in colder and warmer regions

The interface line was used to differentiate whether fish were located in the warmer water above or the colder water below. To do this, we derived the vertical distance from the thermal interface *D*(*t*)

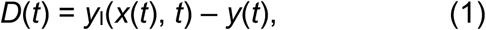

where *y*(*t*) is the vertical position of an individual fish and *y*_I_(*x*(*t*), *t*) is the vertical position of the interface line at the lateral position of the fish *x*(*t*) (Supplementary Fig. 37). Evaluating the sign of *D*(*t*) reveals whether a fish was located above or below the thermal interface (Fig. 2 *A* and *B*).

### Cold- and warm-water excursions

Evaluating the sign of *D*(*t*) allowed the individual fish trajectories to be partitioned into periods of cold- and warm-water occupation (i.e., excursions; Fig. 2*B*). We counted the number of cold- and warm-water excursions (*N_warm_* and *N_cold_*), which represents the number of times each fish crossed the thermal interface in one direction (*N*_crossings_), and for each excursion we quantified the duration *Ω*, and the maximum distance to the interface line (*D*_max_). Excursions can be interpreted as periods of self-inflicted exposure to the cold or warm water.

We derived the duration *Ω* of individual cold and warm water excursions as

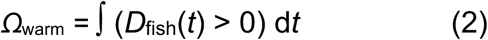

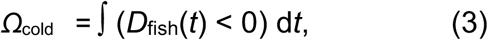

and the maximum excursion depth during cold excursions as

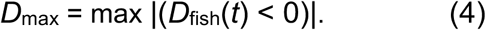

### Time spent in cold and warm regions

We calculated the total time spent above and below the interface as

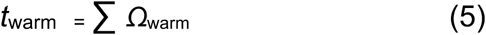

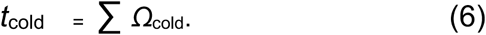

For cold treatments, the time *t*_warm_ represents the time an individual spent above the thermal interface (at 12 °C, which is equal to the acclimation and housing temperature), and *t*_cold_ the time spent in the respective treatment temperature (4 °C, 6 °C, 8 °C, or 10 °C). In the warm treatments, the time *t*_cold_ represents the time the individual spent at 12 °C, and *t*_warm_ the time spent in the respective treatment temperature (14 °C, 16 °C, 18 °C, or 20 °C).

### Body cooling during cold-water excursions

To estimate fish body cooling during cold-water excursions in TR1–TR4, we applied a heat transfer model and calculated the fish’s body temperature after each cold-water excursion^39^. Firstly, the heat transfer rate k for a fish with mass *m*_body_ = 4 g was estimated based on the parameters a = 2.267 and b = - 0.329 (Supplementary Table 4):

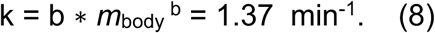

We then assumed an initial body temperature of *T*_b_ = 12 °C (corresponding to the fish residing above gradient for a prolonged period of time) and simulated body cooling during measured durations *Ω* of excursions into colder water with temperature *T*_a_. Thereby, the rate of change of fish body temperature is given by:

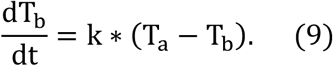

Thus, assuming the cold-water temperature to be a single, constant value (i.e., neglecting the effect of the thermal gradient), the fish body temperature after a cold-water excursion with duration *Ω* is:

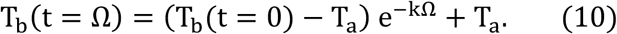

Based on the measured excursion durations in TR1–TR4 (Fig. 4 *D,E* and Supplementary Fig. S16 *A*–*D*), equation (10) allowed us to estimate the median body temperature (*T*_b_) change during cold-water excursions (Supplementary Fig. 25).

### Vertical turning points

Vertical turning points were obtained by fitting a fifth-order polynomial to the vertical position signal *Y*(*t*) of each trajectory within a moving window of 2.1 s (50 frames) (using the Python script scipy.signal.savgol_filter,^73^). This procedure smoothed the vertical position, reducing the noise produced by small-scale variation. We then detected upward turns as local minima and downward turns as local maxima in this smoothed signal (Supplementary Fig. 11). To focus on turning points that resulted in considerable vertical displacements, we eliminated upward- or downward-turn events if a subsequent turn occurred within a vertical distance of d*y* ≤ 3 cm (Supplementary Fig. 14A).

### Swimming speed

The normalized swimming speed for each individual was derived by dividing the instantaneous swimming speed by the average swimming speed of the individual during (i) the entire treatment period of 18 min (*v*_n,t_) or (ii) each phase separately (*v*_n,p_), as described within individual figure captions.

Upon visual inspection, the individual swimming trajectories seemed reasonably smooth (Supplementary Fig. 38A) however, unrealistic fluctuations appear in their first order derivatives (i.e., fish swimming speed) (Supplementary Fig. 38C and D). Before calculating swimming speeds, we therefore fitted a third-order polynomial to the trajectories obtained from ‘TRex’^67^, using a fitted to the *x*,*y* coordinates. For each time step, the polynomial was fitted to a temporal window of 21 measurements (1.4 s) following a procedure described by^74^ (Supplementary Text 3, Fig. 38 *B*-*E* and Movie 5). Further, data gaps (arising mainly from fish occlusions) were filled using temporal linear interpolation, and an upper velocity threshold of 100 cm/s (i.e., an unrealistically high speed of approx. 3× burst swimming) was used to discard unrealistically high values.

### Statistical analysis

The number of replicates used in this study reflects a balance between statistical rigor and the ethical imperative to minimize the use of animals in experimentation. Regarding statistical power, our design (five replicates with groups of four fish each) is consistent with similar studies and represents an adequate sample size.

To compare behavioral responses between treatments, all analyses considered the mean or median values for each group of four individuals within each replicate, accounting for dependency between individuals. Therefore, to compare the number (Fig. 3*C*) of cold-water excursions across cold-water treatments, one-way ANOVA was used (Supplementary Tables 18 and 19). Post-hoc pairwise *t*-tests were used to assess differences between treatments (Supplementary Tables 16, 17 and 20).

To determine whether the duration (Fig. 4*B*) and depth (Fig. 4*C*) of cold-water excursions followed a trend across treatments (i.e., decreased or increased with decreasing treatment temperature), we used the multivariate Mann-Kendall test^75^ (Supplementary Tables 5 and 6). We used pairwise *t*-tests to test whether the normalized swimming speeds differed during warm and cold-water occupation (Fig. 5*A*). Prior to performing statistical analyses, data distribution (normality) was assessed using the Shapiro-Wilk test (Supplementary Tables 14, 15 and 18). All statistical tests were conducted using Python packages.

## Data availability

The Python scripts used for the analyses are available through a public GitHub repository (https://github.com/naroberto/Thermal_paper/). The generated videos for fish-tracking can be accessed in an open access archive (https://doi.org/10.3929/ethz-b-000660506).

## Supporting information

Supplementary revised

## Acknowledgments

We thank Jukka Jokela, Armin Peter, Annamari Alitalo and Samia Bachmann for their support with the animal experiments and in obtention of the license for animal experimentation. We thank Werner Tresch and Kunio Takatsu for supplying and taking care of the experimental animals in Flüelen and at Eawag Kastanienbaum. We thank Thomas Meierhans, Michael Arnold, Ernst Bleiker, Ela Burmeister and Daniel Braun for providing workshop tools and laboratory resources that enabled construction of the experimental setup. We further thank Davide Vanzo, Oliver Selz, Jakob Brodersen, Steffen Schweizer, Francesco Carrara and Johannes Keegstra for helpful discussions.

---

Supplementary Figures

**Fig. 1:**
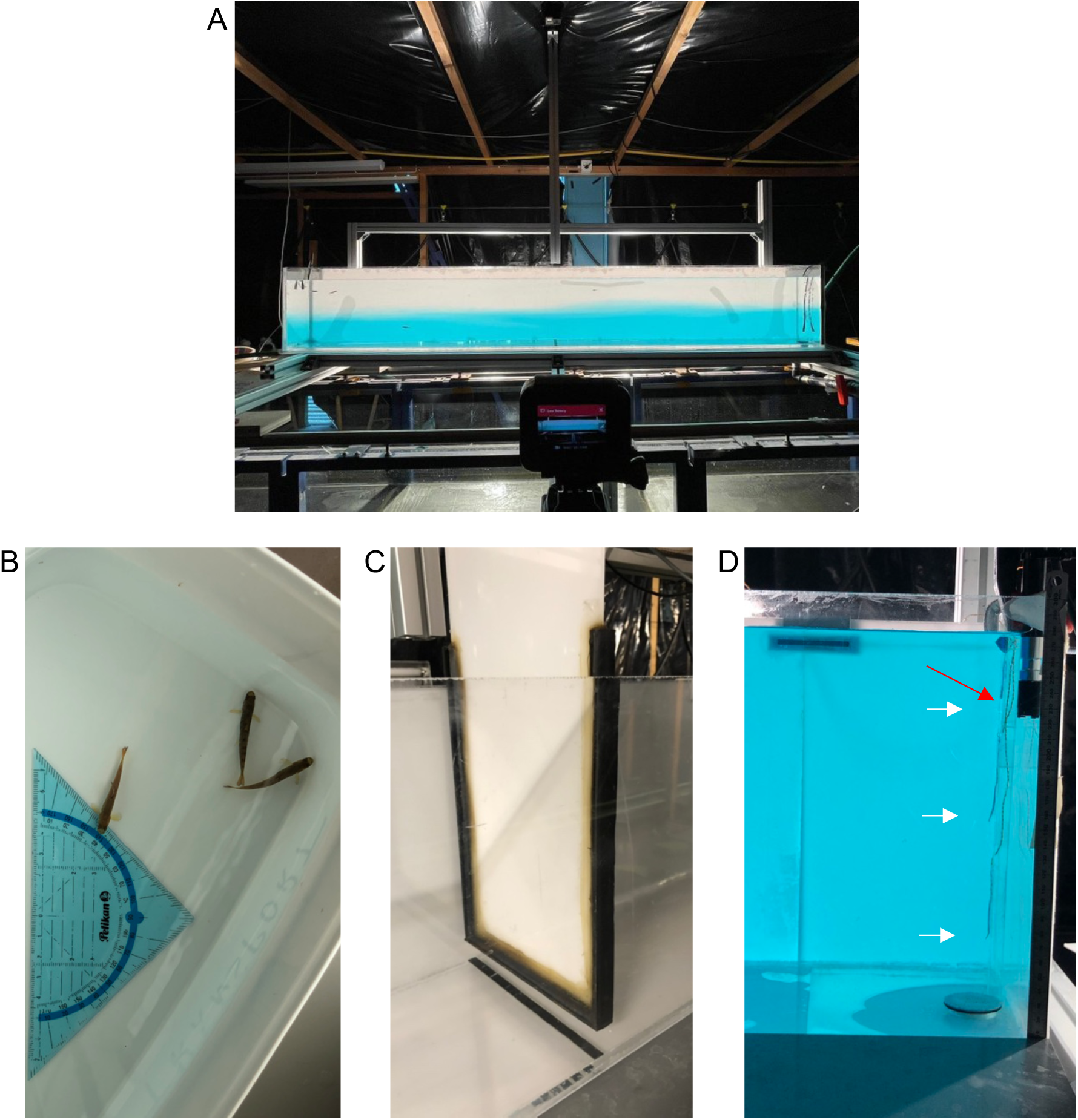
(*A*) Photograph showing a side-view of the experimental tank in phase 2 (*approximately 10 min* after removal of the central gate) with four individual fish. The camera used for imaging is visible in the foreground. The dark casing obstructs the entry of natural light. Experimental animals (*B*); central gate to partition the tank; (*C*) right side of the tank with 3 temperature probes (white arrows) and dissolved oxygen probe (red arrow) (*D*).

**Fig. 2:**
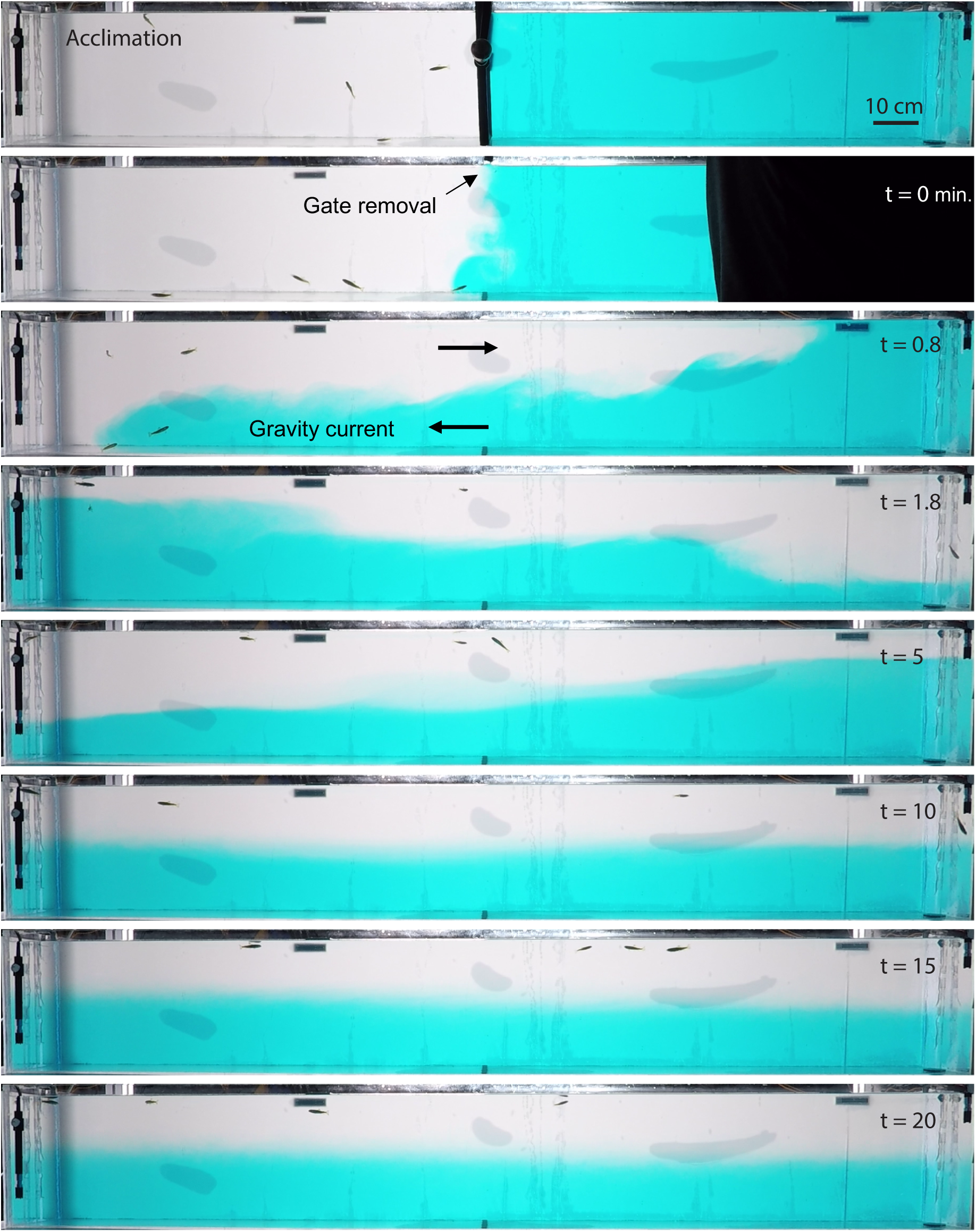
Side view of temporal progression of gravity current for cold treatment. During the acclimation phase (top panel) the water tank is partitioned into 12° C in the left compartment and 4°C (TR4) in the right compartment. After 15 minutes of acclimation the central gate is removed (*here at time:* t = 0) and the density difference drives a lock exchange flow with the transparent warm water flowing *slowly* in the upper half of the tank to the right and the dyed cold water flowing along the channel bottom towards the left (t = 0.8 minutes). After reaching the side walls of the tank, the directionality of the flow reverts and the current sloshes back and forth in the form of an internal wave (t = 1.8 – 5 minutes). A clear second and a third wave passage take place, while flow speed and wave amplitude gradually attenuate through dissipation. During these internal wave passages, the vertical height of the temperature interface remains a periodic function of time. A stationary vertical stratification (i.e. the water is practically still) is reached only after about t = 10 minutes. The start of the first treatment phase (p1) was defined as the time when the vertical gate was fully removed from the water (t = 0).

Temporal movement of the interface

**Fig. 3:**
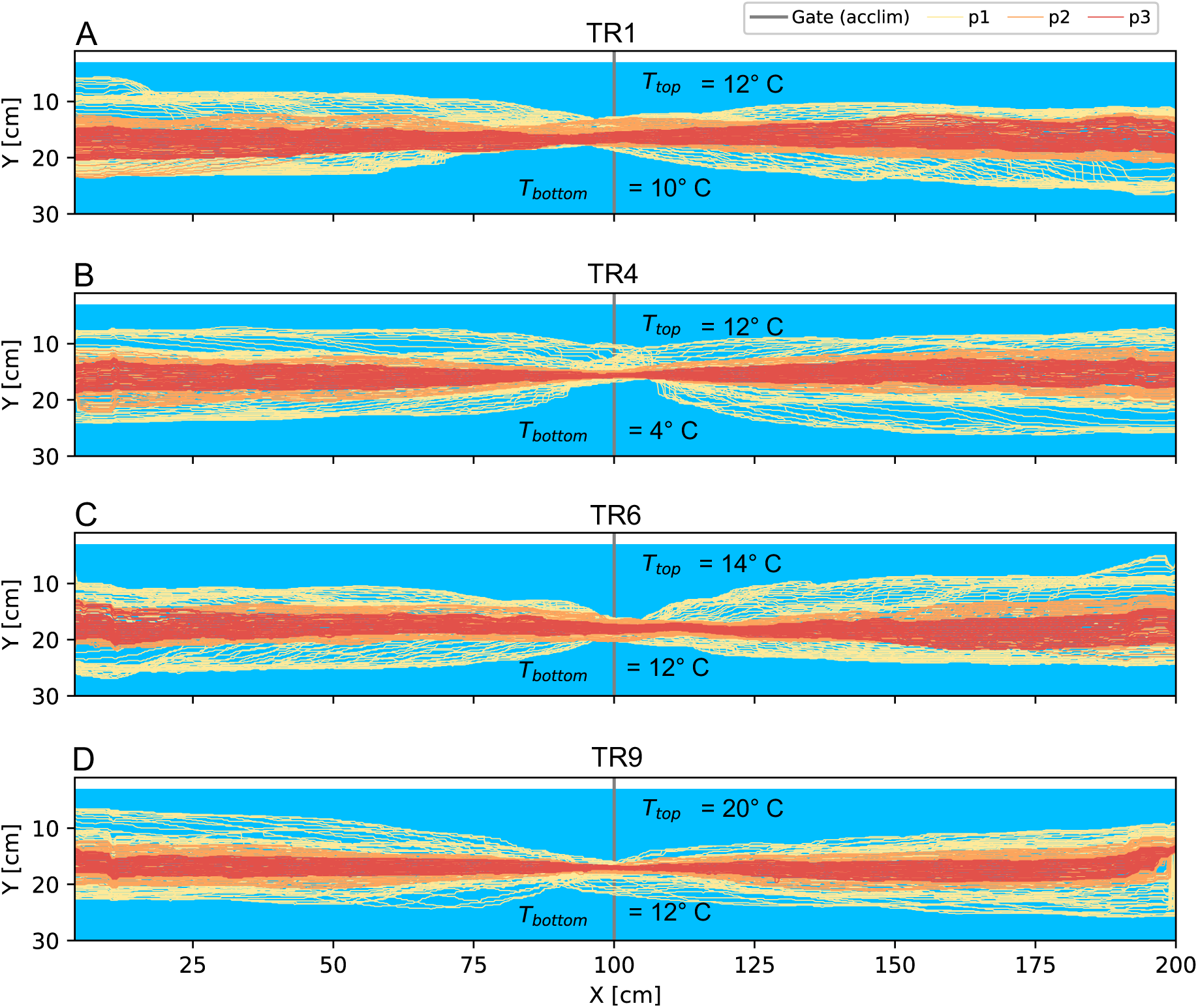
Transient movement of the thermal interface/line during experimental phases (p1, p2 and p3). Coloured lines depict time steps of 10 seconds. The water surface is located at *Y* = 4 cm. The instantaneous position of the thermal interface is shown for each experimental phase (acclim., p1, p2, and p3) at a reduced temporal resolution of 0.2 Hz (the temporal resolution in the experiments was 24 Hz). The interface was approximated as a line (X,Y) for each frame. During the acclimation period (acclim: grey dots) a sealed gate was located at X = 100 cm. The interface is coloured by the respective temporal phase of the experiment (see *A*).

**Fig. 4:**
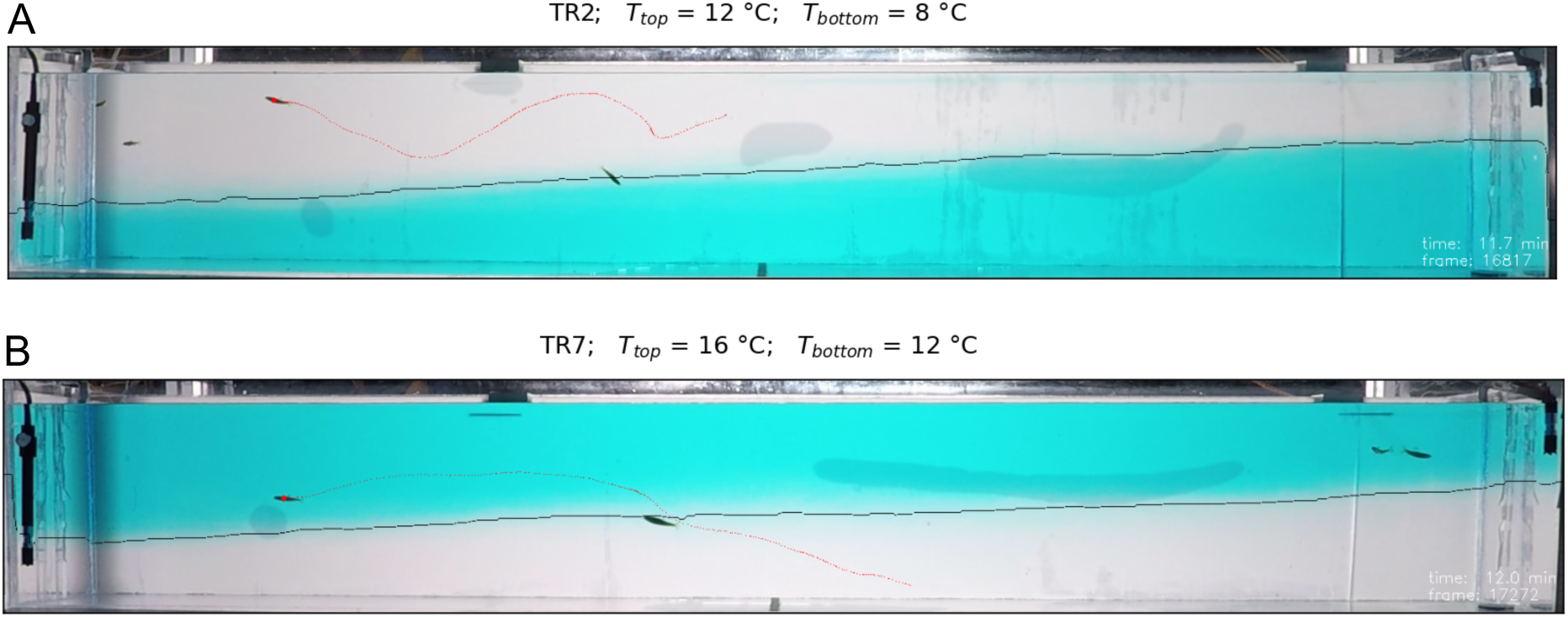
Side view of experimental setup for cold (*A*) and warm (*B*) treatment (TR2 and TR7). The thermal interface line (black) was derived using custom made image analysis pipeline.

Vertical occupation

**Fig. 5:**
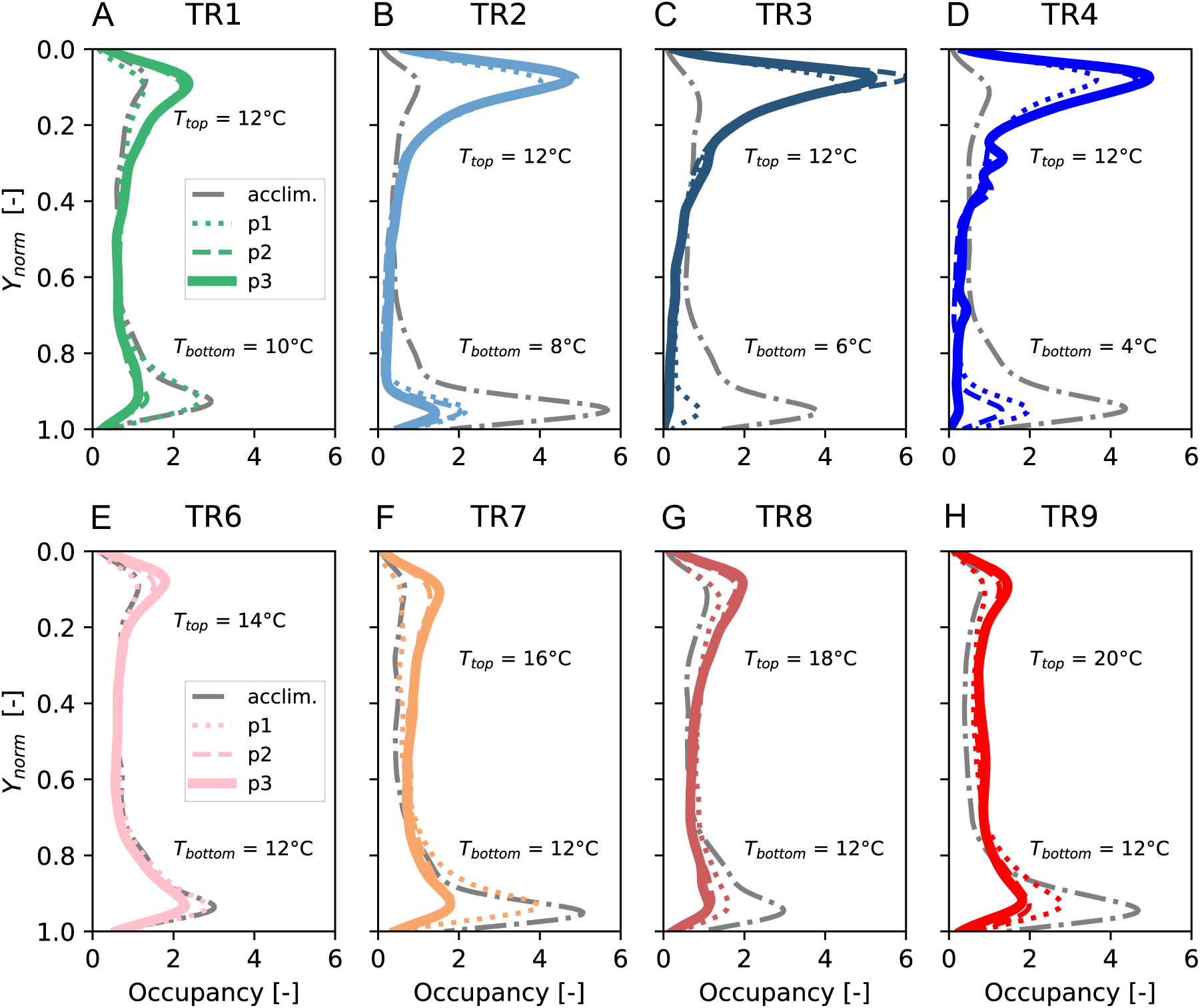
Probability density functions of fish depth for all cold-water (TR1–TR4, *A*–*D*) and all warm-water (TR6–TR9, *E*–*H*) treatments (*N* = 20 fish in each case), separated according to the four phases of analysis (acclimation, and the three experimental phases p1–p3). The data was aggregated at the treatment level; hence each line represents data from 20 fish, tested in groups of 4 individuals in 5 separate experiments. *T*_bottom_ and *T*_top_ indicate the water temperature below and above the thermal interface. The depth was normalized so that *Y*_norm_ = 0 is the water surface and *Y*_norm_ = 1 indicates the bottom of the tank.

Temporal progression of excursions (cold-treatments)

**Fig. 6:**
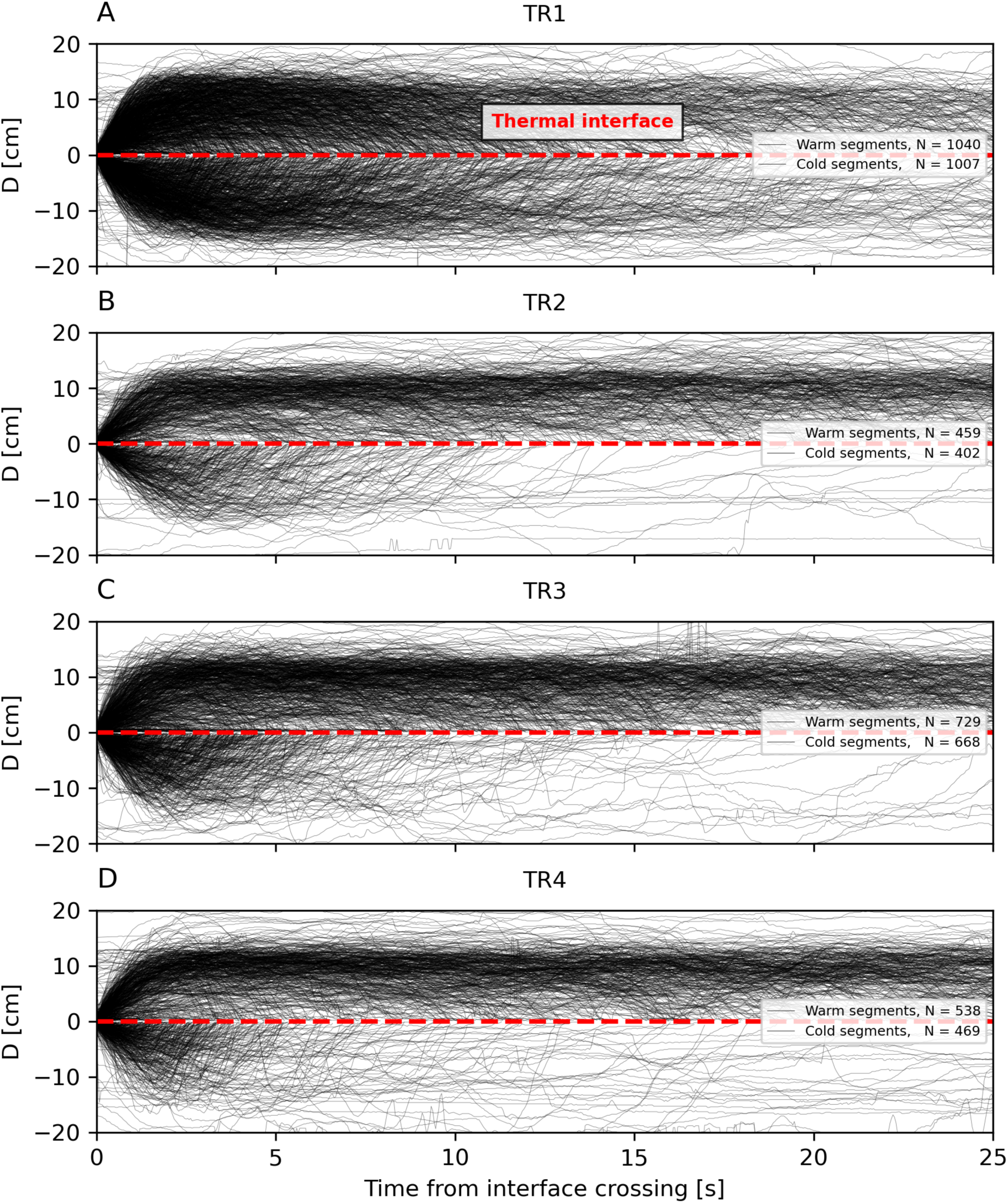
Cold treatments, TR1-TR4. Temporal evolution of the fish’s instantaneous vertical distance to the thermal interface D(t) within the first 25 seconds for each cold- and warm-water interval for all cold treatments. At t = 0 fish crosses the thermal interface line upwards (warm segment; upper half) or downwards (cold segment; lower half). The end of an interval occurs when the interface line was crossed again in the opposite direction.

Temporal progression of excursions (warm-treatments)

**Fig. 7:**
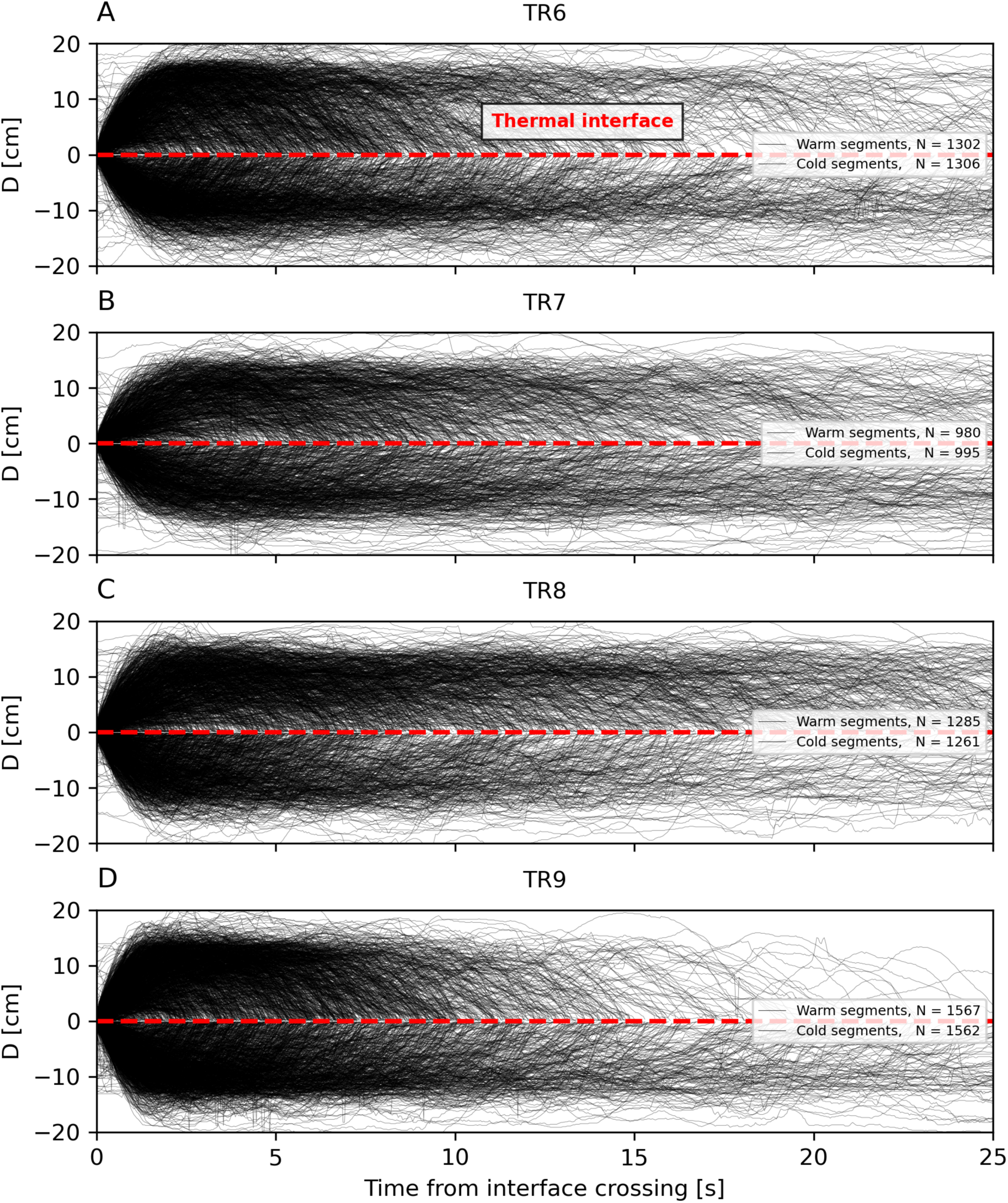
Warm treatments TR6-TR9. Temporal evolution of the fish’s instantaneous vertical distance to the thermal interface D(t) within the first 25 seconds for each cold- and warm-water interval for all warm treatments. At t = 0 fish crosses the thermal interface line upwards (upper half) or downwards (lower half). The end of an interval occurs when the interface line was crossed again in the opposite direction.

Time fraction in acclimation water

**Fig. 8:**
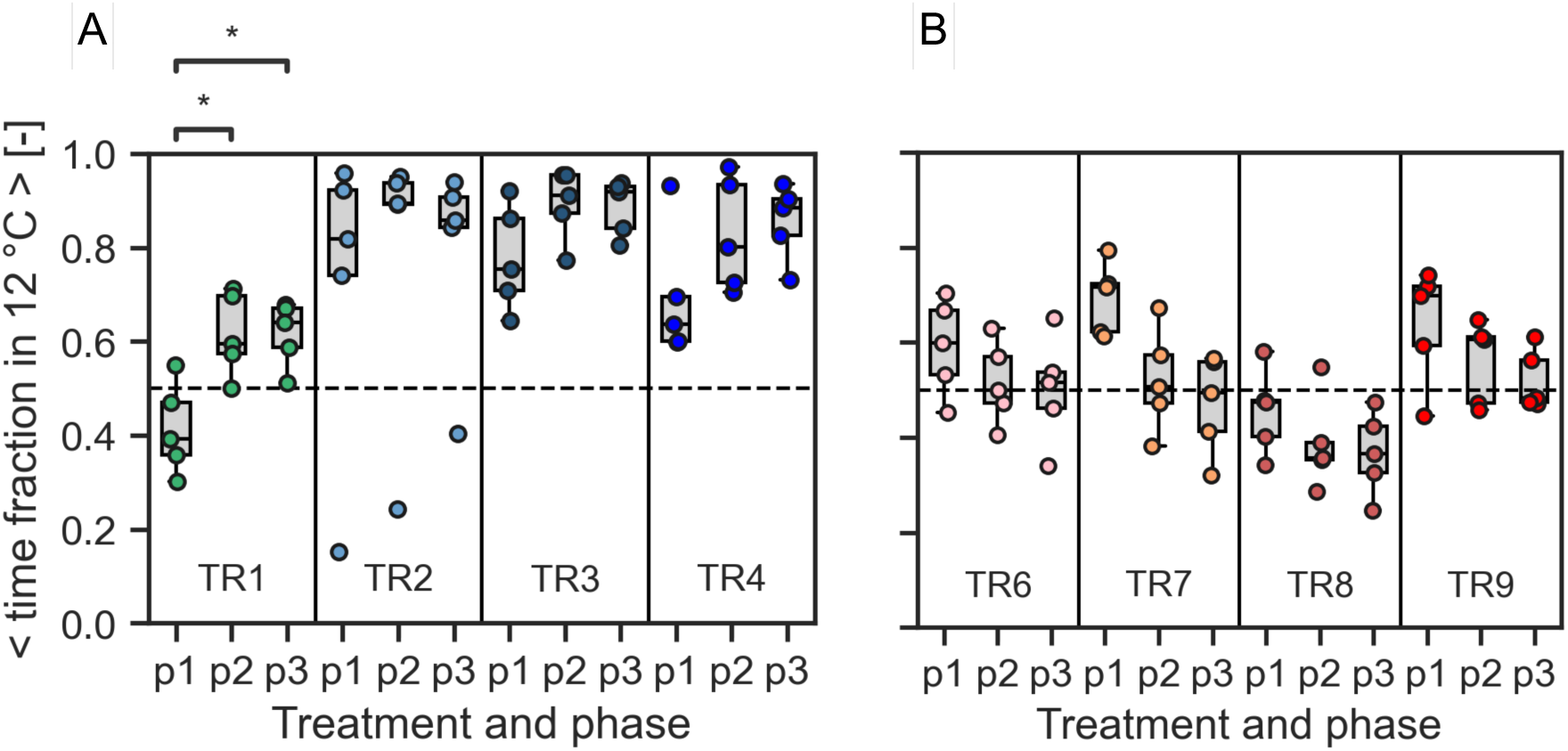
The average time fraction spent in 12 °C was calculated for each fish and phase (p1, p2 and p3) as *T_cold_* / 6 minutes for incoming cold water (TR1-TR4, *A*) and incoming warm water (TR6-TR9, *B*). Each dot represents the average of this quantity over four individuals that were tested simultaneously within one experimental replicate. The Friedman-test suggests that there are significant temporal differences within TR1 (p = 0.022). Significance levels are indicated above the plot (Dunn-test: * : 0.01 < p ≤ 0.05).

Vertical occupation of the tank

**Fig. 9:**
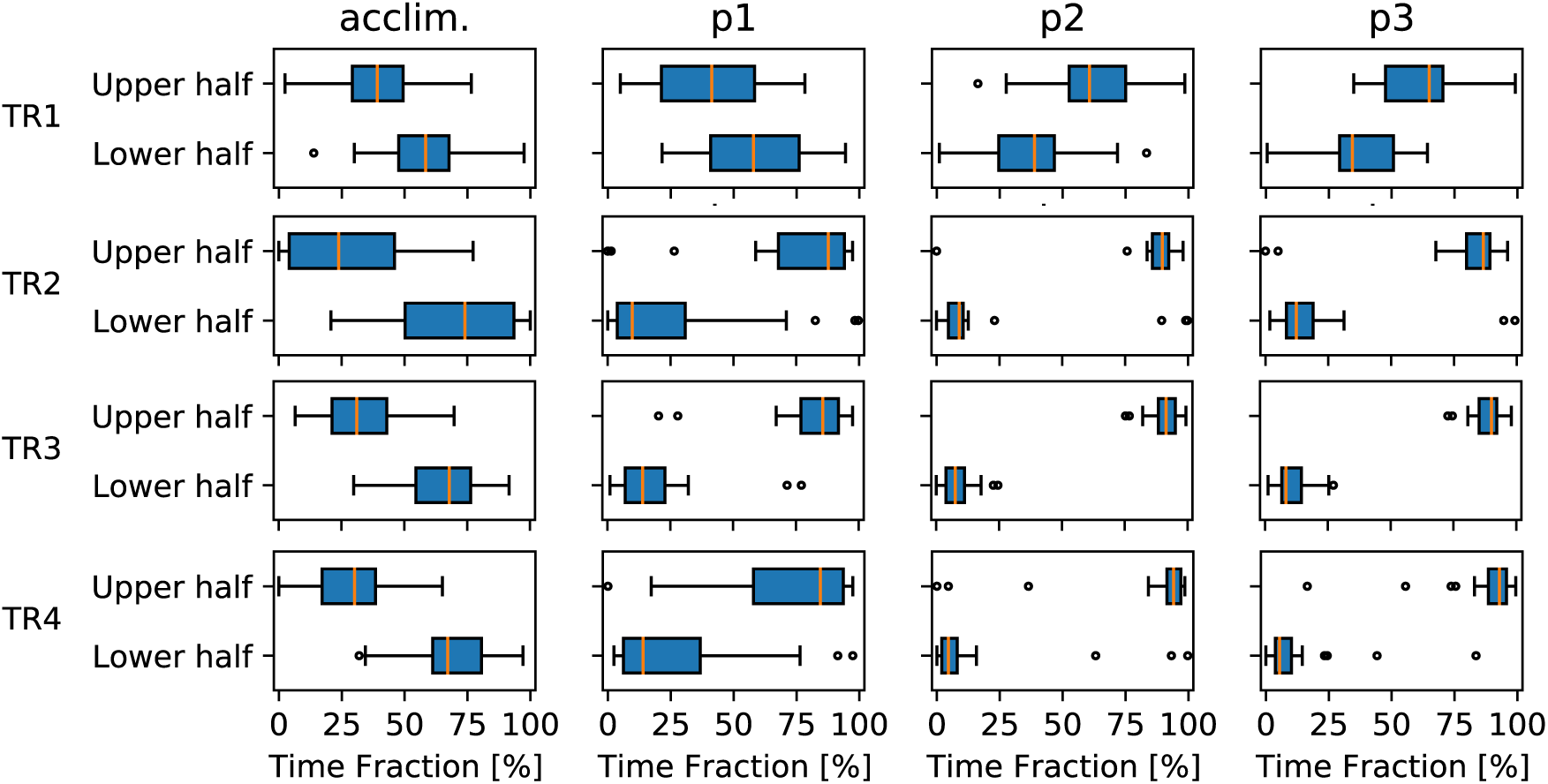
Cold treatments. Boxplots of time fraction spent in upper half (*Y*_norm_ = 0–0.5) and lower half (*Y*_norm_ = 0.5–1) of the tank across experimental phases (acclim., p1, p2 and p3) and treatments (TR1, TR2, TR3 and TR4).

**Fig. 10:**
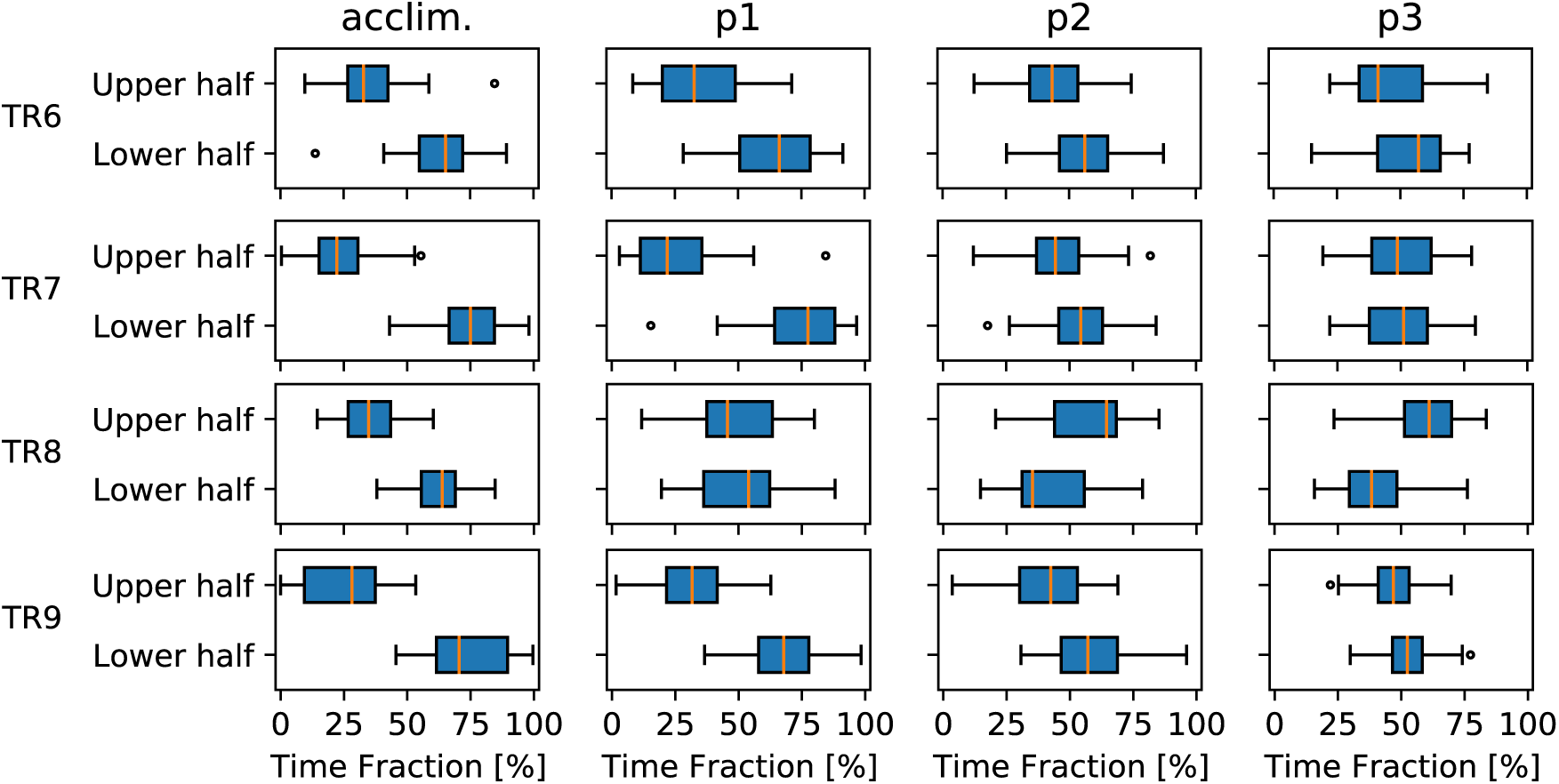
Warm treatments. Boxplots of time fraction spent in upper half (*Y*_norm_ = 0–0.5) and lower half (*Y*_norm_ = 0.5–1) of the tank across experimental phases (acclim., p1, p2 and p3) and treatments (TR6, TR7, TR8 and TR9).

Detection of vertical turning points

**Fig. 11:**
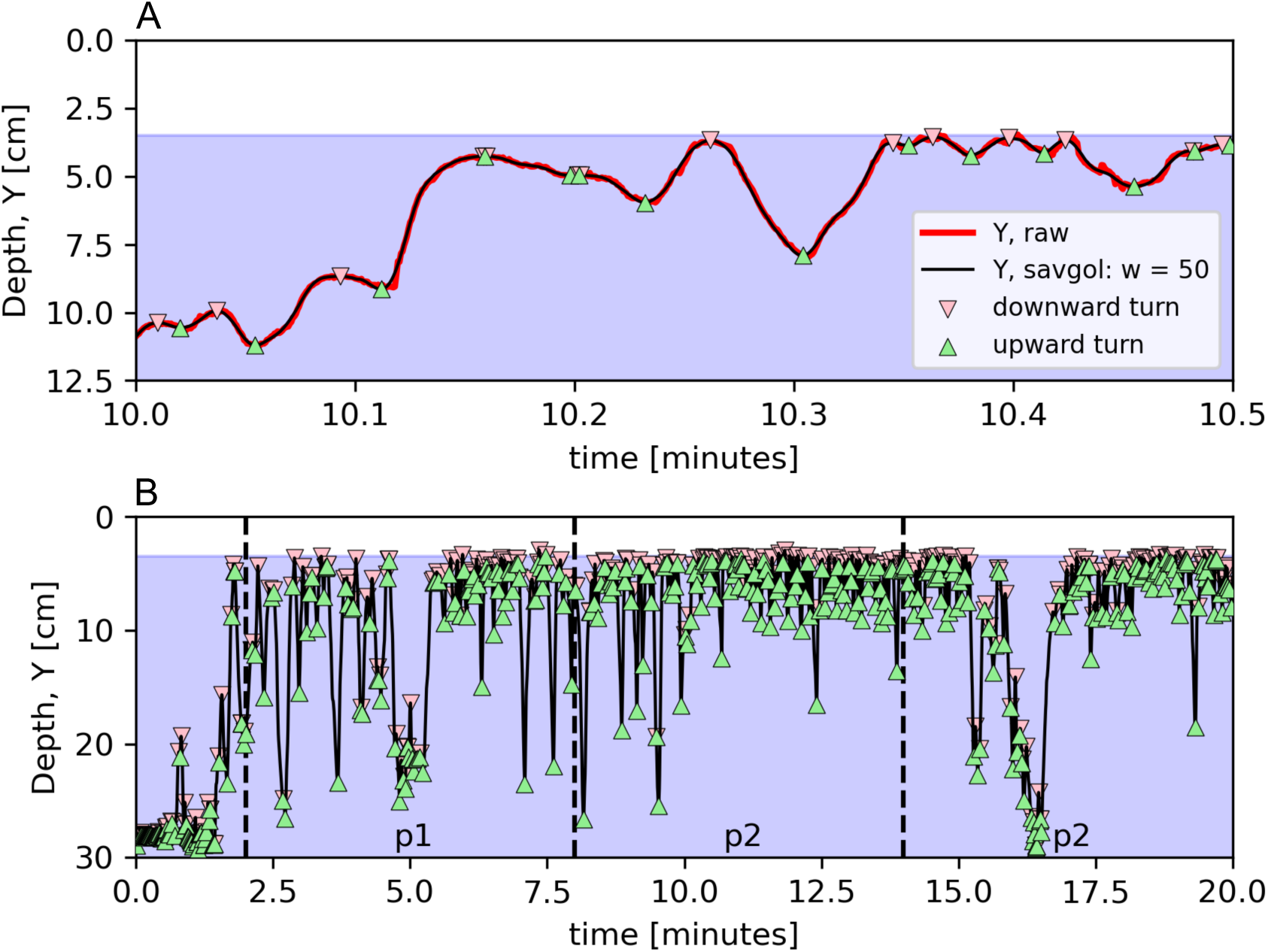
Derivation of vertical turning points. (*A*) To smoothen the fish’s vertical position signal (red line), a fifth order polynomial was fitted to a moving window of 50 consecutive time points (black line). This is a standard practice in particle tracking to filter out noise (e.g. Lüthi et al. JFM 2005). Therefore the SAVGOL function (python: scipy.signal.savgol_filter) was applied on the raw output (black line) from the tracking software. Upward turning (green triangles) were detected as sign change (from negative to positive) of the derivative of the smoothed signal. Downward turning (red triangles) were detected via sign change (from positive to negative). (*B*) Occurrence of turning points after gate removal (at t = 0) for an individual fish in cold treatment (TR3).

Large turns cold treatments (TR1 – TR4)

**Fig. 12:**
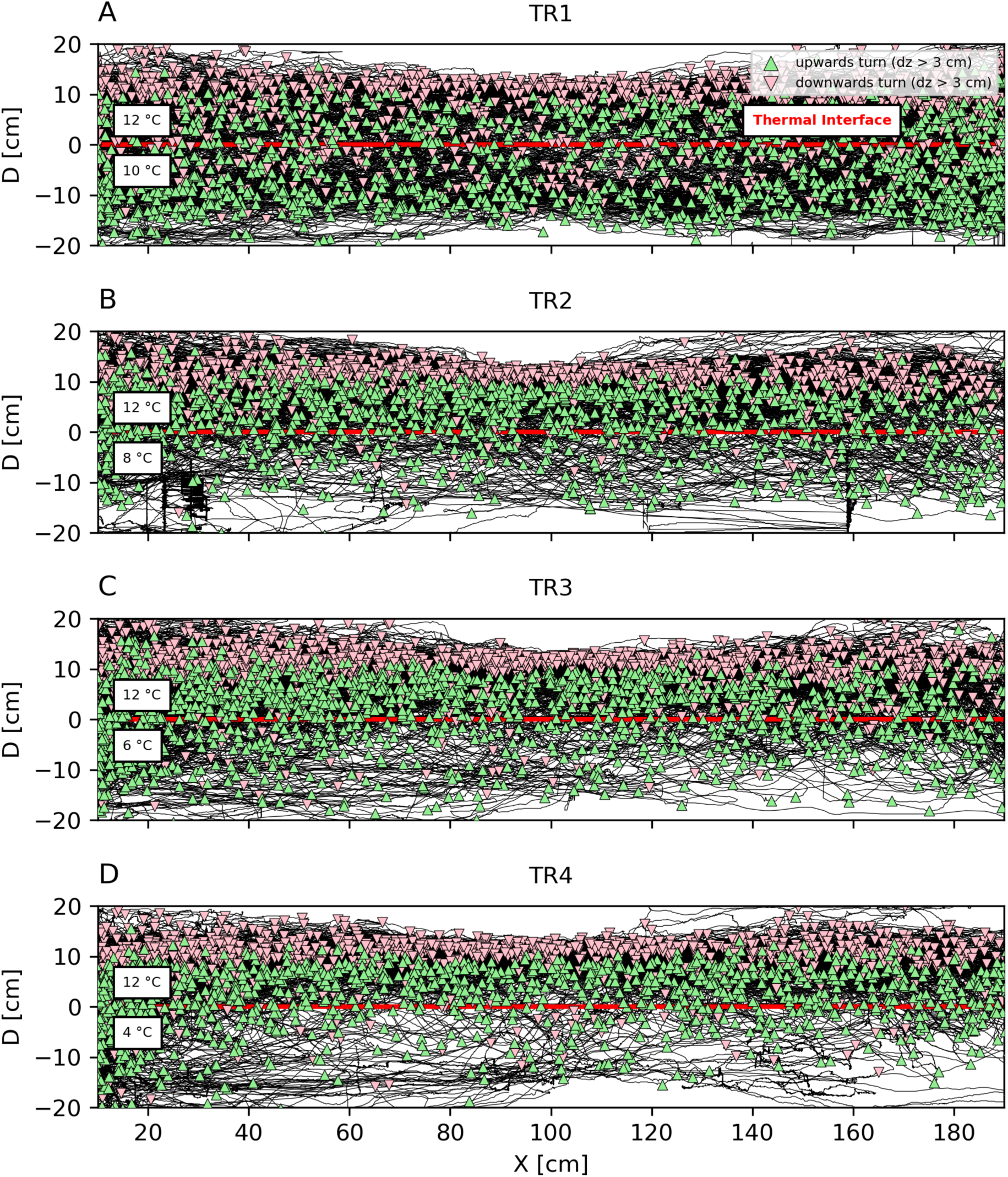
Cold treatments. Swimming trajectories (black lines) relative to the thermal interface (red line) for all fish across cold treatments (TR1 – TR4, *A*-*D*). The horizontal axis depicts the horizontal position in centimetres; the vertical axis depicts the instantaneous distance from the thermal interface. For D > 0 fish is located at a distance D [cm] above the interface and for D < 0 the fish is located below the temperature interface. Red and green triangles depict large downward and upward turns, respectively (see Supplementary Fig. 11 and Fig. 14 for derivation).

Large turns warm treatments (TR6 – TR9)

**Fig. 13:**
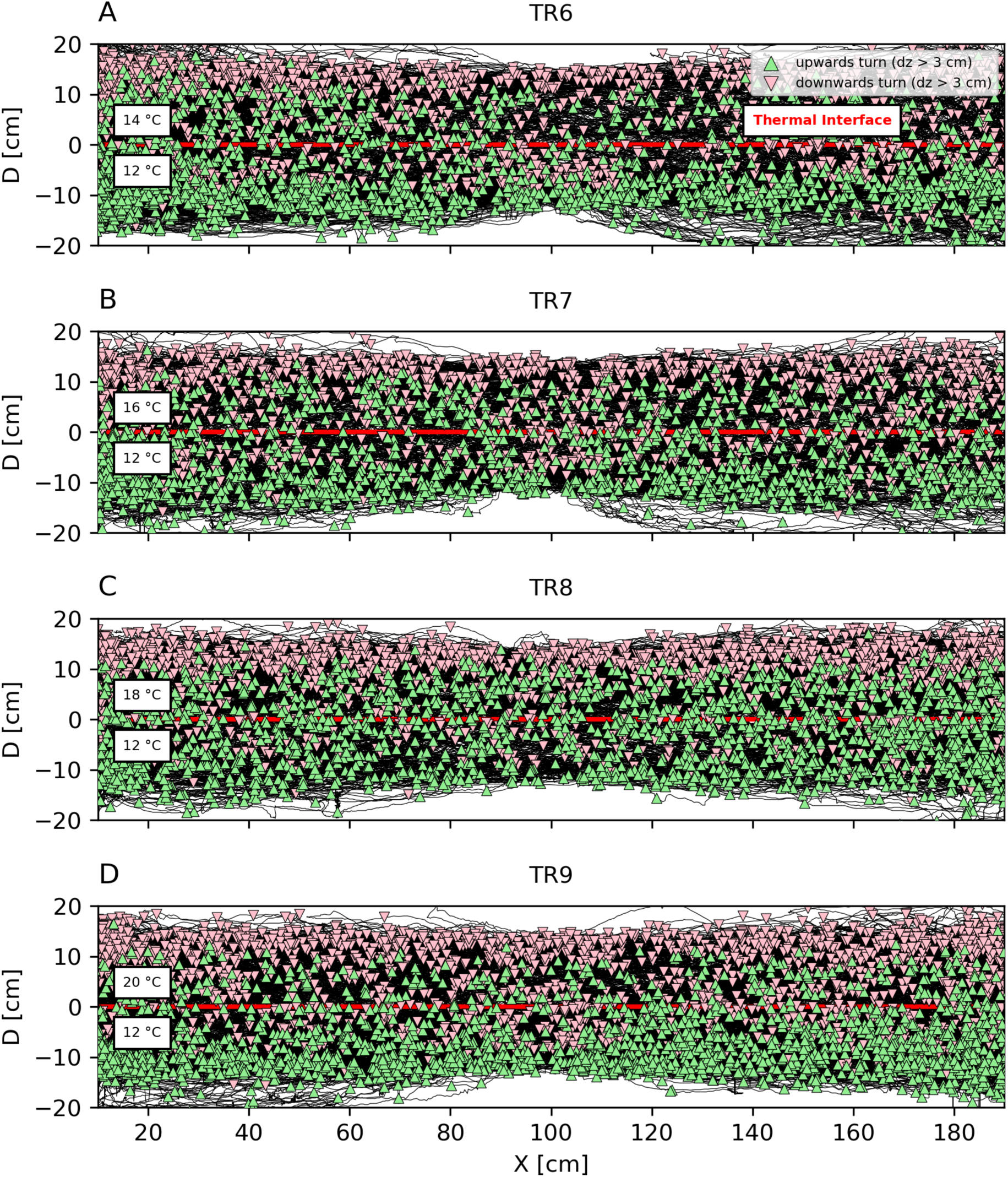
Warm treatments. Swimming trajectories (black lines) relative to the thermal interface (red line) for all fish across cold treatments (TR6 – TR9, *A*-*D*). The horizontal axis depicts the horizontal position in centimetres; the vertical axis depicts the instantaneous distance from the thermal interface. For D > 0 fish is located at a distance D [cm] above the interface and for D < 0 the fish is located below the temperature interface. Red and green triangles depict large downward and upward turns, respectively (see Supplementary Fig. 11 and 14 for derivation).

Extracting large turning points

**Fig. 14:**
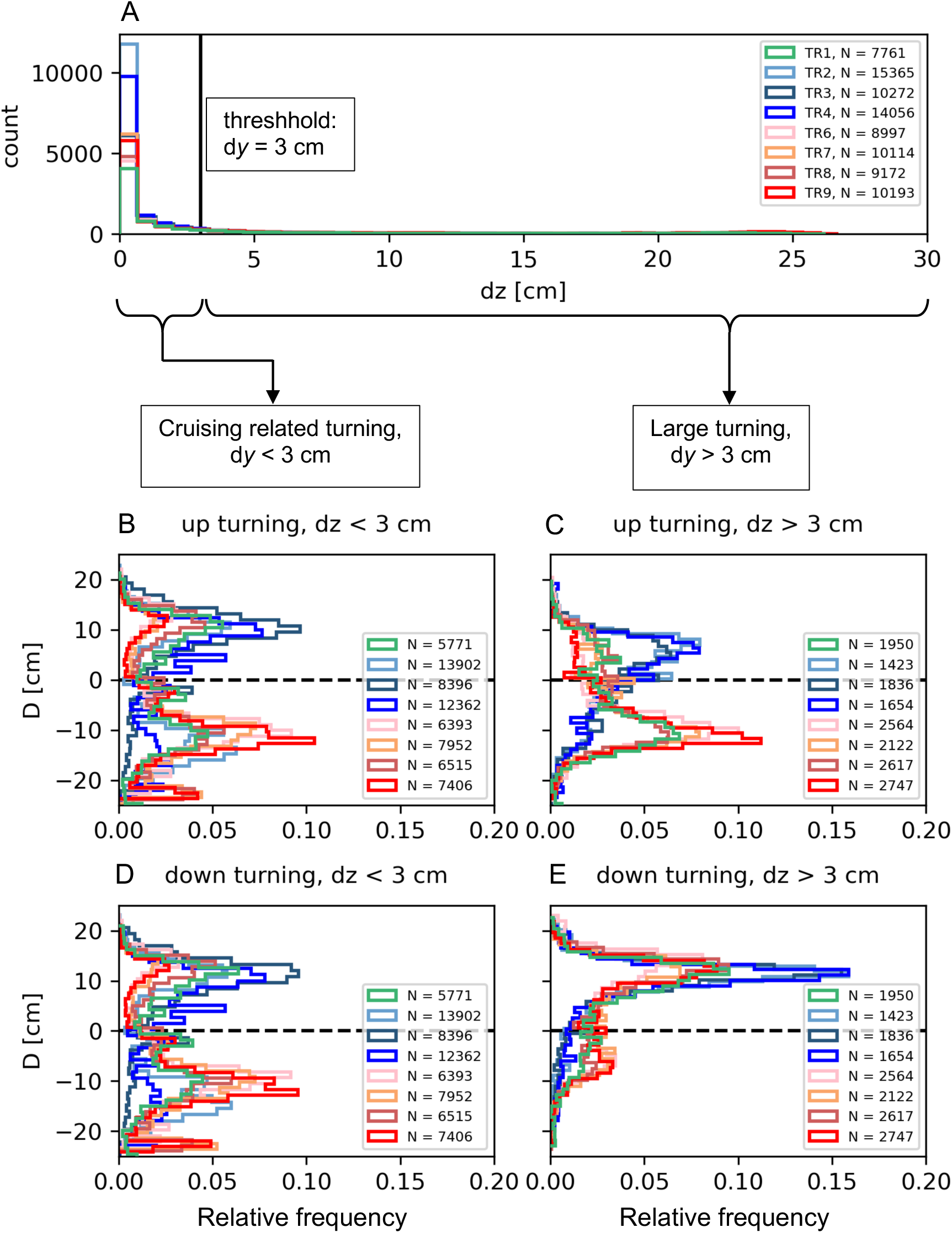
(*A*) Normalized histogram of the vertical distance dz between all consecutive turning points for all treatments (depicted by colors). A threshold of dz = 3 cm was used to differentiate turning events that result in small, cruising related vertical displacements (*B,D*) from vertical turns that resulted in large vertical displacement (dz > 3 cm) (*C, E*).

Number of cold and warm excursions (i.e. interface crossings)

**Fig. 15:**
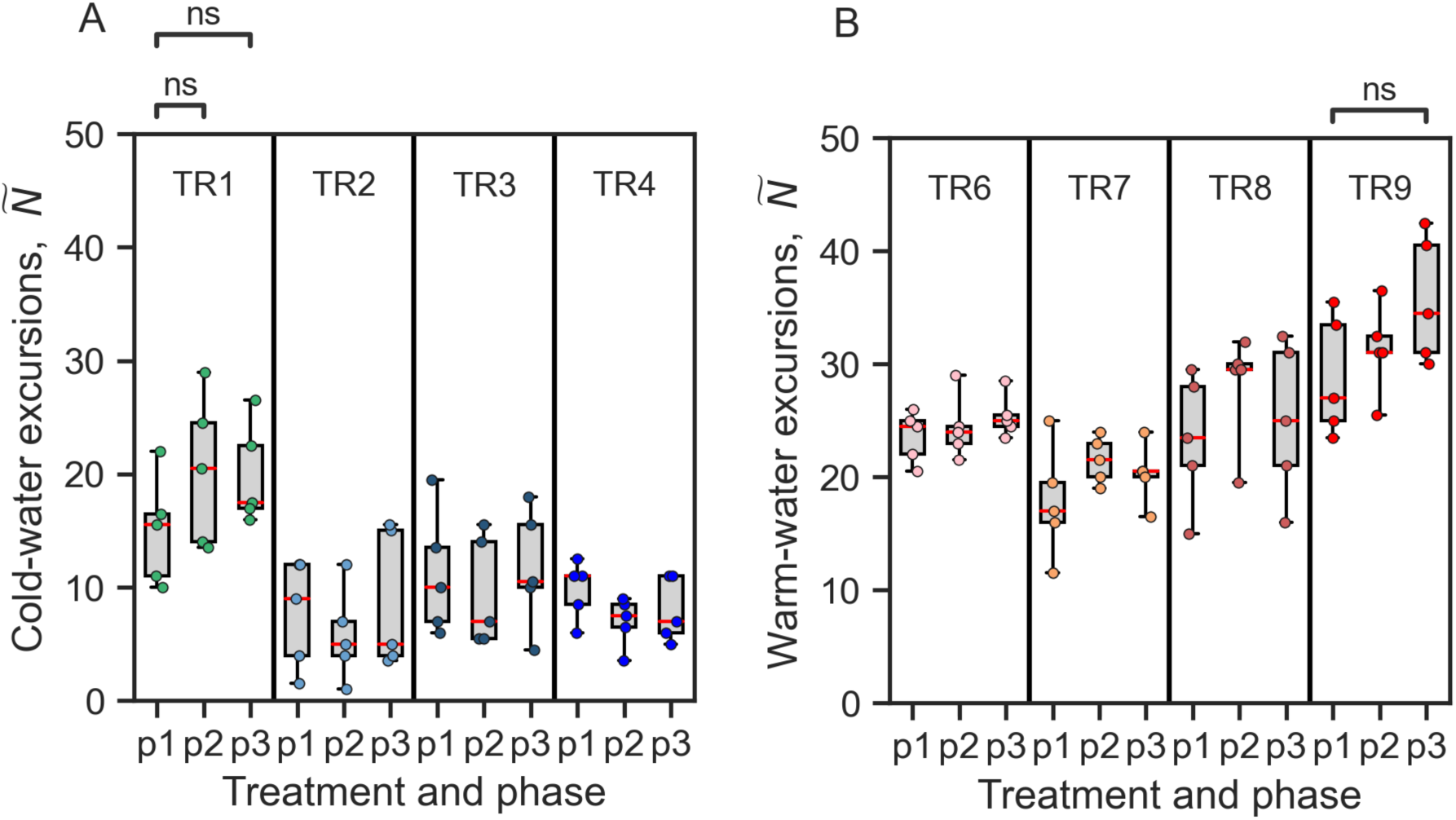
Number of performed cold-water excursions (*A*) and warm-water excursions (*B*) over time (p1 – p3). Boxplots show the median, whiskers extend to the full data range and the boxes are limited by 25^th^ and 75^th^ percentiles. Each dot represents the median across 4 individuals, tested simultaneously.

Duration of excursions

**Fig. 16:**
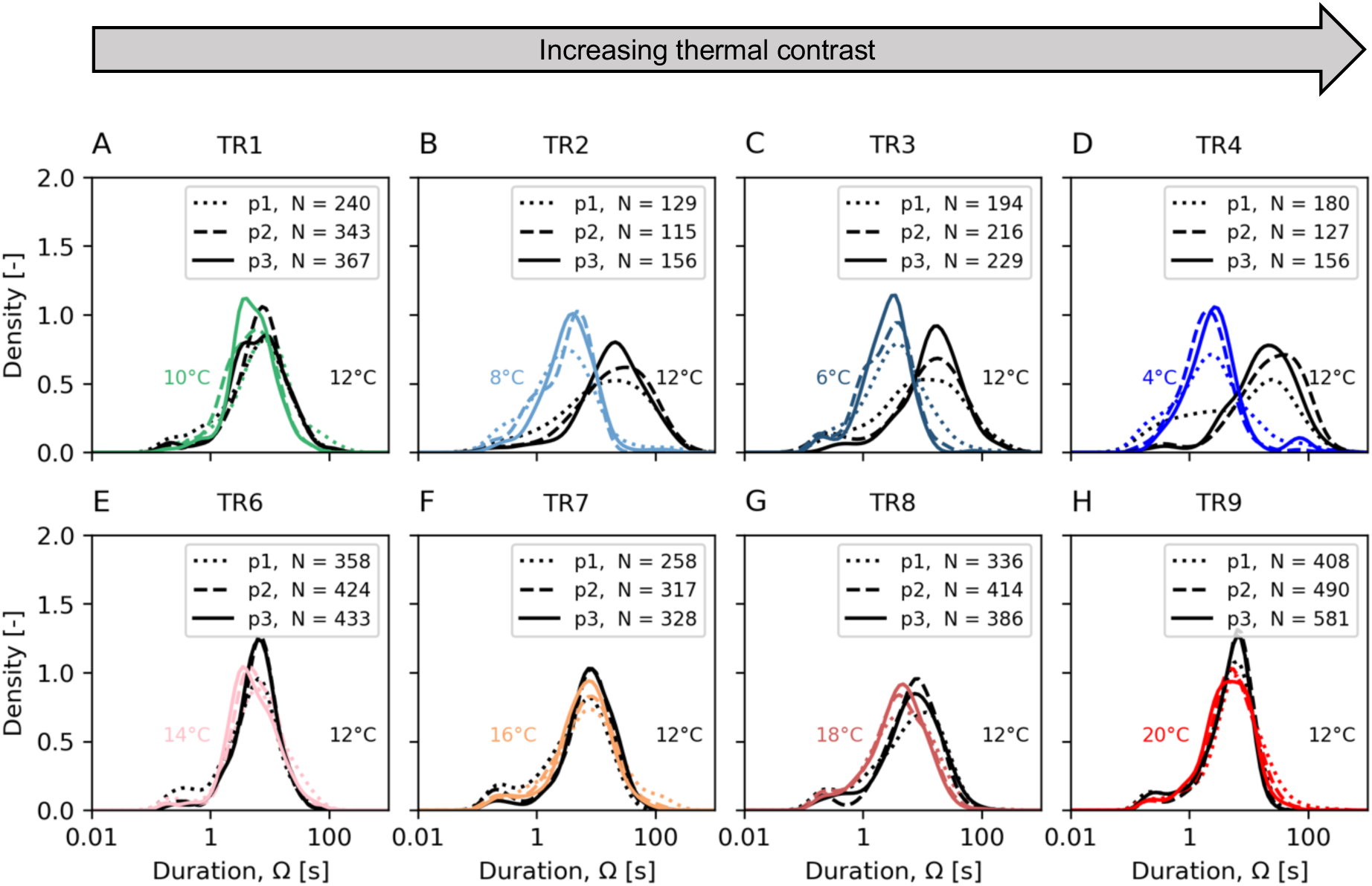
Probability density function of the duration of warm- and cold-water excursions for cold (*A-D*) and warm treatments (*E-H*) and across treatment phases (p1, p2 and p3). As a reference for comparison, black lines depict trajectory excursions in 12°C. Coloured lines show respective distributions in treatment water temperatures below (*A-D*) or above the thermal interface (*E-H*).

Excursions in warm treatments

**Fig. 17:**
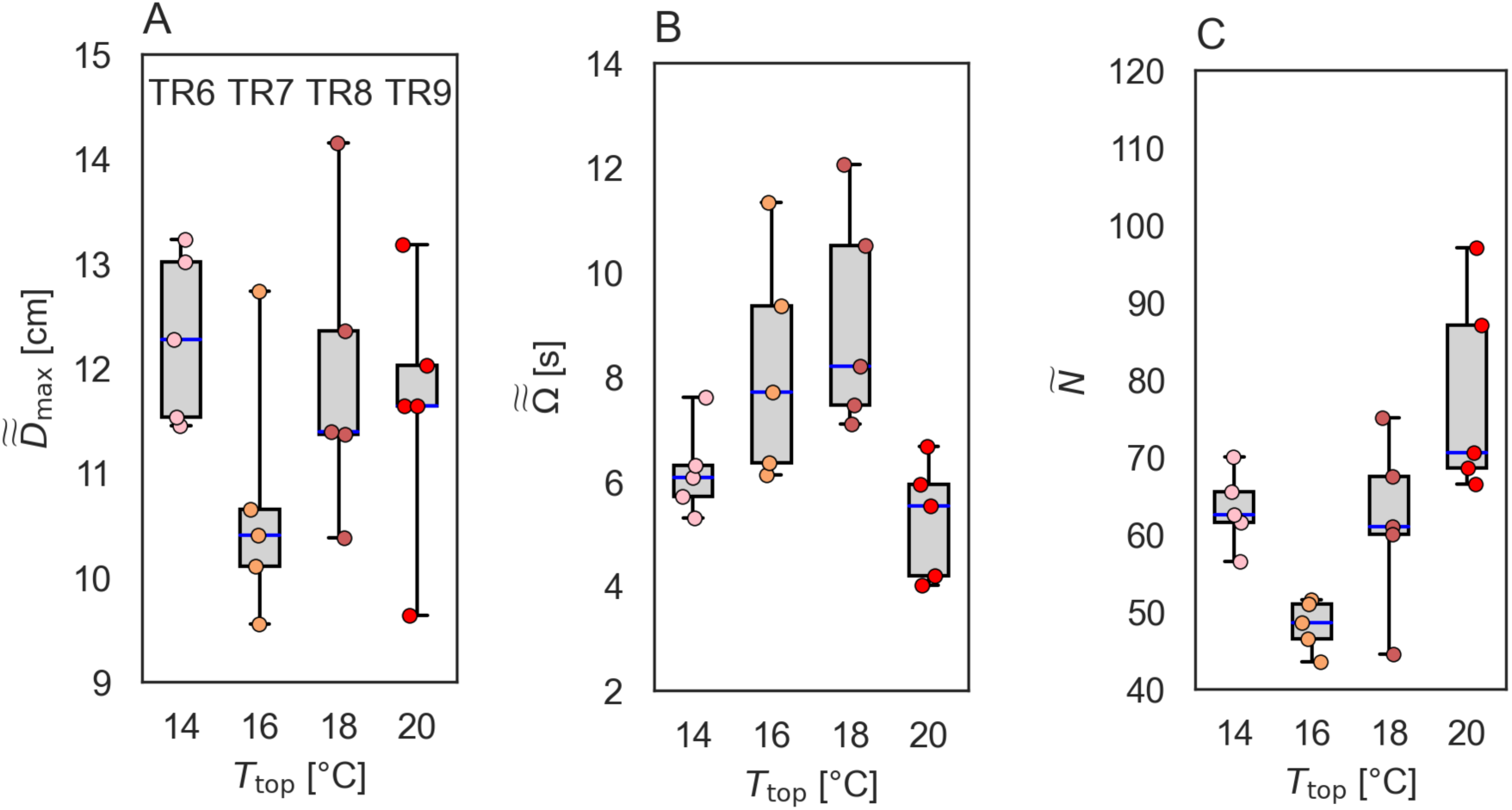
Warm-water excursions during warm treatments (TR6-TR9). For each quantity, the median of 4 individuals (tested simultaneously) was considered; hence each dot represents a single experiment. (*A*) Boxplots of the maximum (upwards) distance to the interface of warm-water excursions. The median of *D*_max_ was applied to all excursions performed by an individual fish and for all phases (p1, p2 and p3). (*B*) Boxplots of the median duration of cold-water excursions for fish. The median of *D*_max_ was applied to all excursions performed by an individual fish and for all phases (p1, p2 and p3). Each dot represents a single experiment. (*C*) Boxplot of the median number of warm-water excursions (upwards) for fish groups across warm treatments and experimental phases (p1, p2 and p3). Each dot represents the median response of four fish in a single experiment. Blue lines indicate the median, boxplots are limited by the 25^th^ and 75^th^ percentiles. Whiskers extent to the full range of observations.

Maximum distance to interface

**Fig. 18:**
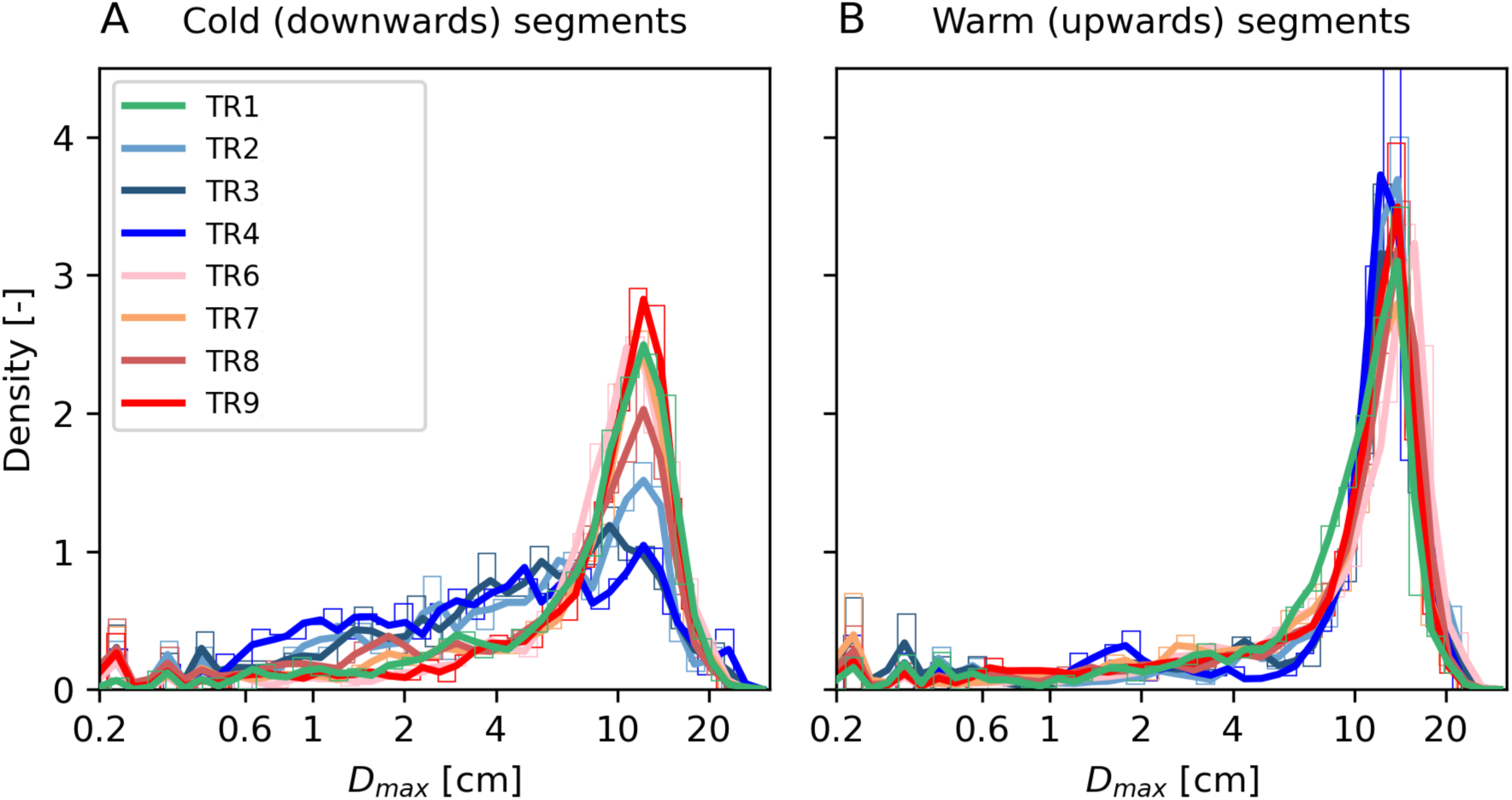
Probability density function of maximum distance to interface line *D*_max_ for cold (left) and warm (right) excursions for all treatments. Underlying normalized histograms are shown in thin colored lines.

**Fig. 19:**
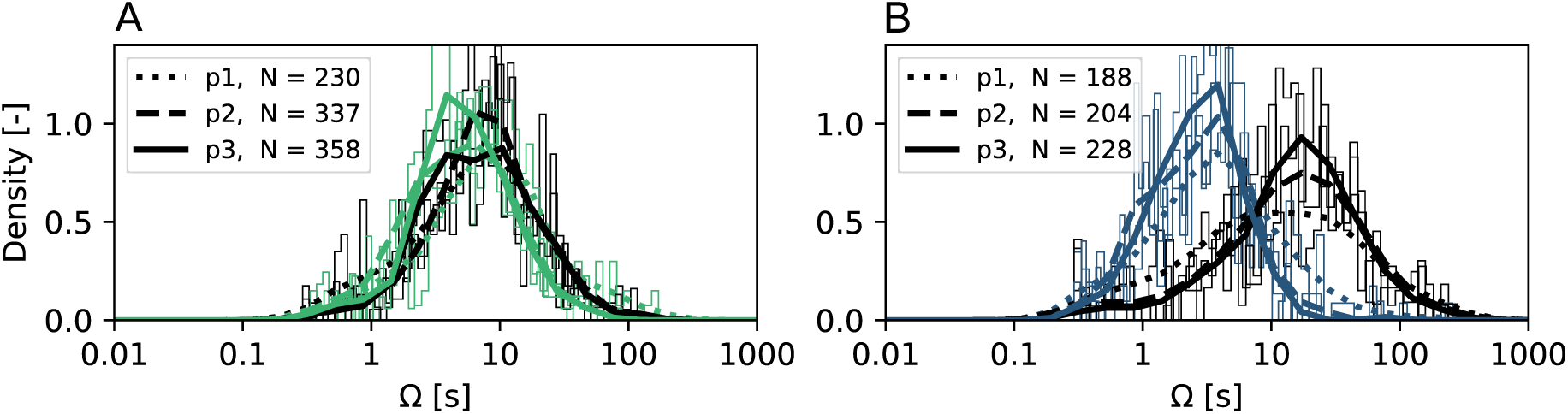
Underlying probability density distributions of the durations *Ω* of warm- and cold-water excursions for TR1 (*A*) and TR3 (*B*), respectively. The distributions of the warm- and cold-water excursion durations become increasingly distinct over exposure time for TR3 but not TR1. Excursions at 12 °C (above the interface) are shown in black, while excursions in the lower, colder water are depicted in the respective color (green: *T*_bottom_ = 10 °C; blue: *T*_bottom_ = 6 °C. Treatment phases are indicated by line type (p1–p3). Values of *N* indicate the total number of excursions of each type within that phase for 20 fish (as the excursions alternate, *N* is equal for cold and warm excursions). Underlying distributions are shown as histograms.

Absolute Swimming speed

**Fig. 20:**
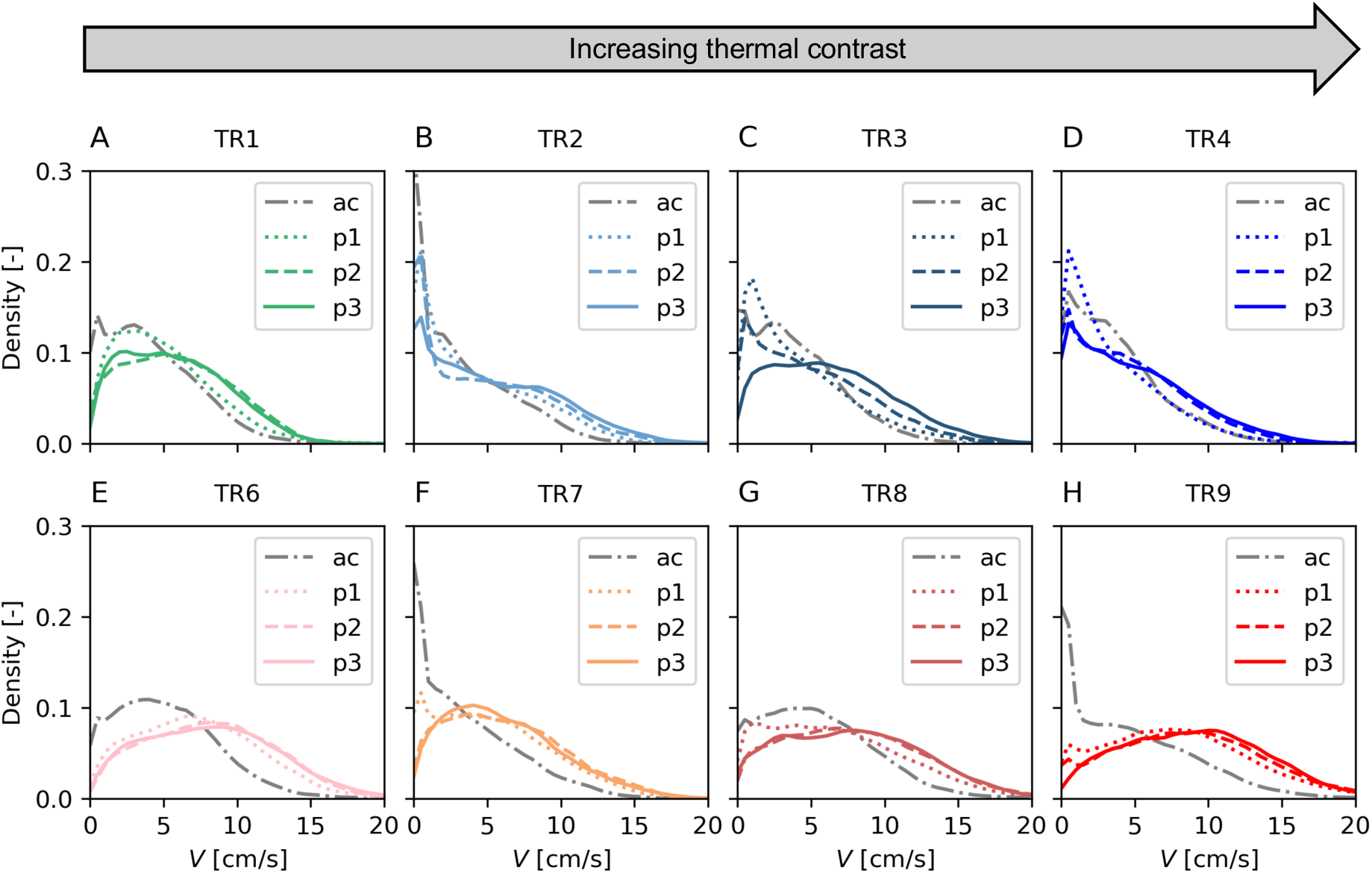
Probability density of absolute swimming speeds *V* per experimental phase (ac: acclimation; p1, p2 and p3) and treatment. Cold treatments (A-D) and warm treatments (E-H). Each lines indicates the normalized distributions of N = 6 min x 60 s x 24 fps x 20 fish = 172’800 observations.

Burst swimming in cold-water treatments

**Fig. 21:**
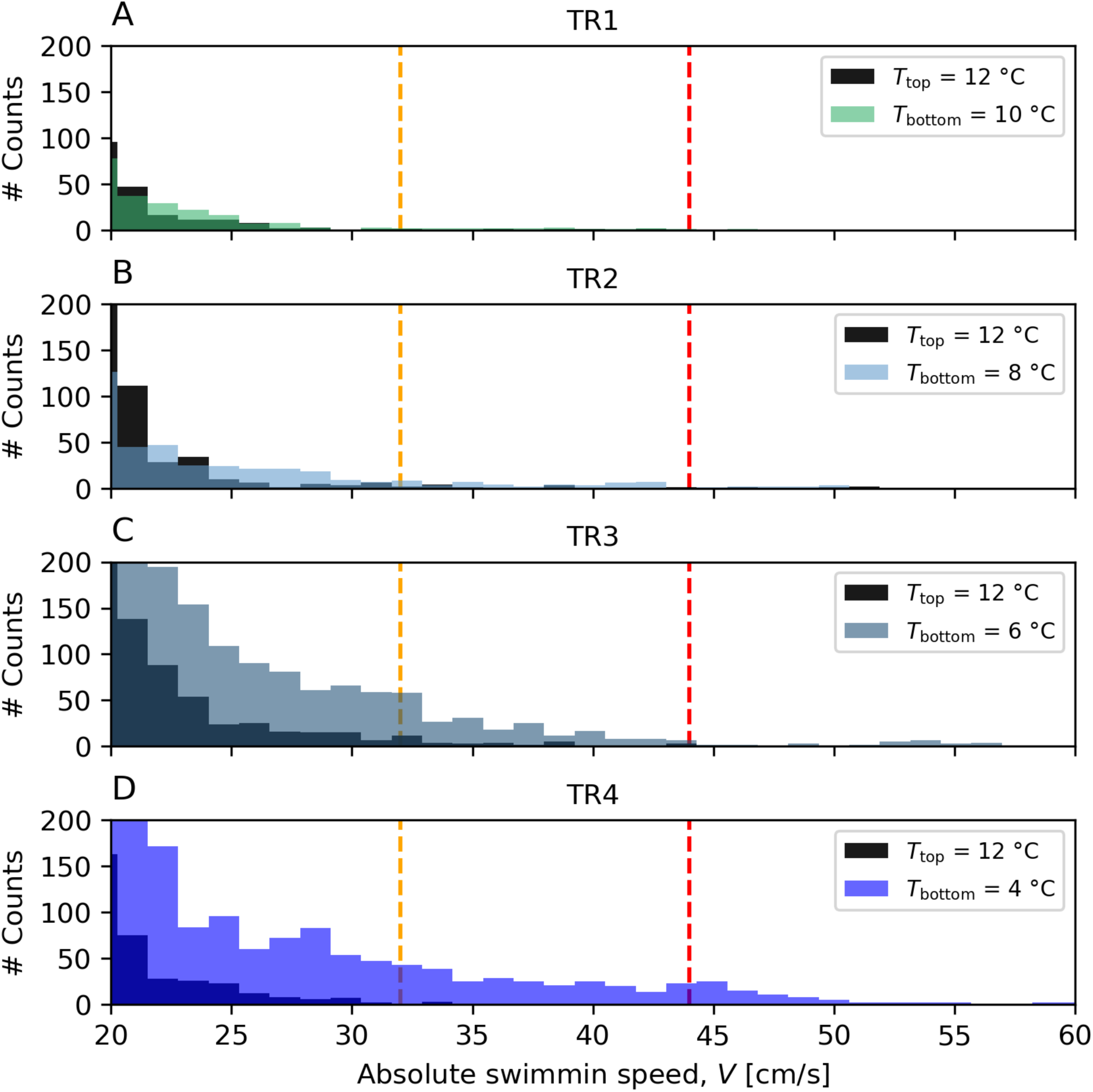
Extreme swimming speeds above *V* = 20 cm/s for TR1-TR4 (*A*-*D*). Black histograms show all observations above the thermal interface (*T*_top_ = 12 °C). Colored histograms display all observations below the thermal interface (in colder water). Dashed vertical lines depict literature thresholds for burst swimming speeds of rheophilic fish based on temperature (orange line: *T* = 12°; red line: *T* = 4 °C), body length (3.6 cm), and swimming durations (5 seconds) as proposed by (^77^) (see also Supplementary Text 4). High swimming speeds were most frequent during cold-water occupation in colder treatments (TR2-TR4).

Normed swimming speed in warm

**Fig. 22:**
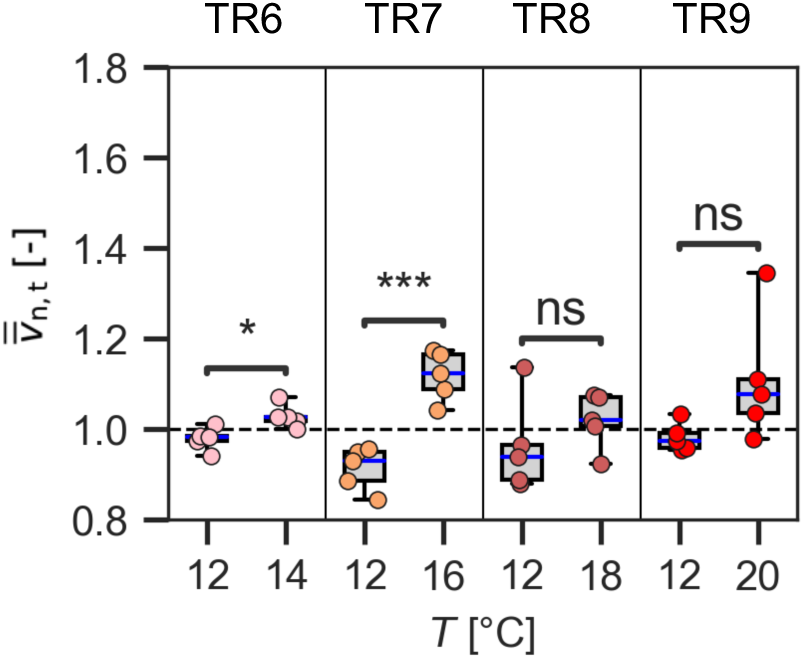
Mean swimming speed when above (in warmer water) and below (in colder water) the thermal interface. Speed was standardized by the average swimming speed of each individual during the treatment phases (p1–p3). Data was aggregated at the replicate level; hence *v̅̅*_n,t_ represents the average response of the fish group (four fish) within one replicate. Symbols represent the results of *t*-tests (ns, *p* > 0.05; *, 0.01 < *p* ≤ 0.05; ***, 0.0001 < *p* ≤ 0.001, see also Table S17). Whiskers extent to the full range of observations and red lined indicate the median.

**Fig. 23:**
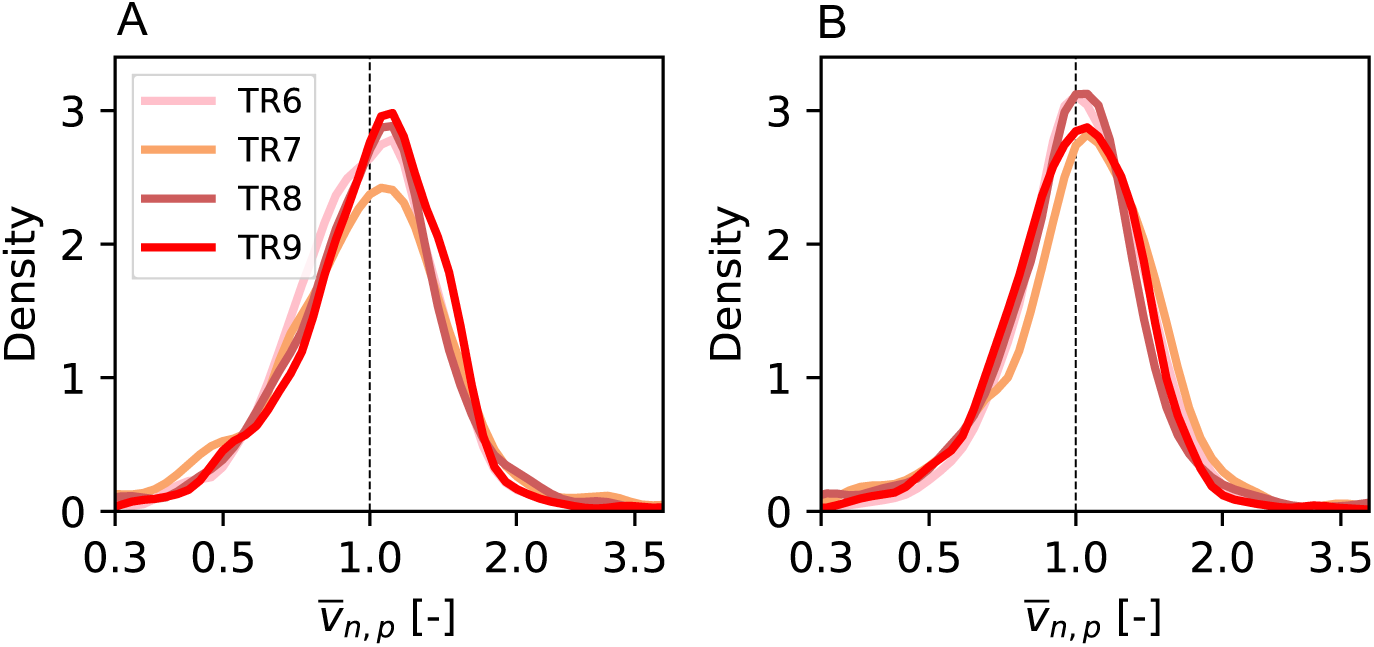
Averaged phase normalized swimming speed during warm water excursions (*A*) and cold-water excursions (*B*) in warm treatments (TR1, TR2, TR3 and TR4). Data has been transformed to log10 and a Kernel density estimator was fitted to the normalized histogram.

Averaged normalized speed vs. duration of excursions

**Fig. 24:**
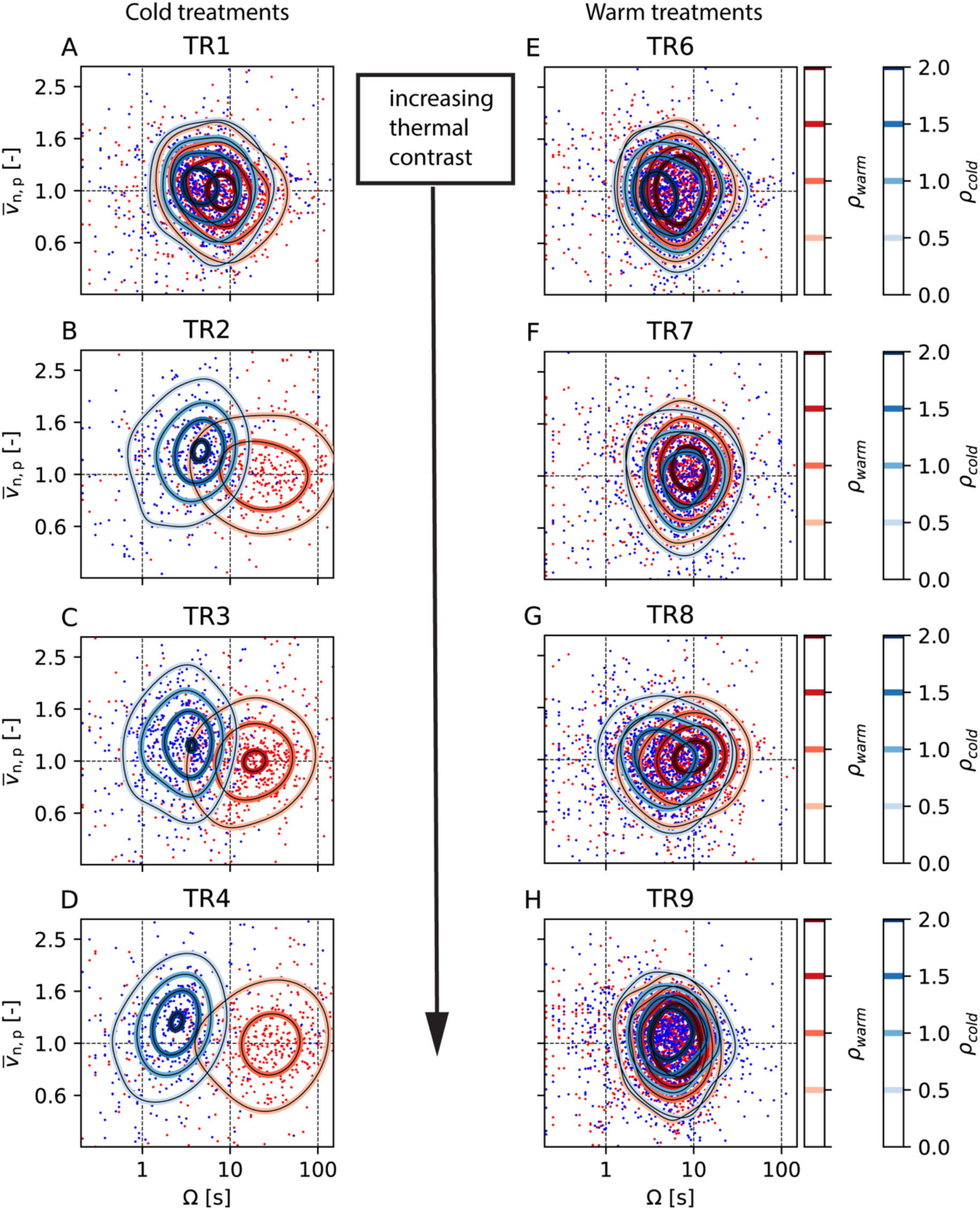
Normalized swimming speed *v̅*_n,p_ averaged over excursions plotted as a function of the duration *Ω* of each cold (blue points) and warm (red points) water excursion for all cold (*A* - *D*) and all warm treatments (*E* - *H*). To control for temporal trends, speed was standardized by the average swimming speed of each individual within each phase. Fitting a 2D Kernel density estimator to the point clouds (red and blue lines) reveals for cold treatments with *T*_Bottom_ ≤ 8 °𝐶 (TR2-TR4, *B*-*D*), clusters separate with respect to swimming speed (y-axis) and duration (x-axis).

Body cooling

**Fig. 25:**
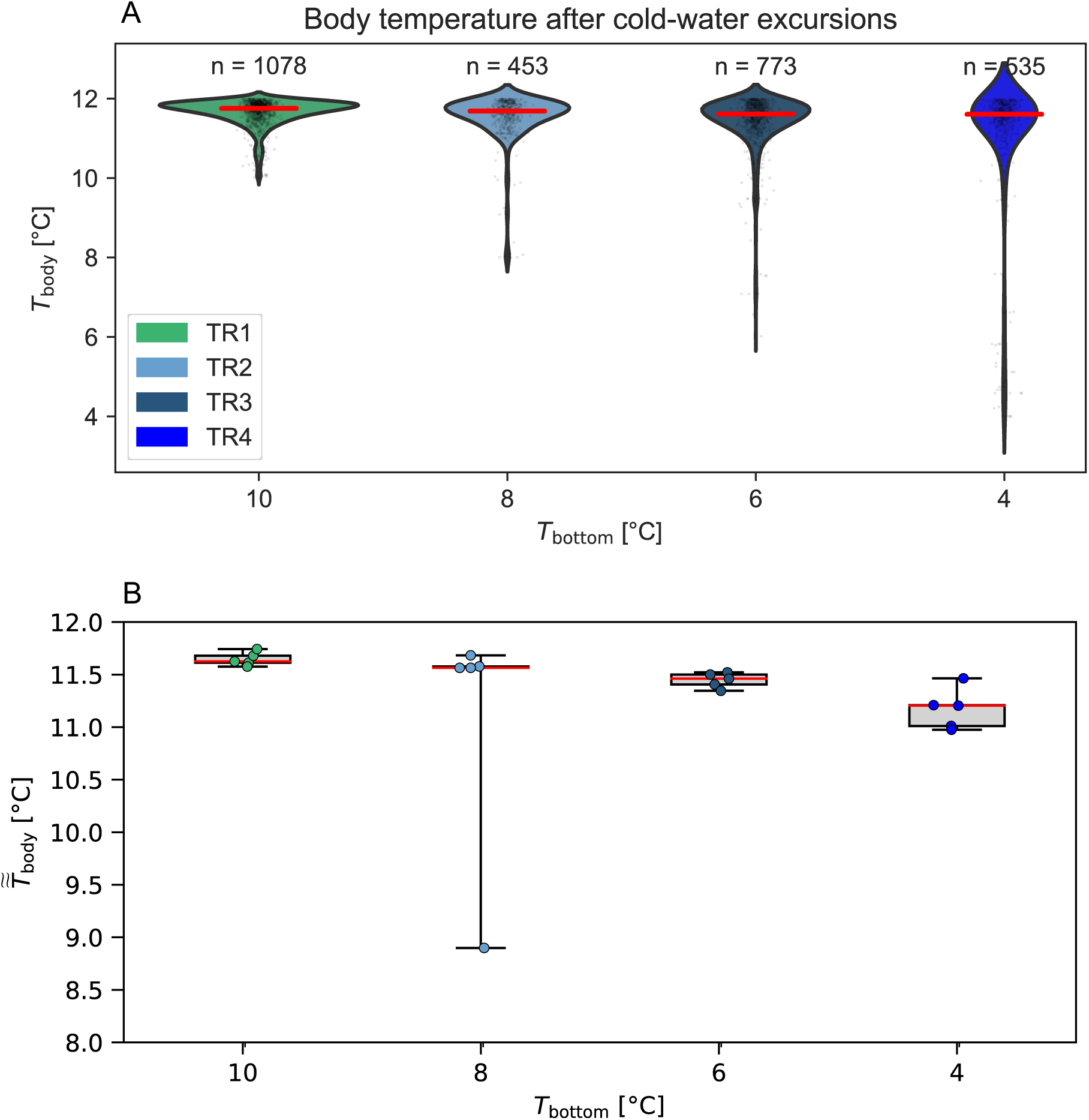
(*A*) Violinplots of simulated body temperature *T*_body_ after each performed cold-water excursion for cold-water treatments (TR1-TR4). Each dot is one excursion, red line indicates the median and *n* is the number of performed excursions by 5x4 = 20 fish within each treatment. The model (see SI Text) assumed fish body mass of m = 4 gramm, initial body temperature of *T*_0_ = 12°C and a cooling rate coefficient of k = 1.373 min-1. (*B*) Boxplot of the median body temperature (*T͌*_body_) after cold-water excursions for each experimental replicate. Red line indicates the median across experimental replicates.

Time spent

**Fig. 26:**
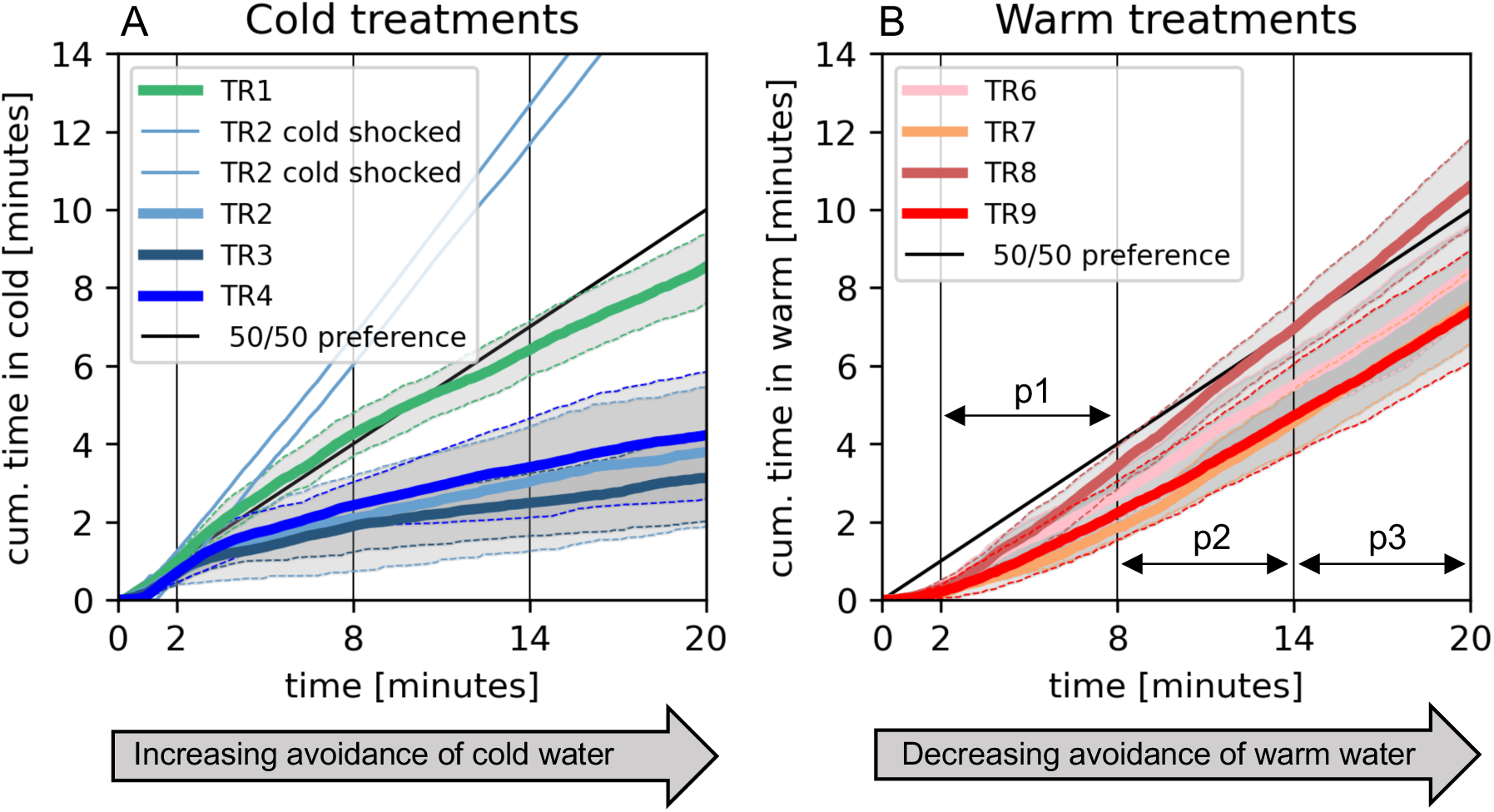
Cumulative time spent in incoming cold water (*A*) and warm water (*B*). Thick lines represent the averaged cumulative time spent in cold water on the replicate level. Thin lines depict the 84^th^and 16^th^ percentile across experimental replicates. Two cold shocked individuals (in TR2) were excluded from the average as outliers. Fish displayed opposing responses to incoming warm and cold water over time. Namely, fish decreased the occupancy of cold water, while in warm treatments fish enhanced occupancy of warm water (see also Fig. 8).

Water temperature and dissolved oxygen

**Fig. 27:**
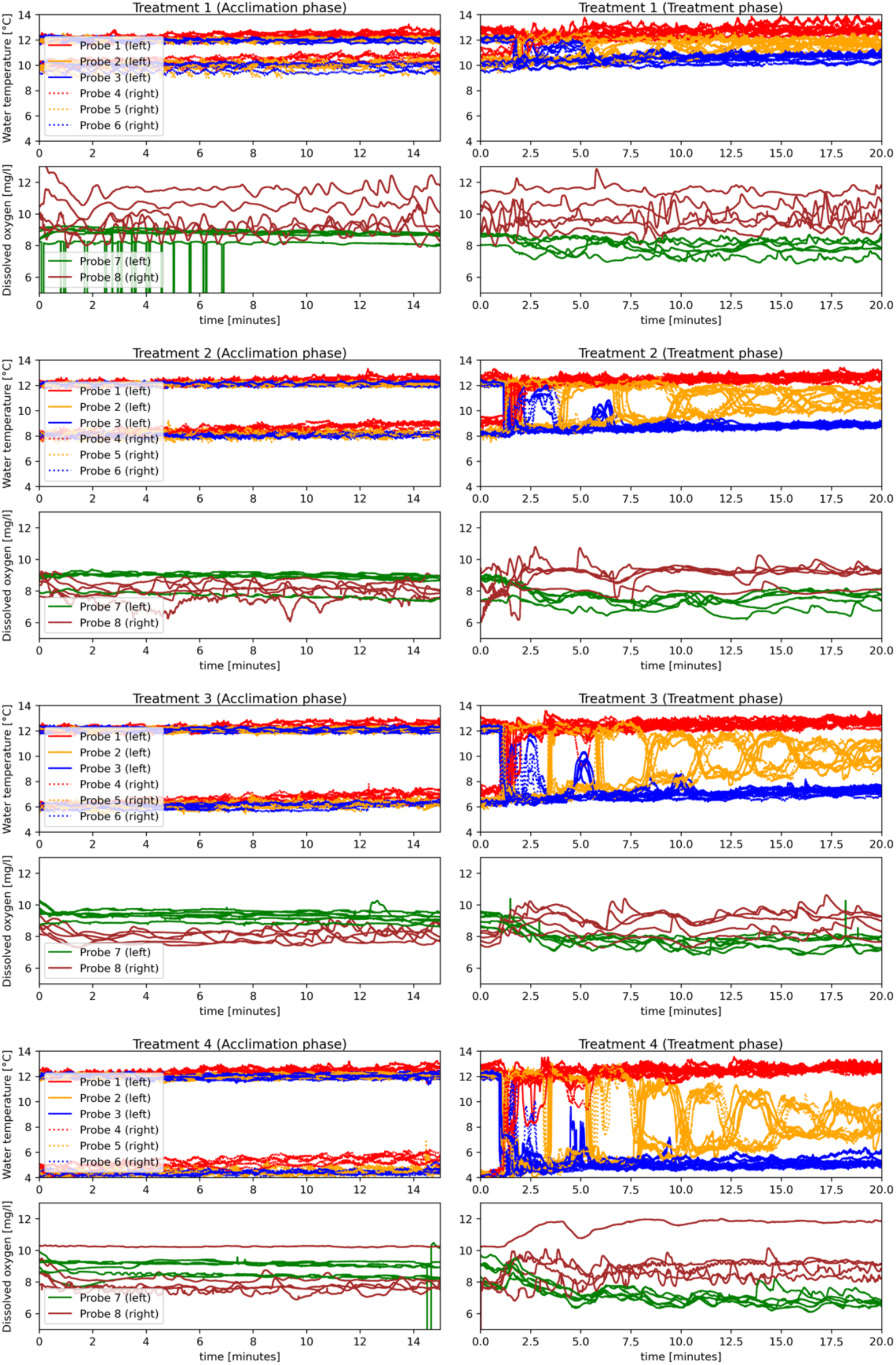
Cold treatments. Time traces of temperature [°C] and dissolved oxygen concentration [mg/L] in experiments during acclimation (left) and treatment phase (right). Temperature (Probe 1-6) was measured at three different depths (red, orange, blue) at both side walls of the experimental tank for all experiments (upper plots). Dissolved oxygen was measured on both sides of the tank. All measurements were continuously obtained at a frequency of 5 Hz.

**Fig. 28:**
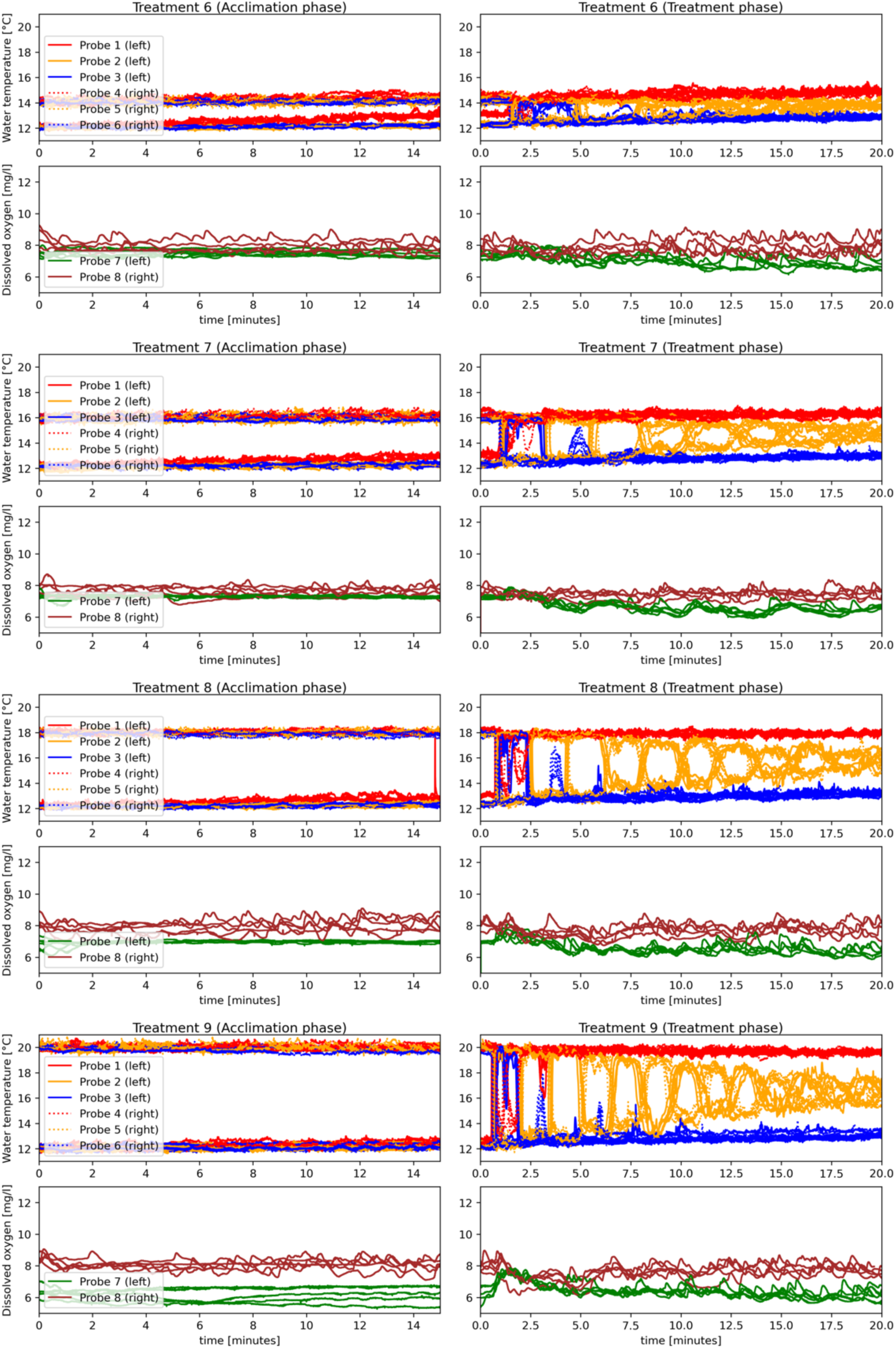
Warm treatments. Time traces of temperature [°C] and dissolved oxygen concentration [mg/L] in experiments during acclimation (left) and treatment phase (right). Temperature (Probe 1-6) was measured at three different depths (red, orange, blue) at both side walls of the experimental tank for all experiments (upper plots). Dissolved oxygen was measured on both sides of the tank. All measurements were continuously obtained at a frequency of 5 Hz.

**Fig. 29:**
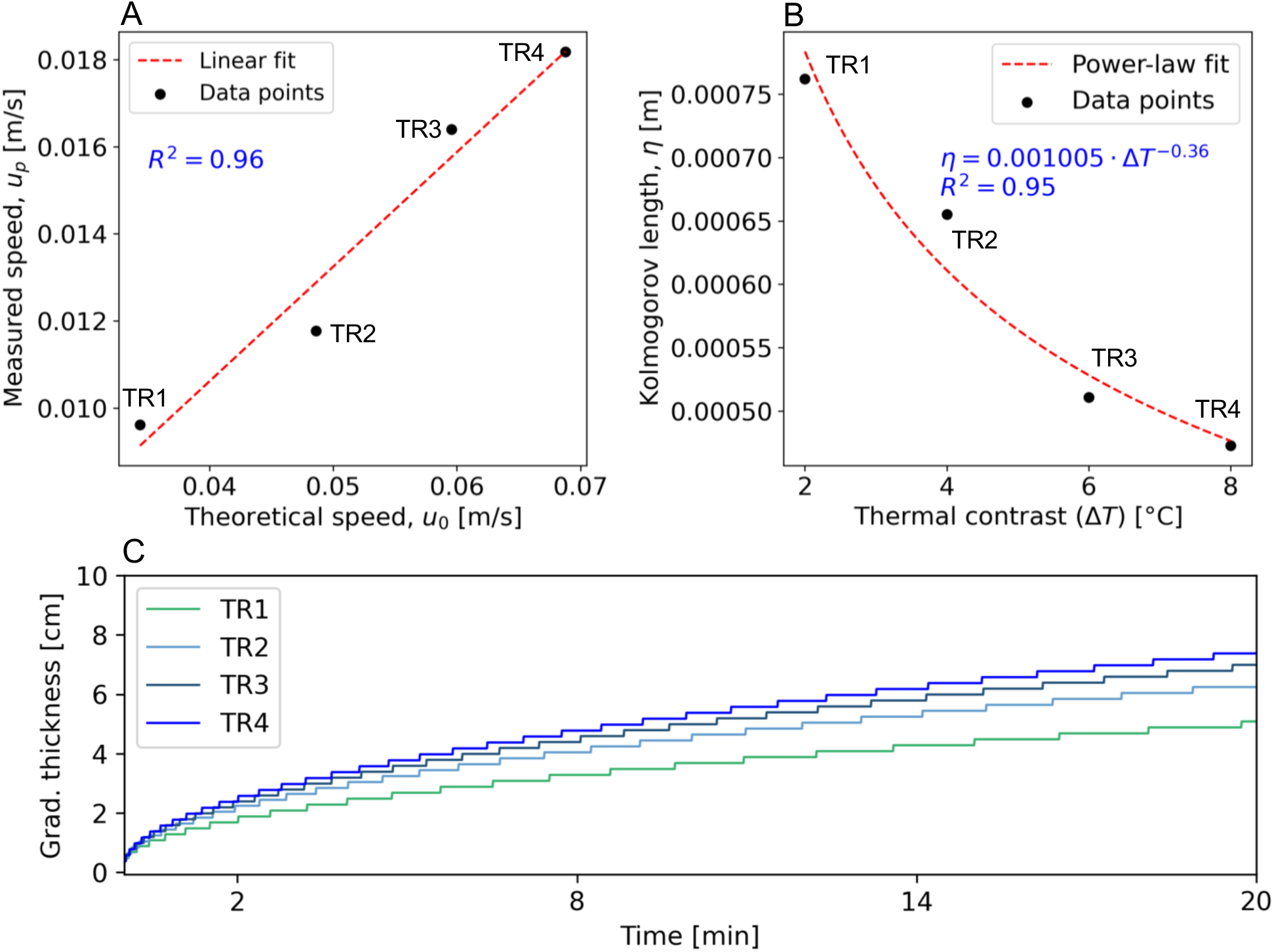
The theoretical large-scale gravitational velocity 𝑢_0_, plotted against the cold current propagation speed 𝑢_-_obtained from experimental videos of cold-water treatments (TR1–TR4). A linear regression fit (red dashed line) is shown. (B) The Kolmogorov length 𝜂 as a function of the temperature contrast Δ𝑇. The black dots represent experimental data, while the red dashed line indicates a power-law fit. (C) Temporal evolution of the thermal interface thickness, influenced by turbulence and diffusion. A 2D numerical simulation was conducted over a 20-minute period, using an initial interface thickness estimated as 𝛿_k_ ⋍ 3.7 𝜂 (see Methods). The spatial domain was discretized into 1 mm^2^ grid cells (dx = dy = 0.001 m) and a constant thermal diffusivity of D = 1.43e-07 m^2^/s was assumed. Results indicate that, over the course of the experiment, the gradient thickness 𝛿_k_ increased over time, rising to values larger than the length the individual fish.

Trajectories

**Fig. 30:**
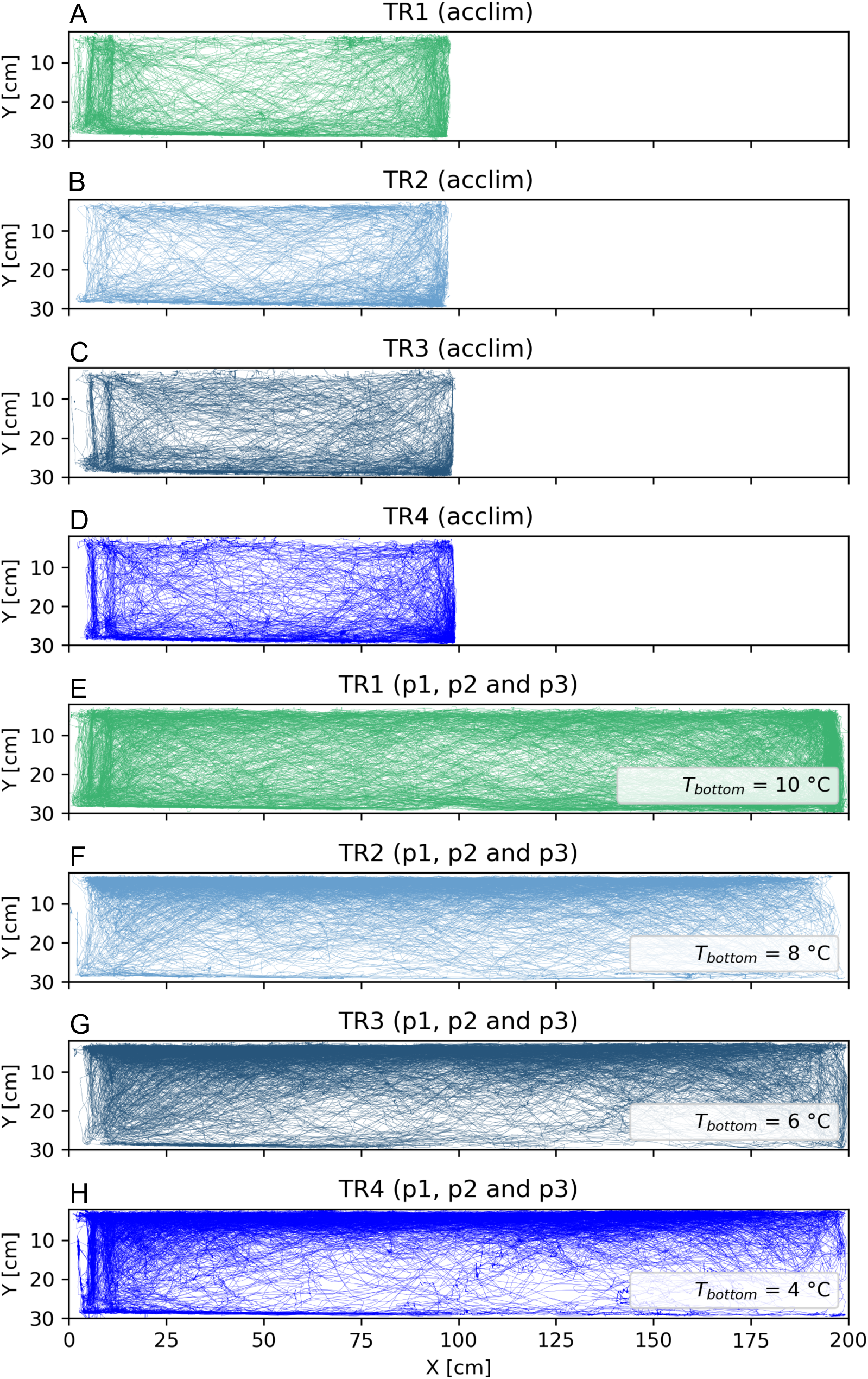
Cold treatments. Trajectories for all fish during acclimation phase (A - D) und treatment phases (E - F).

**Fig. 31:**
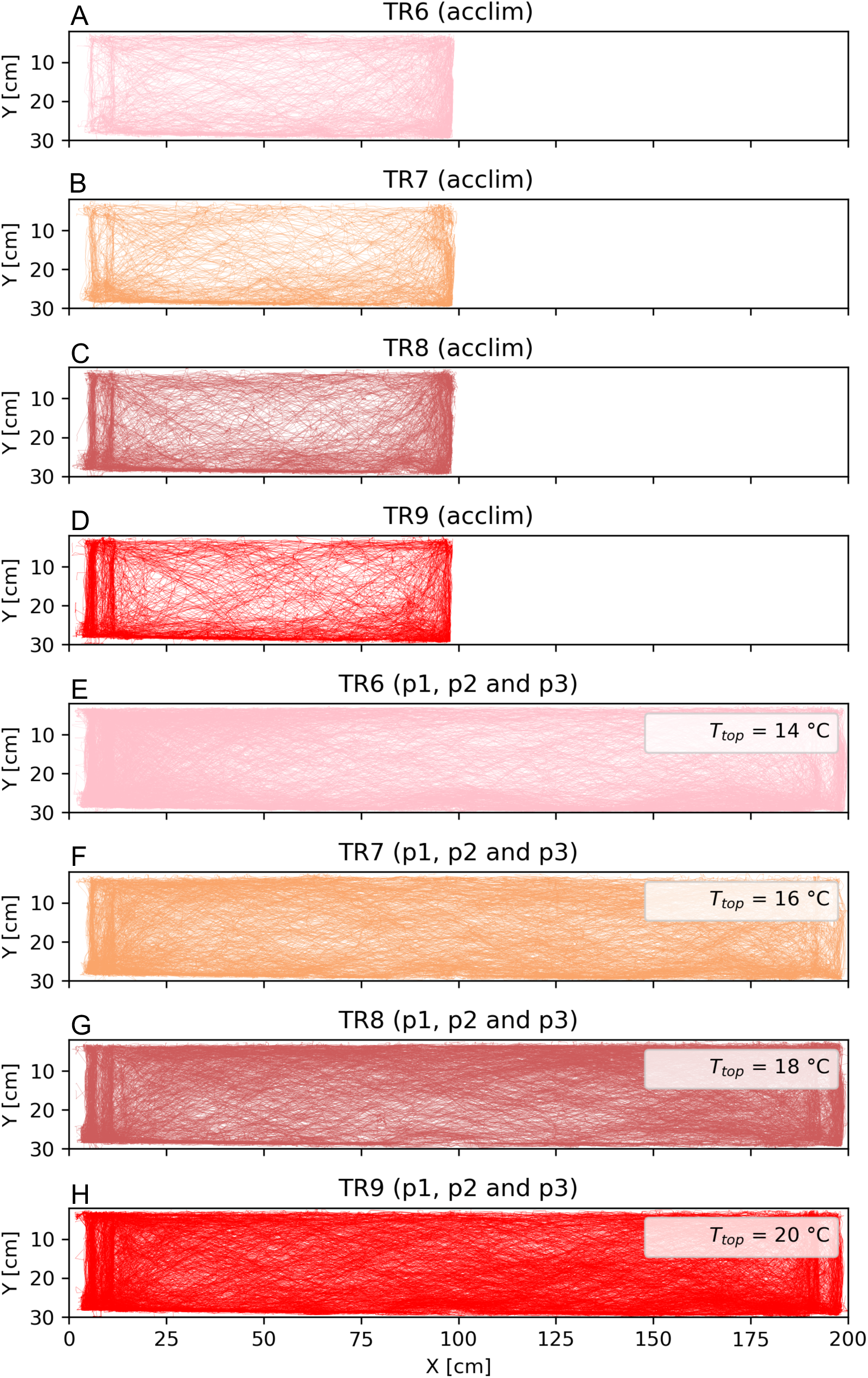
Warm treatments. Trajectories for all fish during acclimation phase (*A* - *D*) und treatment phases (p1, p2 and p3, *E* - *F*).

Fish-tracking

**Fig. 32:**
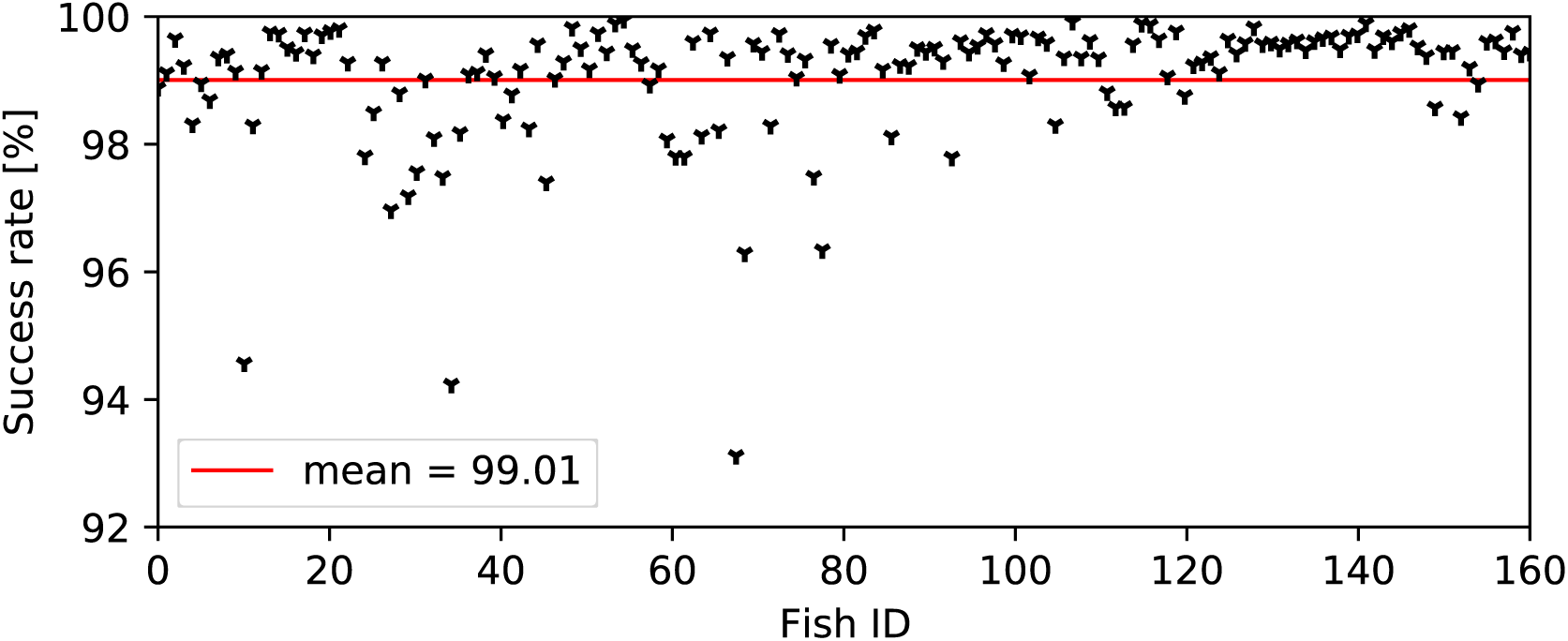
Tracking success rate for each individual during experimental phases (p1, p2 and p3). We captured the trajectory for on average 99% of the time. Low values are explained by fish occluding over longer periods of time (without moving) or being located behind the dissolved oxygen probes. For these cases a second camera perspective was used to resolve the occlusions and data gaps were closed using linear interpolation. The open-source tracking software “TRex” (67) was used to track fish at a temporal resolution of 24 frames per second.

Color splitting

**Fig. 33:**
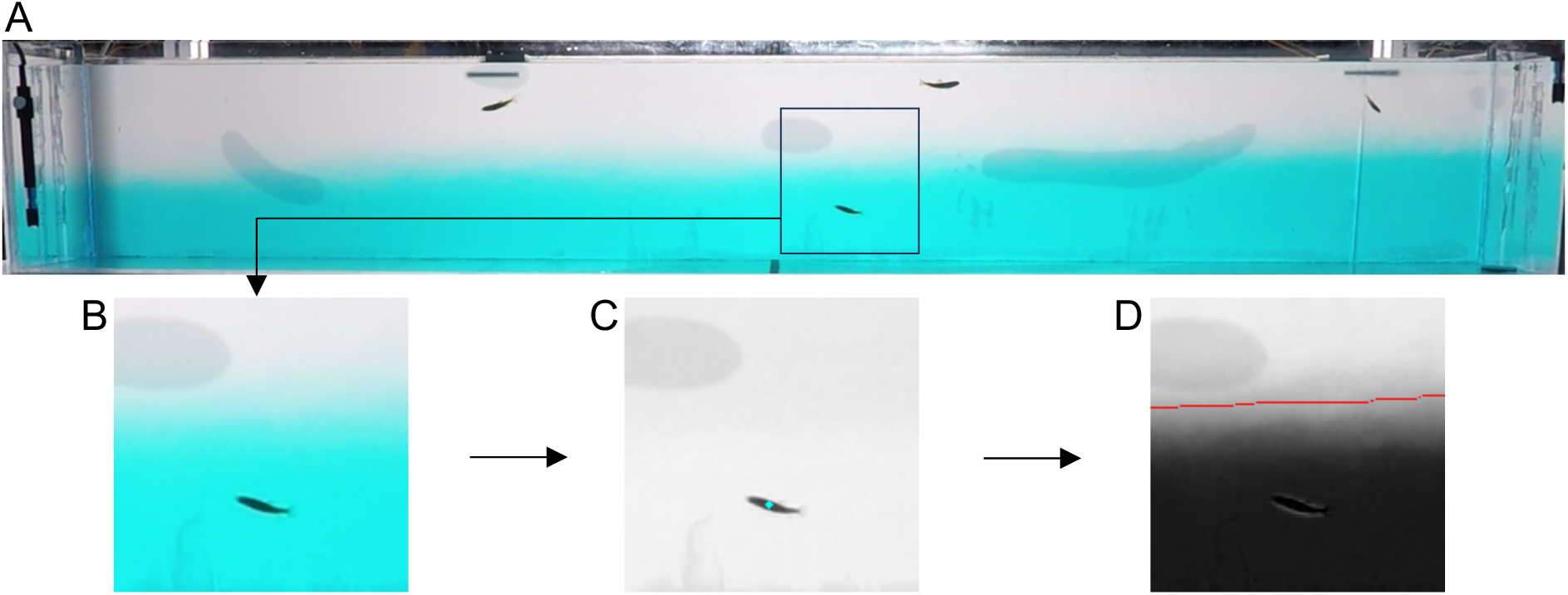
(A) Rotated and cropped RGB image of the side camera (GoPro hero7). (B) Subsection of RGB image. (C) Red channel. The turquoise dot indicates the fish’s center of gravity for this time instance, derived from tracking in TRex. (D) Green channel. Red line indicates the thermal interface, derived by custom made python scripts.

Control experiment

**Fig. 34:**
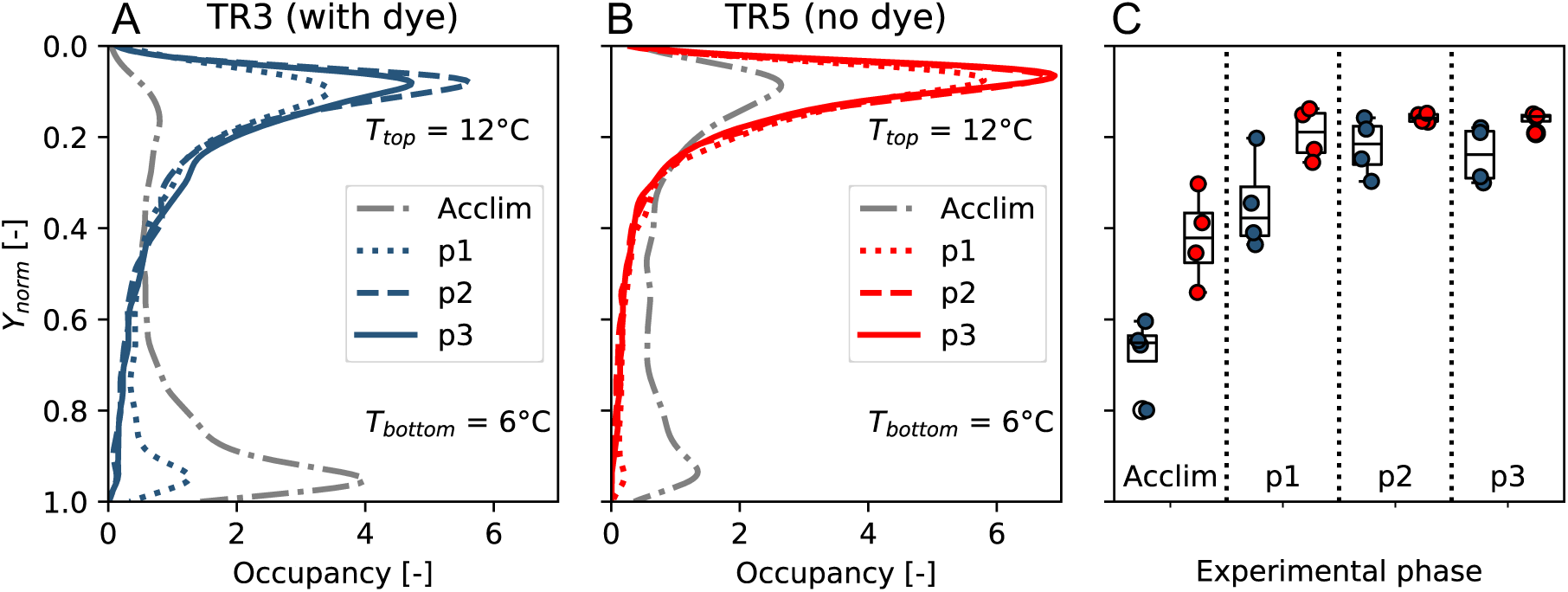
(A *and B*) Probability density functions of fish depth during *experimental phases* (*acclim., p1, p2 and* p3) of the experiments for cold-water treatment 3 (TR3*, A, N* = 16 fish) and its control without the addition of blue dye (TR5, B, *N* = 16 fish). Both cases had identical thermal conditions (T_bottom_ = 6 °C, T_top_ = 12 °C) above and below the thermal interface. The depth was normalized so that *Y*_norm_ = 0 is the water surface and *Y*_norm_ = 1 indicates the bottom of the tank. Due to a camera failure one of the control experiments could not be processed, hence the control is based on 4 replicate*s*, while TR3 is based on 5. (C) Boxplots aggregated on the fish group level did not show significant differences with regards to the vertical relocation across experimental phases.

**Fig. 35:**
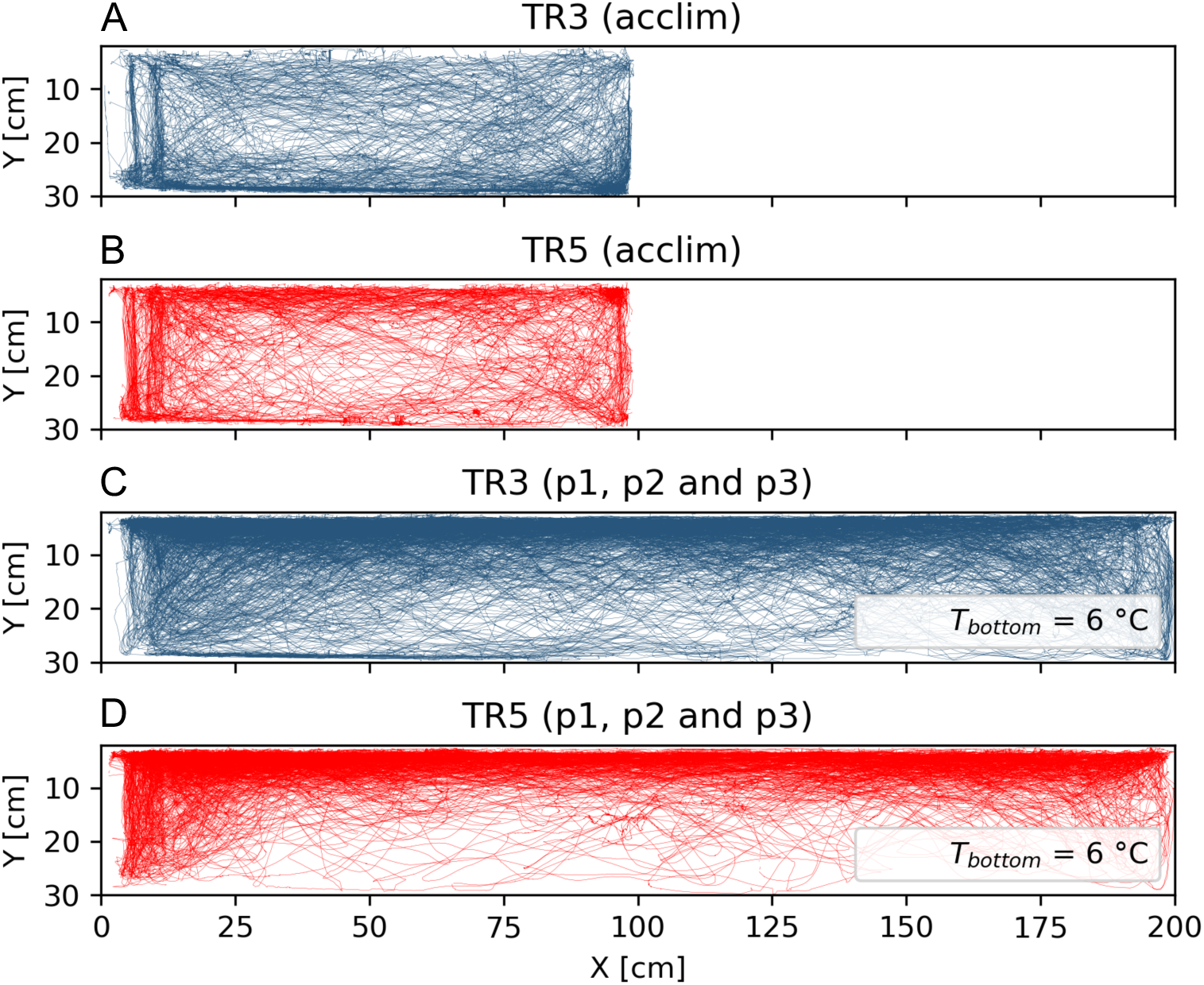
2D-swimming trajectories for TR3 (*A* and *C*) and its control without blue dye (TR5, *B* and *D*) during 6 min acclimation (*A* and *B*) and later experimental phases (p1, p2 and p3, *C* and *D*). Both treatments had identical temperature conditions (T_left_ = 12 °C; T_right_ = 6 °C).

Movement of thermal interface

**Fig. 36:**
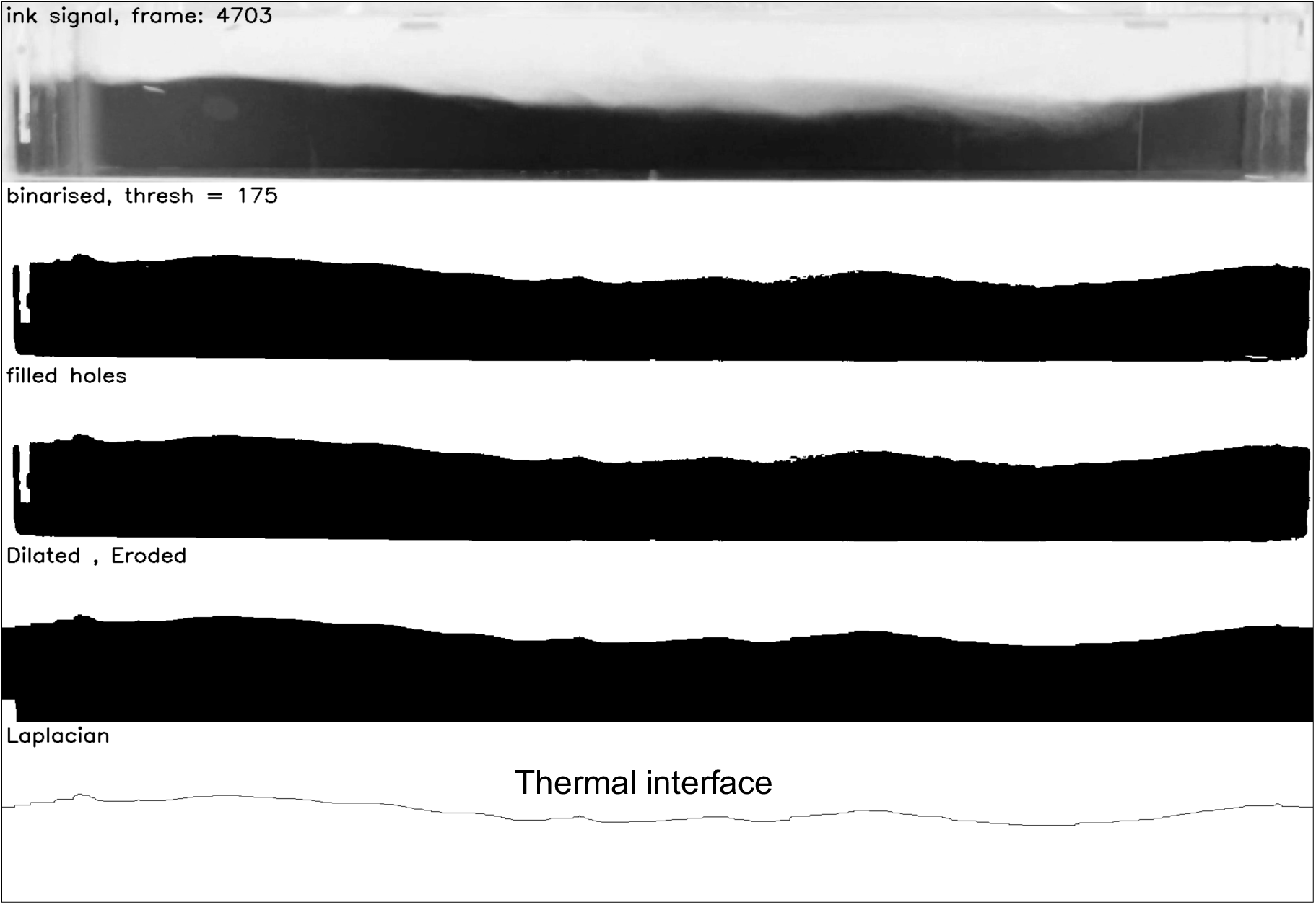
Image operations in python (opencv) to derive the instantaneous position of the thermal interface. As the lock exchange flow is primarily a 2D phenomenon and the width of the tank is narrow (20 cm) as compared to its length (200 cm), the temperature interface was approximated as a 2D line (*X*(t),*Y*(t)). The interface coordinates were derived for each time frame using a custom-made image analysis pipeline (python, opencv).

Binarization of Trajectory

**Fig. 37:**
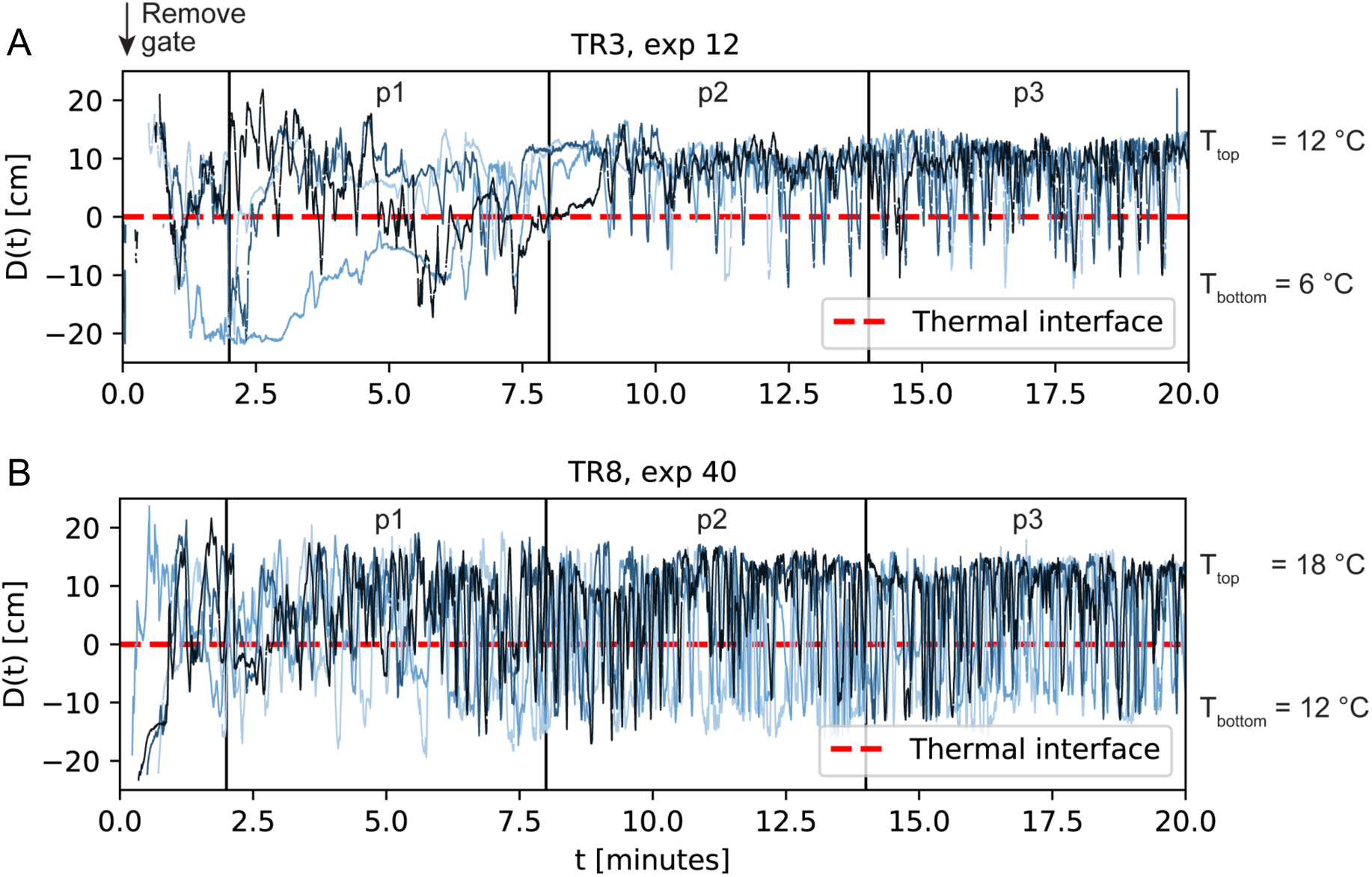
Vertical distance to thermal interface line. The temporal progression of the vertical distance to the thermal interface line, D(t), is shown for 4 individuals in an experiment of cold treatment 3 (TR3) and warm treatment 8 (TR8). The vertical gate was removed at time *t* = 0. Negative values of D(t) depict times when the center of gravity of the tracked fish was below the thermal interface hence in the colder water and vice versa. Each sign change of D(t) depicts an interface crossing. Data gaps within the first two minutes arise from the fact that the gravity current did not yet fully span the entire horizontal dimension of the tank (i.e., in this period was excluded from our analysis, fish may neither be located below nor above but laterally displaced from the thermal interface). Shortly after the current reaches the wall, maximum distances |𝐷(𝑡)| of up to 20 cm are possible (see also Movie 1). After the vertical stratification becomes stationary (t ≈ 10 minutes), the interface is approximately located at half the tank depth (*h* = 14 cm) and values for |𝐷(𝑡)| are below 14 cm.

Velocity smoothing

**Fig. 38:**
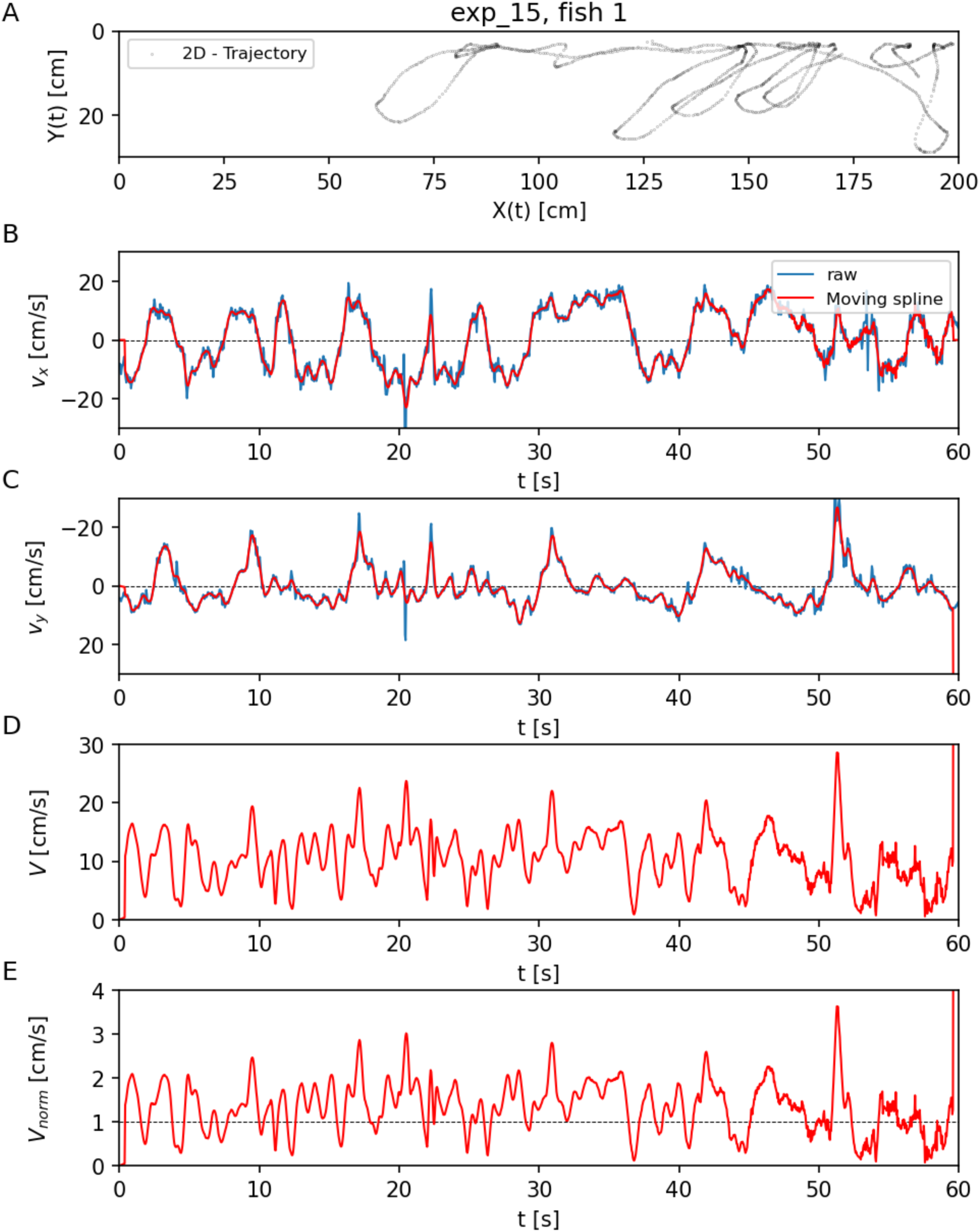
Example time traces illustrating the derivation of fish swimming speed (*V*) and normed swimming speed *V*_norm_ from the raw tracking output. (A) Raw trajectory output from tracking software (TRex) for a single fish during 1 minute of treatment 3 (*T_bottom_* = 6°C). (B,C) Horizontal and vertical velocity component (*v*_x_, *v*_y_) derived from raw output (blue line) and after applying the spline smoothing (Lüthi et al. 2005). (D) Absolute 2D-swimming speed *V* based on smoothed velocity components. (E) Normalized swimming speed *V_norm_* speed derived as *V_norm_* = *V(t)* / *V_mean_*. The dashed line indicates the average swimming speed during the entire treatment duration (p1, p2 and p3 combined).

**Fig. 39:**
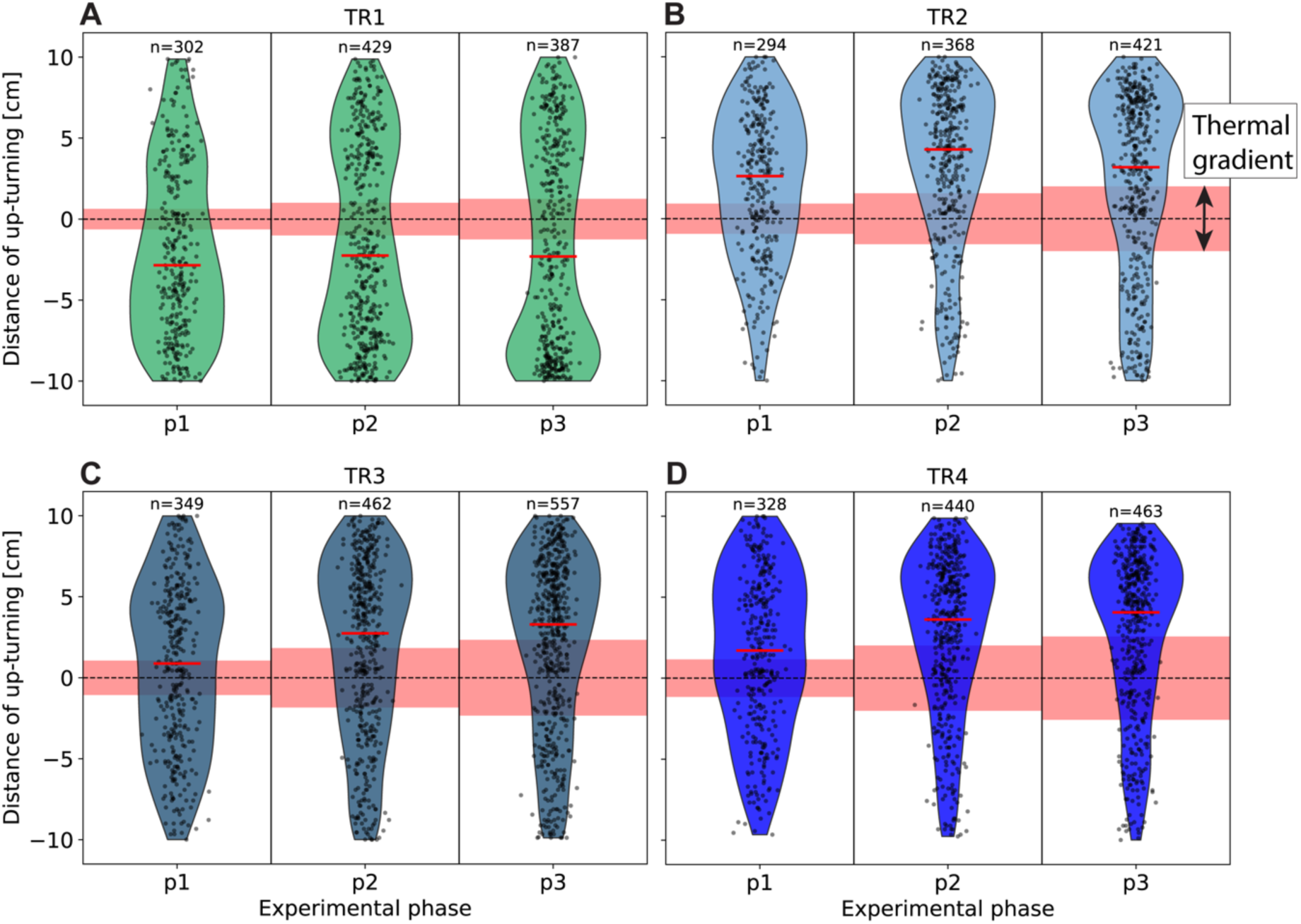
Violin plots showing the vertical distance from the thermal interface at which up-turning events occurred during cold-water treatments (TR1–TR4; panels A–D). Each black dot represents a single up-turning event, and the red line indicates the median distance. Data are grouped by experimental phase (p1–p3). The dashed line denotes the position of the thermal interface (0 cm), and the red shaded area indicates the estimated thickness of the vertical thermal gradient (see Methods).

**Supplementary Tables**

**Table 1:**
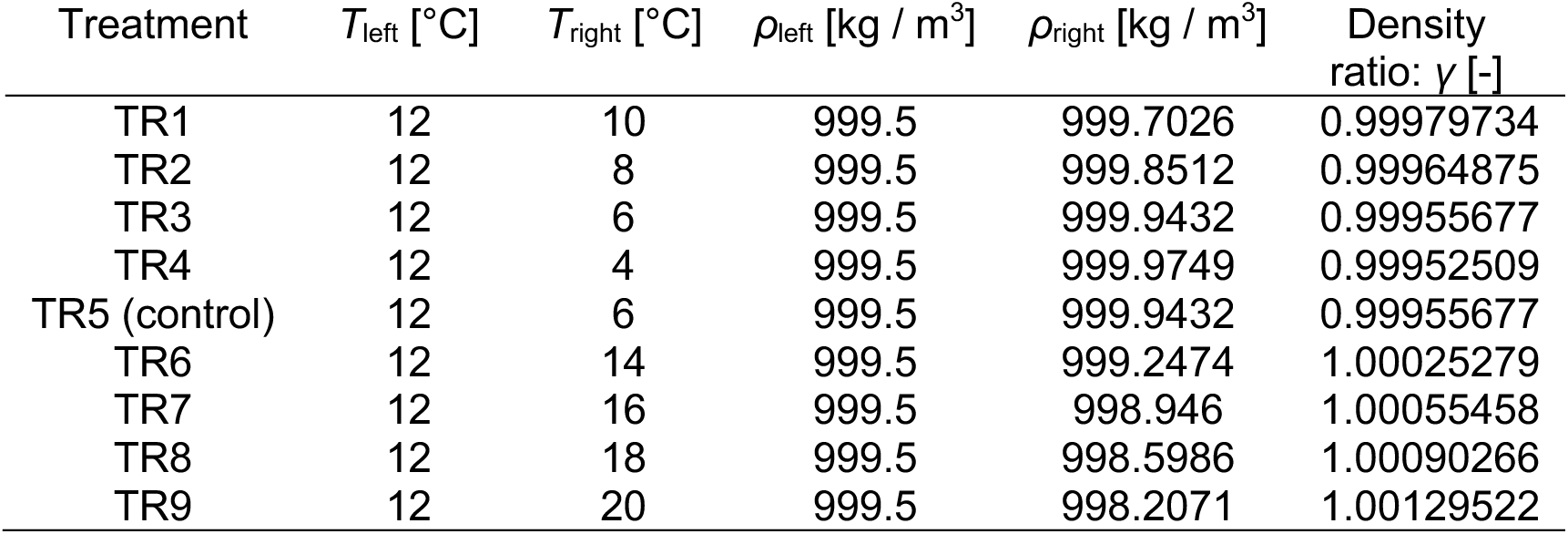
Definition of treatments. We contrasted water at the acclimation temperature (12°C) with four cold treatments (TR1-TR4) and four warm treatments (TR6-TR9) hence we varied the incoming water temperature in both directions (Table S1). Initial water temperature (*T*) and density (*ρ*) in the left (subscript: _left_) and right (subscript: _right_) compartments of the tank for each treatment. Treatment 5 (TR5) was the control treatment, conducted without the dye (see Supplementary Figs. 33 and 34).

**Table 2:**
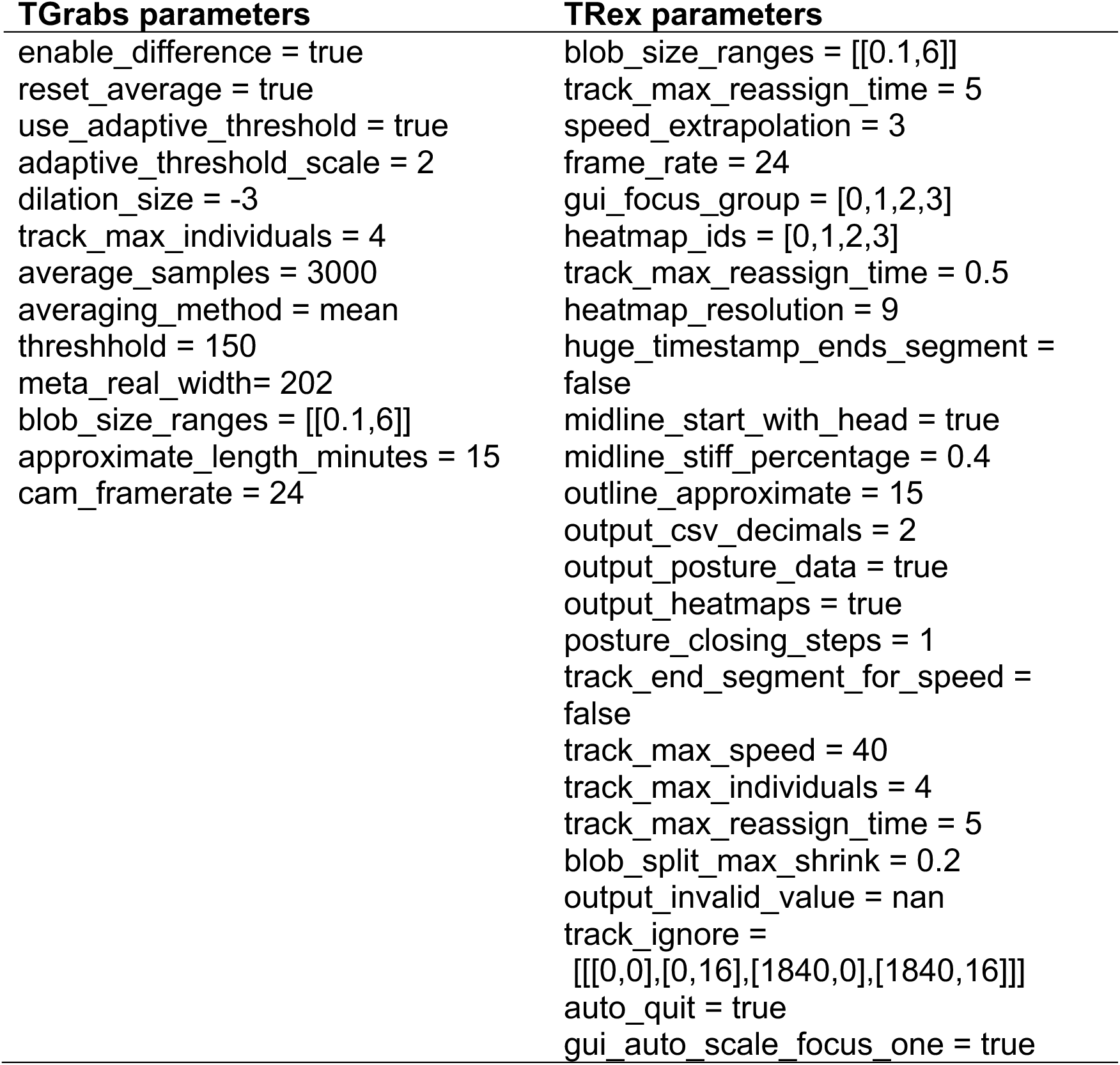
Tracking parameters used for fish tracking in TRex software. Detailed description of parameter can be found here: https://trex.run/docs/parameters_trex.html

**Table 3:**
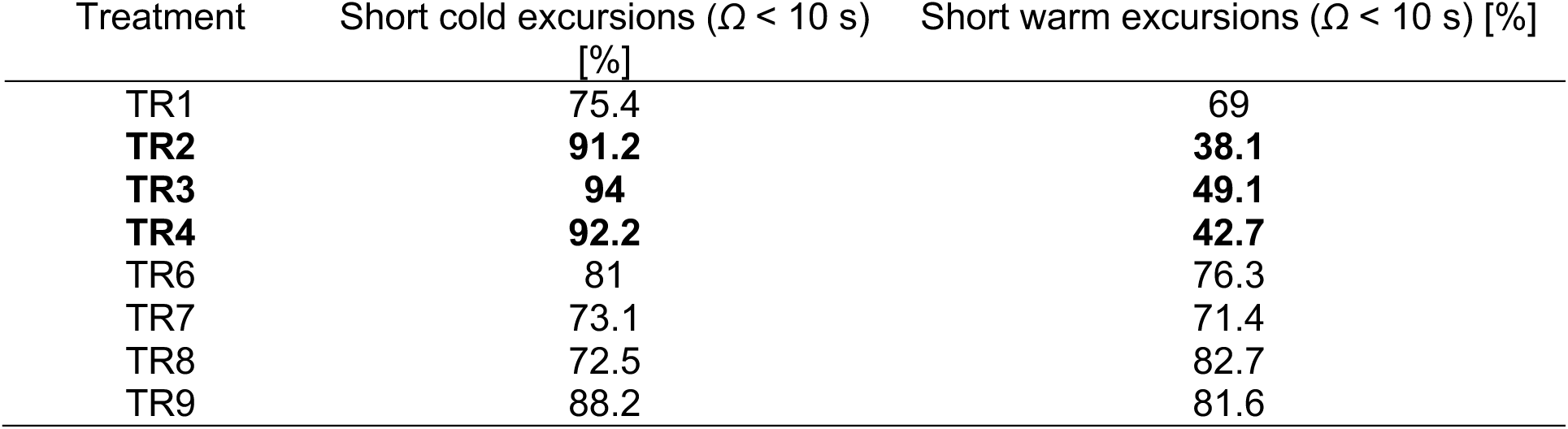
Percentage of all captured cold and warm excursions shorter than 10 seconds for all phases (p1, p2 and p3) of each treatment.

**Table 4:**
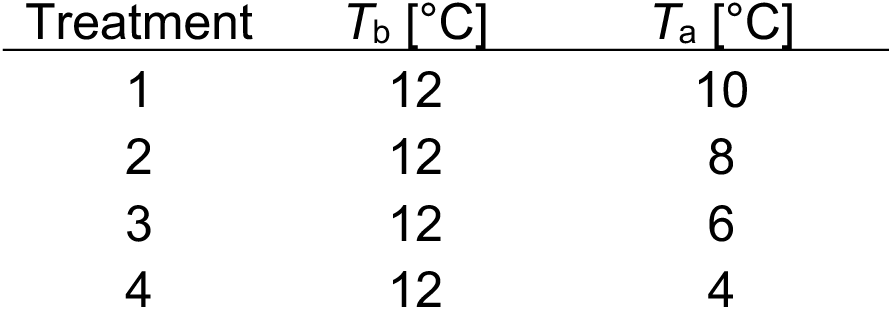
Input to body-cooling model. Summary of fish body temperature at beginning of cold-water excursion (*T*_b_) and ambient water temperature (*T*_a_) during cold-water excursions.

**Table 5:**
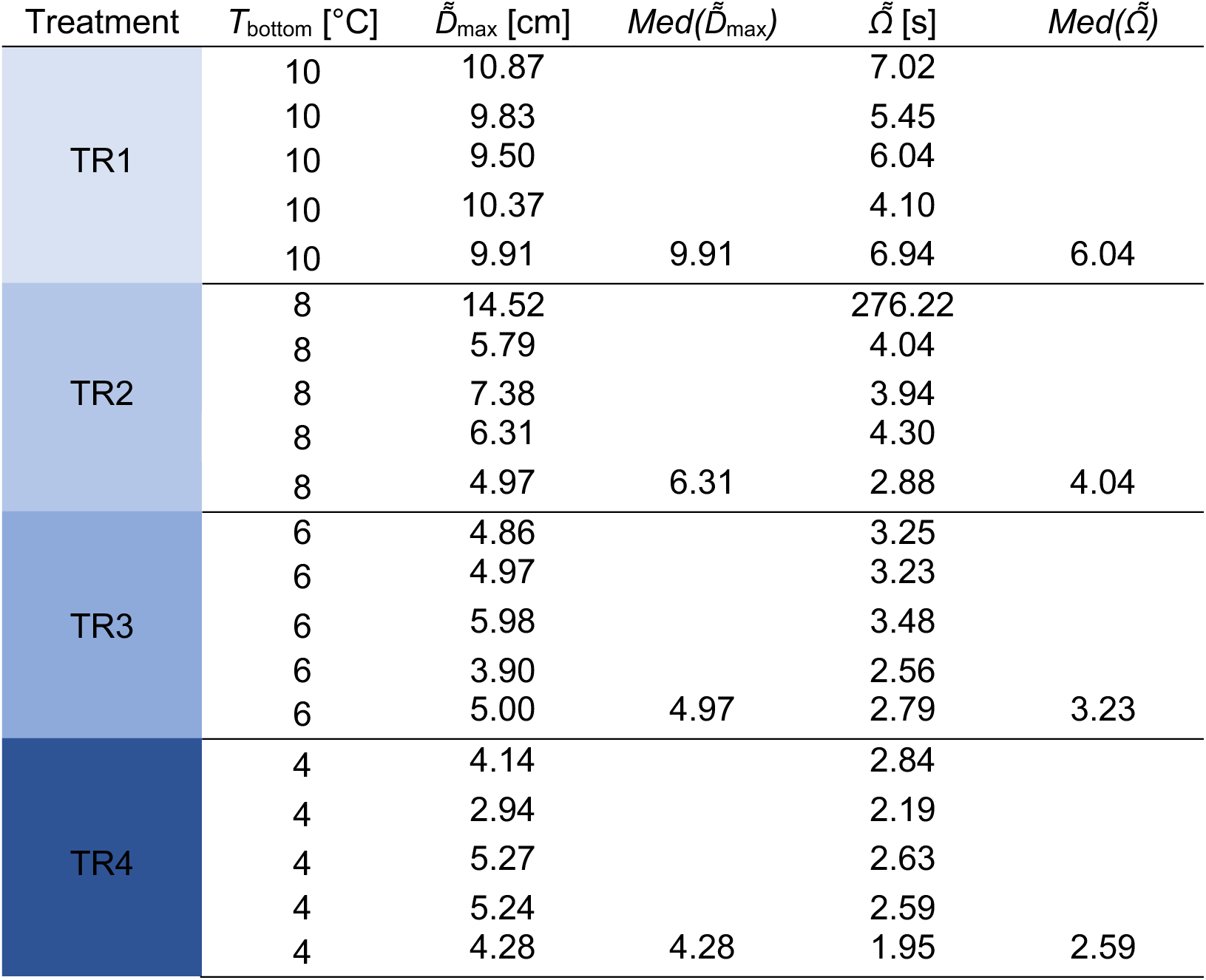
Mann-Kendall test. Input data: Median distance *D͌*_max_ (in centimeters) and median durations *Ω͌* (in seconds) across all treatments and experimental replicates.

**Table 6:**
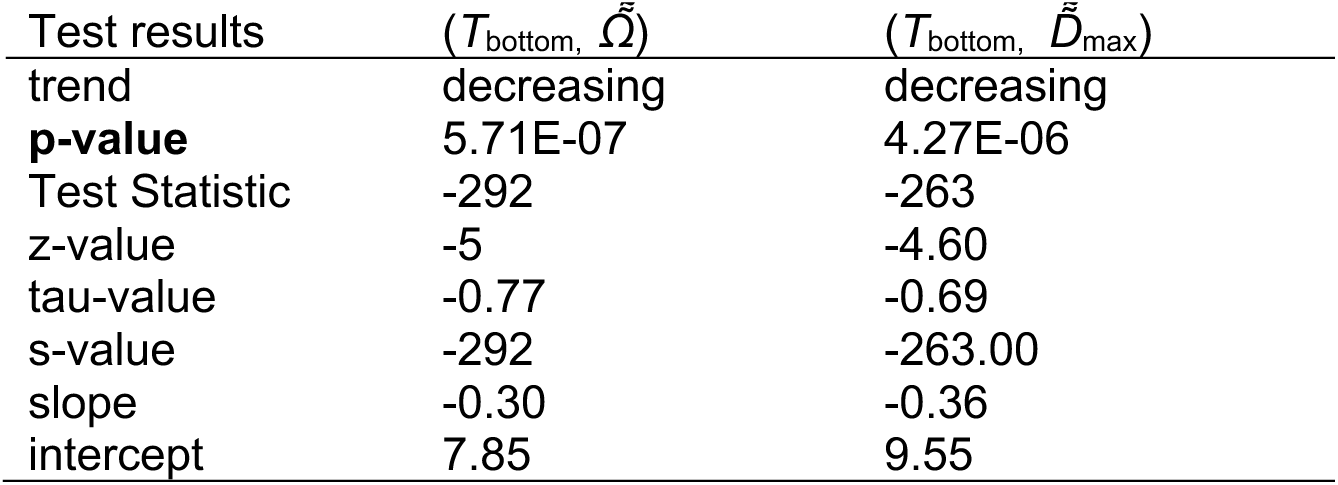
Mann-Kendall test result. A p-value lower than some significance level (common choices are 0.10, 0.05, and 0.01), indicates statistically significant evidence that a trend is present in the data. We found in both cases p < 0.001.

**Table 7:**
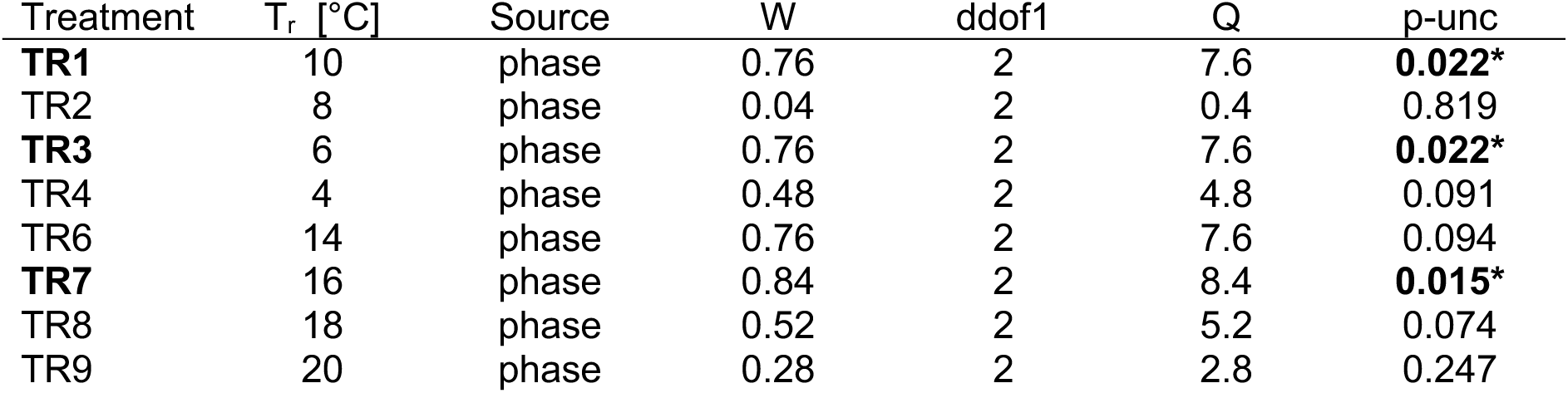
Friedman test within each treatment, < time-fraction spent in 12°C >, T_r_ depicts the treatment temperature initially in the right compartment of the tank.

**Table 8:**
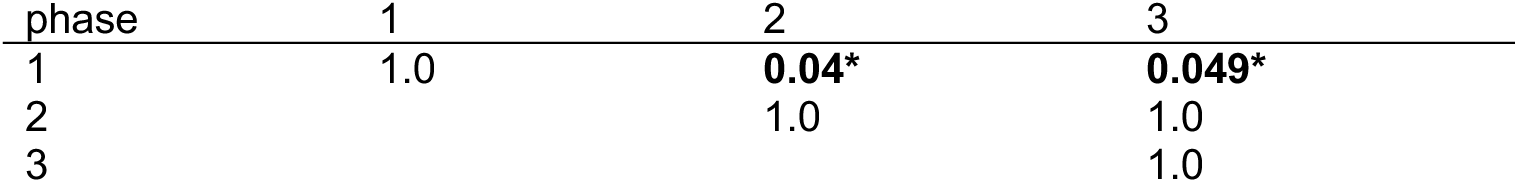
TR1, Dunn-test

**Table 9:**
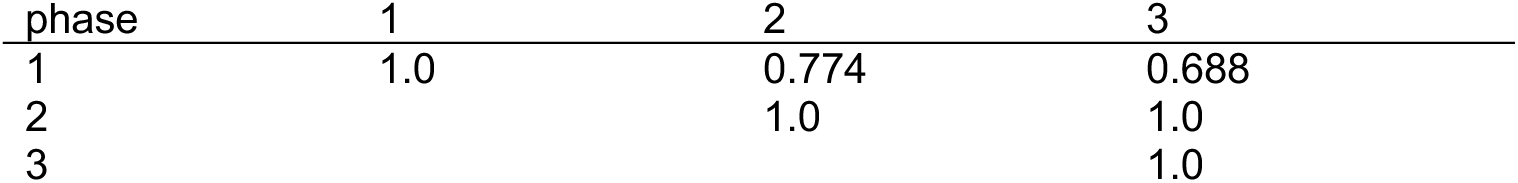
TR3, Dunn-test

**Table 10:**
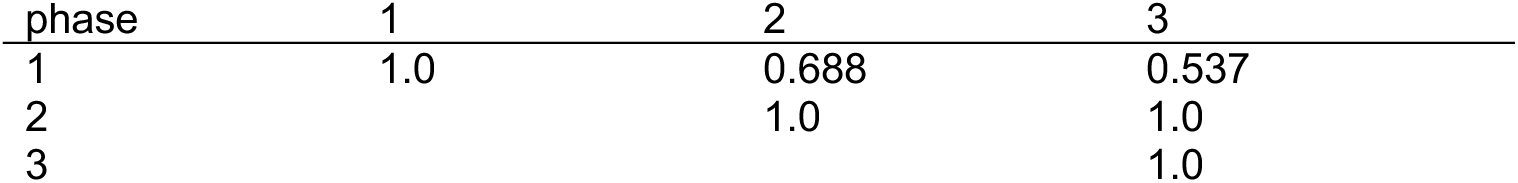
TR7, Dunn-test

**Number of cold-water excursions (Friedman and Dunn test)**

No significant reduction in the number of cold-water excursions were detected over time (see also *Supplementary* Fig. 15).

**Table 11:**
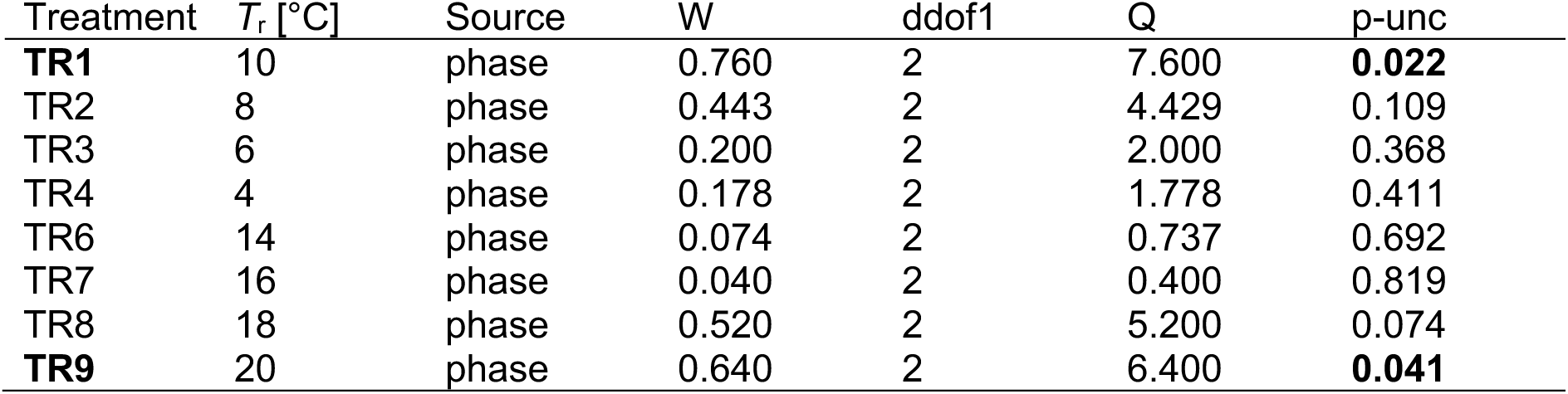
Friedman test, result, median number of excursions over time for all all-treatments.

**Table 12:**
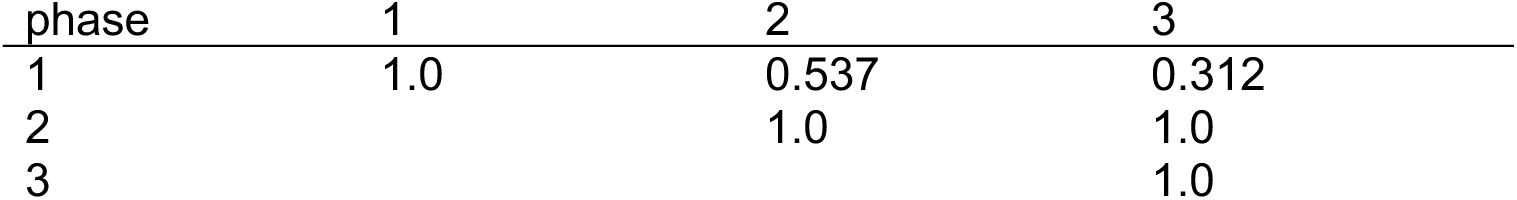
TR1, Dunn-test (post-hoc)

**Table 13:**
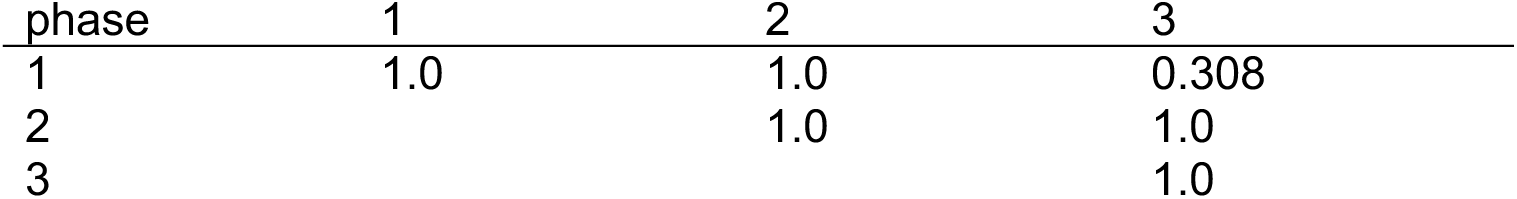
TR9, Dunn-test (post-hoc)

**Normed swimming speed (Shapiro-Wilk and t-test)**

Out of 20 fish, 2 fish in TR2 were basically ‘cold shocked’, hence they spent very little time in the warm. One of them only spent 2 seconds above the interface. For TR2 the data was thus not normally distributed (p > 0.05).

**Table 14:**
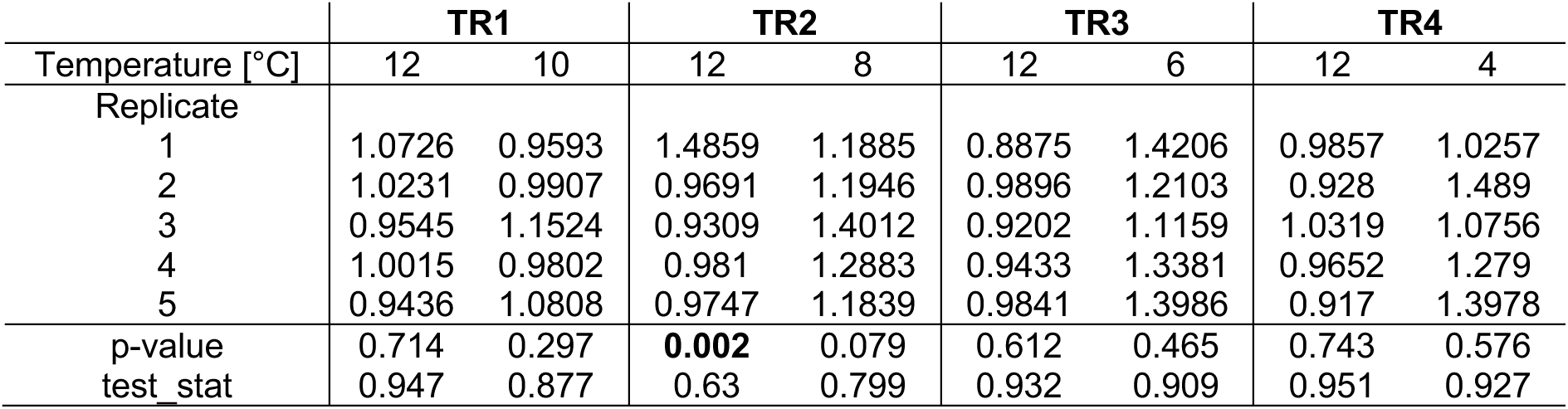
Shapiro-Wilk test for cold treatments. Group averaged and normed swimming speeds *v̅̅*_n,t_ for different temperatures (above and below interface).

**Table 15:**
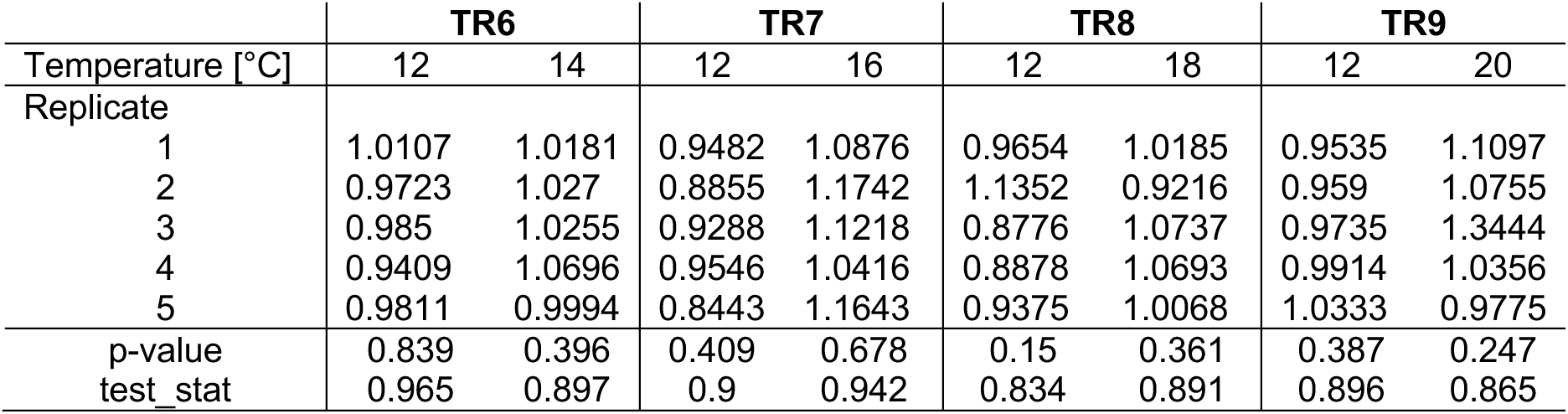
Shapiro-Wilk test for warm treatments. Group averaged and normed swimming speeds for different temperatures (above and below interface).

**Table 16:**
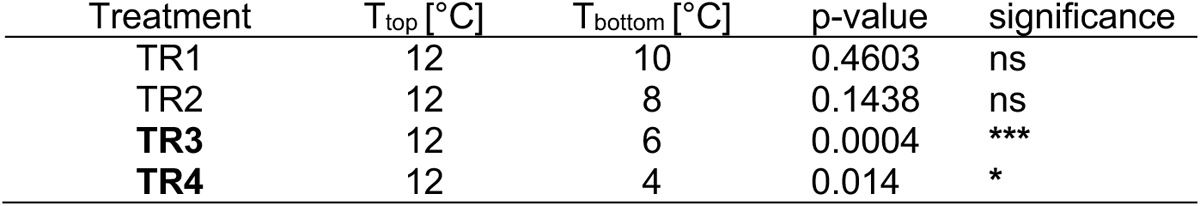
t-test results for cold treatments. ns: not significant; * : 0.01 < p ≤ 0.05; *** : 0.0001 < p ≤ 0.001

**Table 17:**
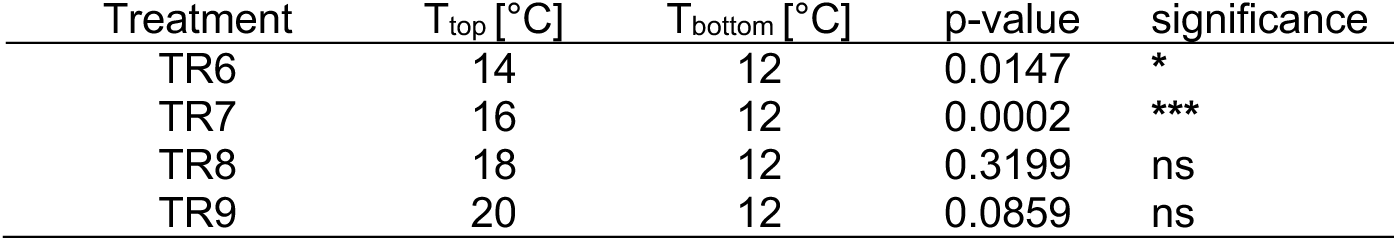
t-test results for warm treatments.

**Number of cold-water excursions (Shapiro-Wilk, ANOVA, t-test)**

To test whether the median number of cold-water excursions *Ñ* is normally distributed, we conducted a Shapiro-Wilk test for each of our treatments. The data of each treatment was confirmed to be normally distributed (Table S18) (i.e. p-value > 0.05) and thus, a one-way ANOVA (python: scipy.stats.f_oneway) was applied to test for significant difference between the treatments (Table S19). To find specific differences, we then applied a t-test to see which treatments are significantly different (Table 20).

**Table 18:**
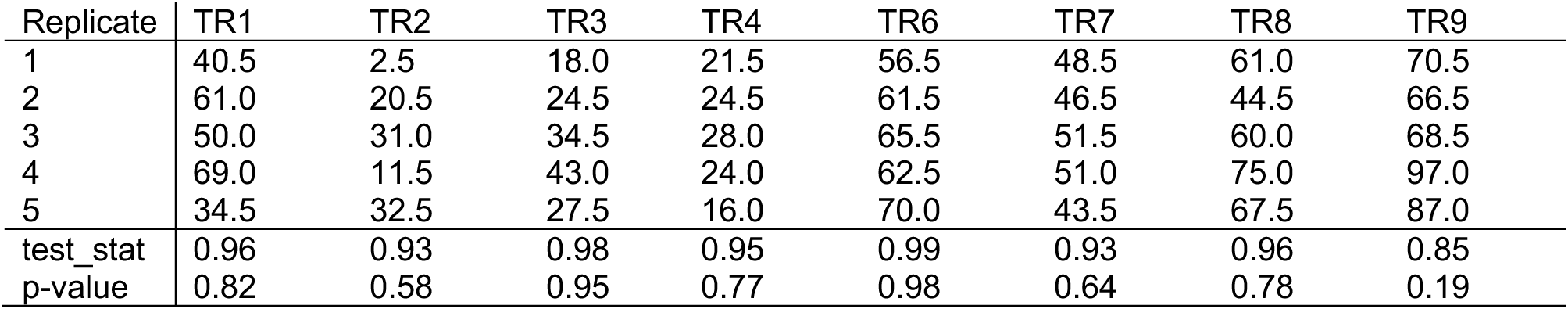
Median number of cold-water excursions *Ñ* of 4 fish tested simultaneously within each experimental replicate. Shapiro-Wilk-test results.

**Table 19:**
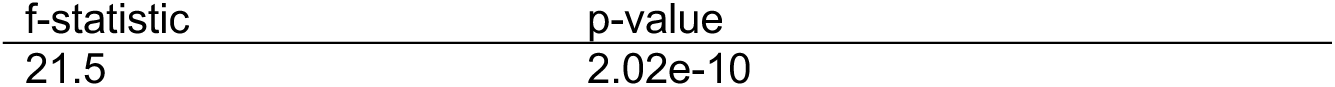
One-way ANOVA test result

**Table 20:**
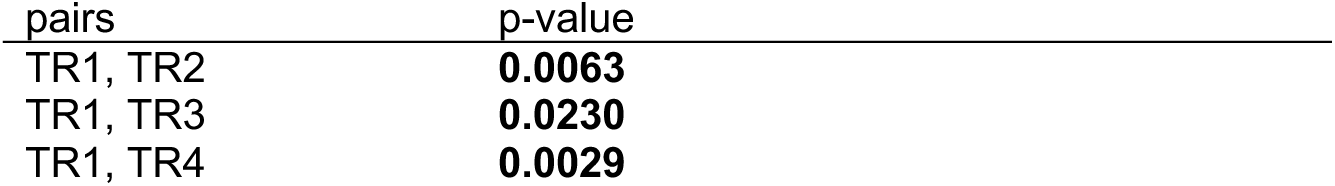
Pairwise t-test result

## Supplementary Text

### Text 1: Gravity currents

The difference in specific weight between warmer and colder fluid provided the driving force for the gravity currents that unfolded during the first 2 min after gate removal. After impounding at the wall, the thermal interface separated the cold water in the bottom from the warmer water in the top of the tank across the entire length *L* = 2 m of the tank. However, the thermal interface was not stationary in time. Continuously dissipating the initial energy of the system, the interface was swaying between left and right side of the tank with diminishing movement over time (Fig. 2 and Fig**. 3**). For all treatments, its vertical position varied more strongly across *X* and in time during p1 and decreased for p2 and p3. The maximum movement velocity was small (*v*_interface_ ≈ 0.1 cm/s) when compared to the typical swimming speed of the tested fish (*v*_Fish_ ≈ 2 - 8 cm/s, Fig. 20). The effect of the flow-field was therefore neglected.

### Text 2: The thermal interface

The lateral view of the tank was recorded at 24 frames per second with a digital video camera (GoProHero7). The light was absorbed by the dye and hence, the intensity of the image could be related to the dye concentration along the light path from the back to the front of the tank ^66^. With this arrangement the reduction in intensity is exponentially related to the dye concentration. Since the blue dye diffuses at about the same rate as (in both cases diffusion is negligible for this flow), the dye concentration is a surrogate for the density and therefore the water Temperature.

Our setup allowed to investigate fish behavior in an isolated fashion following a reductionist approach where non-thermal stimuli were either absent (prey, predators, chemical gradients) or kept constant throughout the water tank, in time and across treatments (illumination, dissolved oxygen, dissolved ink).

On the left side, fish (tested in a group of four in each replicate) were acclimated (15 min) to water at 12 °C, which was identical to the housing temperature. On the right side, we added water at a different temperature (colder or warmer), dyed with 0.29 mg/l pre-dissolved *Patent blue V sodium salt* (C_27_H_31_N_2_NaO_7_S_2_, Sigma-Aldrich) to allow visualization of the thermal gradient.

### Text 3: Trajectory smoothing

The fishs location at any time instance is represented by the center of gravity of its detected shape. Naturally this location is displaced by short-term body rotations or body wobble leading to unwanted artefacts in the trajectory which propagate into its derivatives (i.e velocity and acceleration). To dampen these high frequency fluctuations, we fitted a ‘moving-spline’ to the trajectory acting as a low pass filter (Supplementary Fig. 38 B,C and *D*) – an approach suggested by ^76^. For each time step, t, and around each point of a measured trajectory a third order polynomial is fitted from t − 10 · Δt to t +10· Δt for each component, *x*_1_, *x*_2_ (*x*- and *y*-position). This results in a fit to 21 measured trajectory points. The constants, *c*_i_ , for the polynomial of type

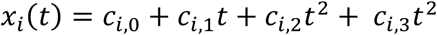

are determined as

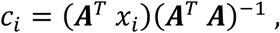

where

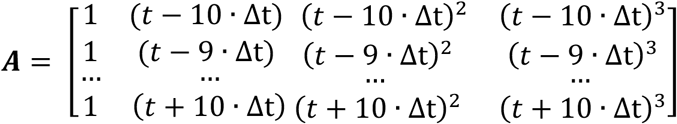

and

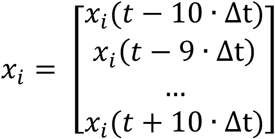

The filtered position, velocity and acceleration, 𝑥’_i_ (𝑡), 𝑢’_i_(𝑡) 𝑎𝑛𝑑 𝑎d_i_(𝑡), where then obtained per component as:

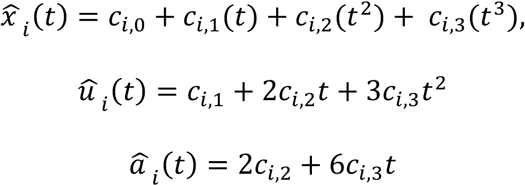

the swimming speed 𝑣’(𝑡) was then derived as:

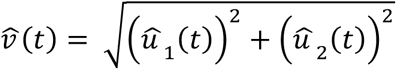

Finally, an individuals’ normalized swimming speed 𝑣_789:_(𝑡) is calculated by division through the individuals’ temporally averaged speed during the treatment phase, 𝑣^g^’:

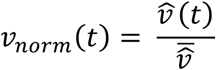

The effect of the moving spline on the raw velocity components and the derivation of 𝑣’(𝑡) and 𝑣_789:_(𝑡) is visualized in Supplementary Fig. 38 A-E and in Movie 5.

#### Text 4: Burst swimming model

We compared our observed maximum swimming speeds to a multivariate swimming speed model for rheophilic fish species ^77^. Thereby fish burst swimming speed (*u*) was calculated for a duration of *t* = 5 seconds, body length of *BL* = 0.036 m and temperatures of *T* = 12 °C and *T* = 6 °C (Supplementary Fig. 21).

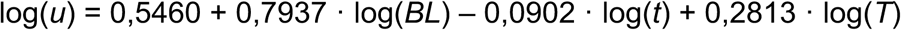

The direct comparison of our observed absolute swimming speeds to the model is however difficult because:

1. The model is based on swimming speed derived from respirometers where a steady flow is present. The absence of flow in our experiments does not compare to this and fish inherently displays lower swimming speeds.
2. The confined space and the presence of walls forces fish to decelerate frequently, preventing it from reaching very high swimming speeds.
3. We are looking at the 2D-projection of the swimming trajectory, thus our observations neglect the 3^rd^ velocity component.

## Supplementary Movies

All movies are played at 20x real speed. Original framerate was 24 frames per second.

**Movie 1: Tracking of 1 fish in cold treatment (TR2).**

Side-view of the tank with individual fish swimming-trajectory and gravity current for single experimental replicate (TR2, cold-water in light blue). Individual fish movement trajectory is shown in red and its instantaneous vertical distance to the thermal interface *D(t)* in the lower plot for the progression of the entire treatment (p1, p2 and p3) immediately after gate-removal. Playback is 20x real speed.

**Movie 2: Tracking of 4 fish in cold treatment (TR2).**

Same for all 4 individuals.

**Movie 3: Tracking of 1 fish in warm treatment (TR7)**

Side-view of the tank with individual fish swimming-trajectory and gravity current for single experimental replicate (TR7, warmer water in light blue). Individual fish movement trajectory is shown in red and its instantaneous vertical distance to the thermal interface *D(t)* in the lower plot for the progression of the entire treatment (p1, p2 and p3) immediately after gate-removal. Playback is 20x real speed.

**Movie 4: Tracking of 4 fish in warm treatment (TR7).**

Same for all 4 individuals.

**Movie 5: Derivation of normed swimming speed**

Derivation of normed swimming speed for trajectory segment of 60 seconds. (*A*) Single fish trajectory. (*B*) and (*C*) horizontal and vertical velocity components. (*D*) Absolute swimming speed. (*E*) Normed swimming speed.

