## Supplementary revised for "Salmonids elicit an acute behavioral response to heterothermal environments"

### Supplementary information for
Salmonids elicit an acute behavioral response to heterothermal environments

Robert Naudascher[a,*], Stefano Brizzolara[a,c], Jonasz Slomka[a], Robert M. Boes[b], Markus Holzner[a,c], Luiz G. M. Silva[a] and Roman Stocker[a,*]

(a) Institute of Environmental Engineering, Department of Civil, Enivironmental and Geomatic Engineering, Swiss Federal Institute of Technology Zürich, 8093 Zürich, Switzerland
(b) Laboratory of Hydraulics, Hydrology and Glaciology (VAW), Swiss Federal Institute of Technology, 8093 Zürich, Switzerland
(c) Swiss Federal Institute of Forest, Snow and Landscape Research WSL, 8903 Birmensdorf, Switzerland

**Supplementary information for this manuscript include the following:**

Supplementary Figures 1–38
Supplementary Tables 1–20
Supplementary Text 1-4
Legends for Supplementary Movies 1-5
Supplementary References

#### Supplementary Figures

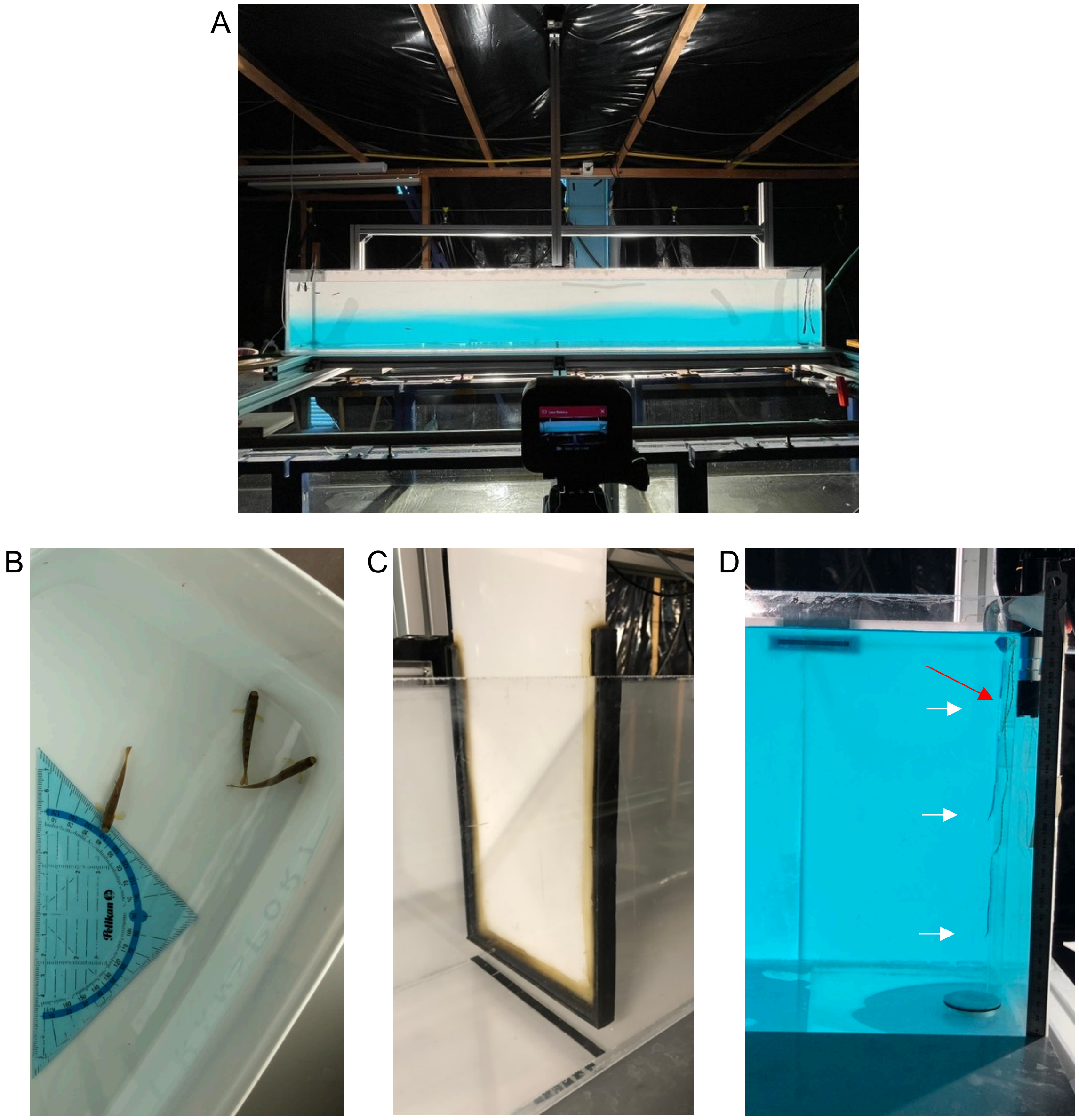

**Fig. 1:** (*A*) Photograph showing a side-view of the experimental tank in phase 2 (*approximately 10 min* after removal of the central gate) with four individual fish. The camera used for imaging is visible in the foreground. The dark casing obstructs the entry of natural light. Experimental animals (*B*); central gate to partition the tank; (*C*) right side of the tank with 3 temperature probes (white arrows) and dissolved oxygen probe (red arrow) (*D*).

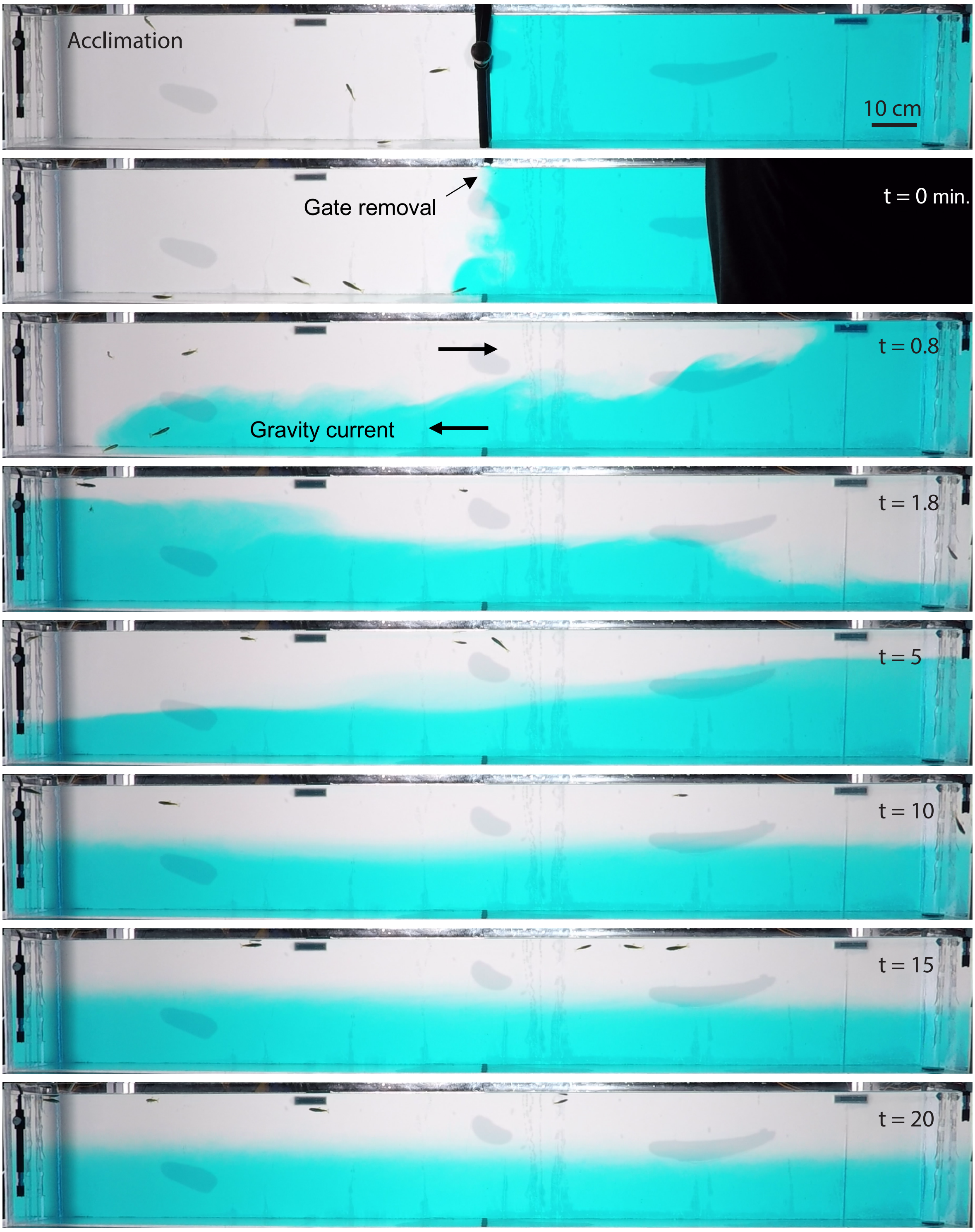


**Fig. 2:** Side view of temporal progression of gravity current for cold treatment. During the acclimation phase (top panel) the water tank is partitioned into 12° C in the left compartment and 4°C (TR4) in the right compartment. After 15 minutes of acclimation the central gate is removed (*here at time:* t = 0) and the density difference drives a lock exchange flow with the transparent warm water flowing *slowly* in the upper half of the tank to the right and the dyed cold water flowing along the channel bottom towards the left (t = 0.8 minutes). After reaching the side walls of the tank, the directionality of the flow reverts and the current sloshes back and forth in the form of an internal wave (t = 1.8 – 5 minutes). A clear second and a third wave passage take place, while flow speed and wave amplitude gradually attenuate through dissipation. During these internal wave passages, the vertical height of the temperature interface remains a periodic function of time. A stationary vertical stratification (i.e. the water is practically still) is reached only after about t = 10 minutes. The start of the first treatment phase (p1) was defined as the time when the vertical gate was fully removed from the water (t = 0).

#### Temporal movement of the interface

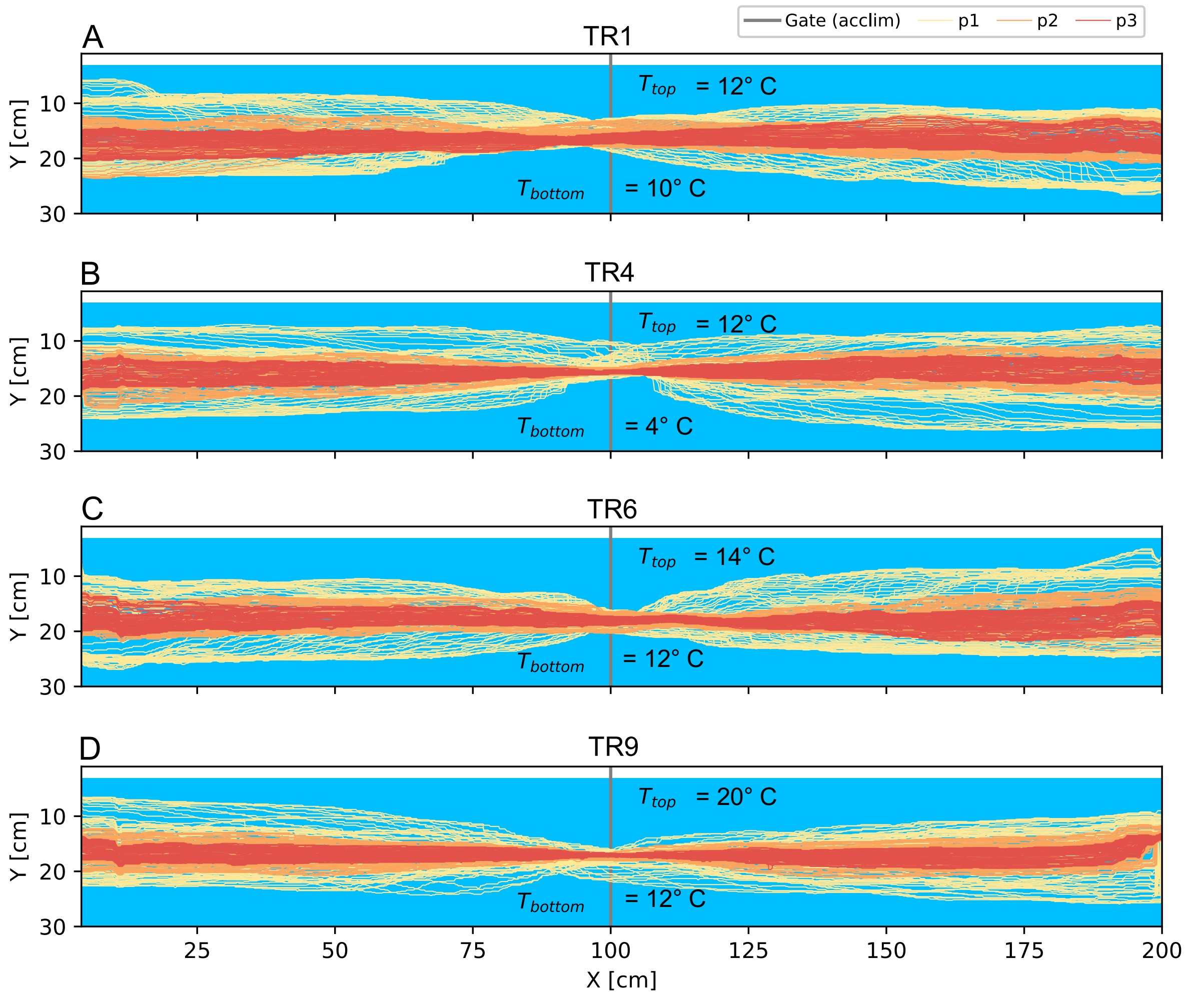


**Fig. 3**: Transient movement of the thermal interface/line during experimental phases (p1, p2 and p3). Coloured lines depict time steps of 10 seconds. The water surface is located at *Y* = 4 cm. The instantaneous position of the thermal interface is shown for each experimental phase (acclim., p1, p2, and p3) at a reduced temporal resolution of 0.2 Hz (the temporal resolution in the experiments was 24 Hz). The interface was approximated as a line (X,Y) for each frame. During the acclimation period (acclim: grey dots) a sealed gate was located at X = 100 cm. The interface is coloured by the respective temporal phase of the experiment (see *A*).

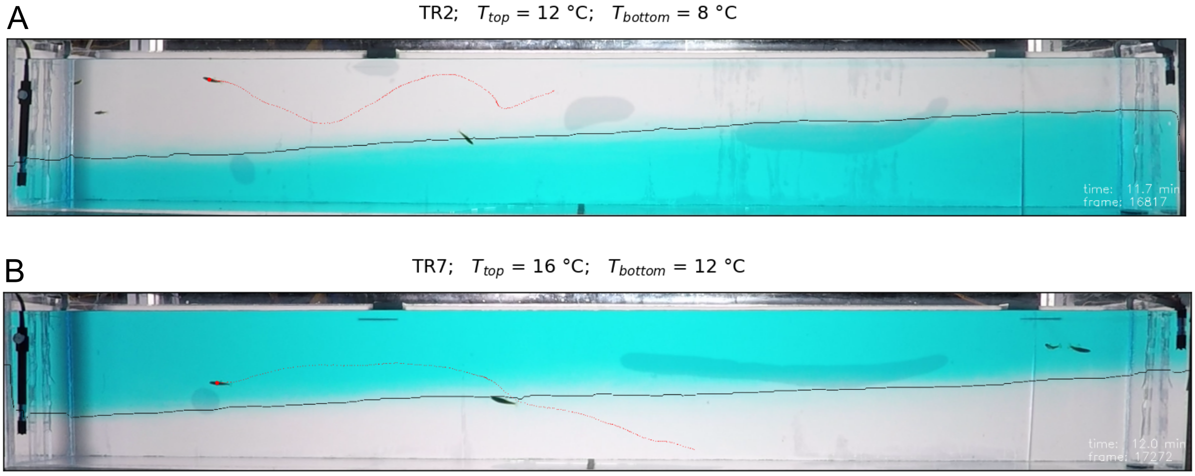


**Fig. 4:** Side view of experimental setup for cold (*A*) and warm (*B*) treatment (TR2 and TR7). The thermal interface line (black) was derived using custom made image analysis pipeline.

#### Vertical occupation

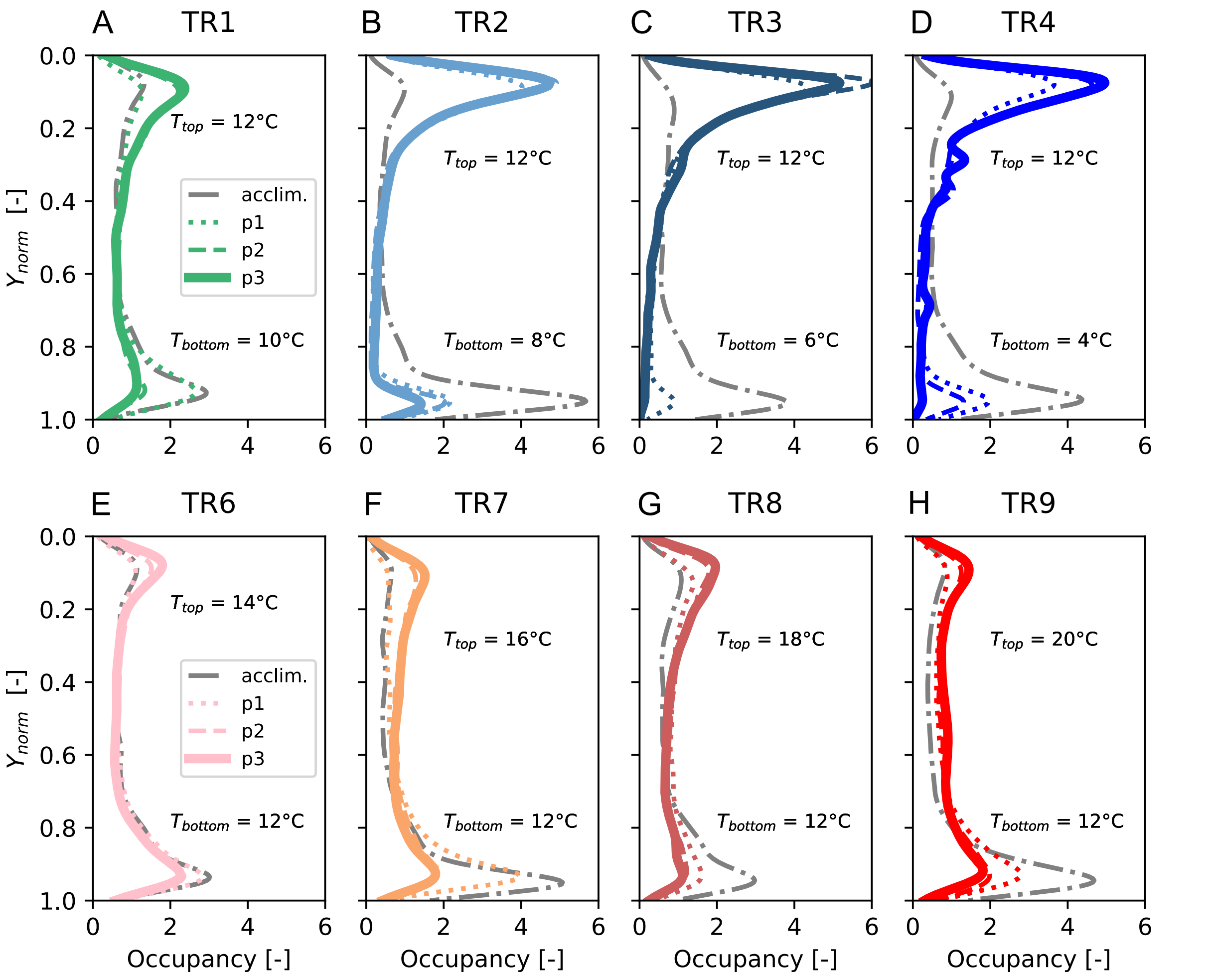


**Fig. 5:** Probability density functions of fish depth for all cold-water (TR1–TR4, *A*–*D*) and all warm-water (TR6–TR9, *E*–*H*) treatments ($N$ = 20 fish in each case), separated according to the four phases of analysis (acclimation, and the three experimental phases p1–p3). The data was aggregated at the treatment level; hence each line represents data from 20 fish, tested in groups of 4 individuals in 5 separate experiments. $T_{\text{bottom}}$ and $T_{\text{top}}$ indicate the water temperature below and above the thermal interface. The depth was normalized so that $Y_{\text{norm}}$ = 0 is the water surface and $Y_{\text{norm}}$ = 1 indicates the bottom of the tank.

#### Temporal progression of excursions (cold-treatments)

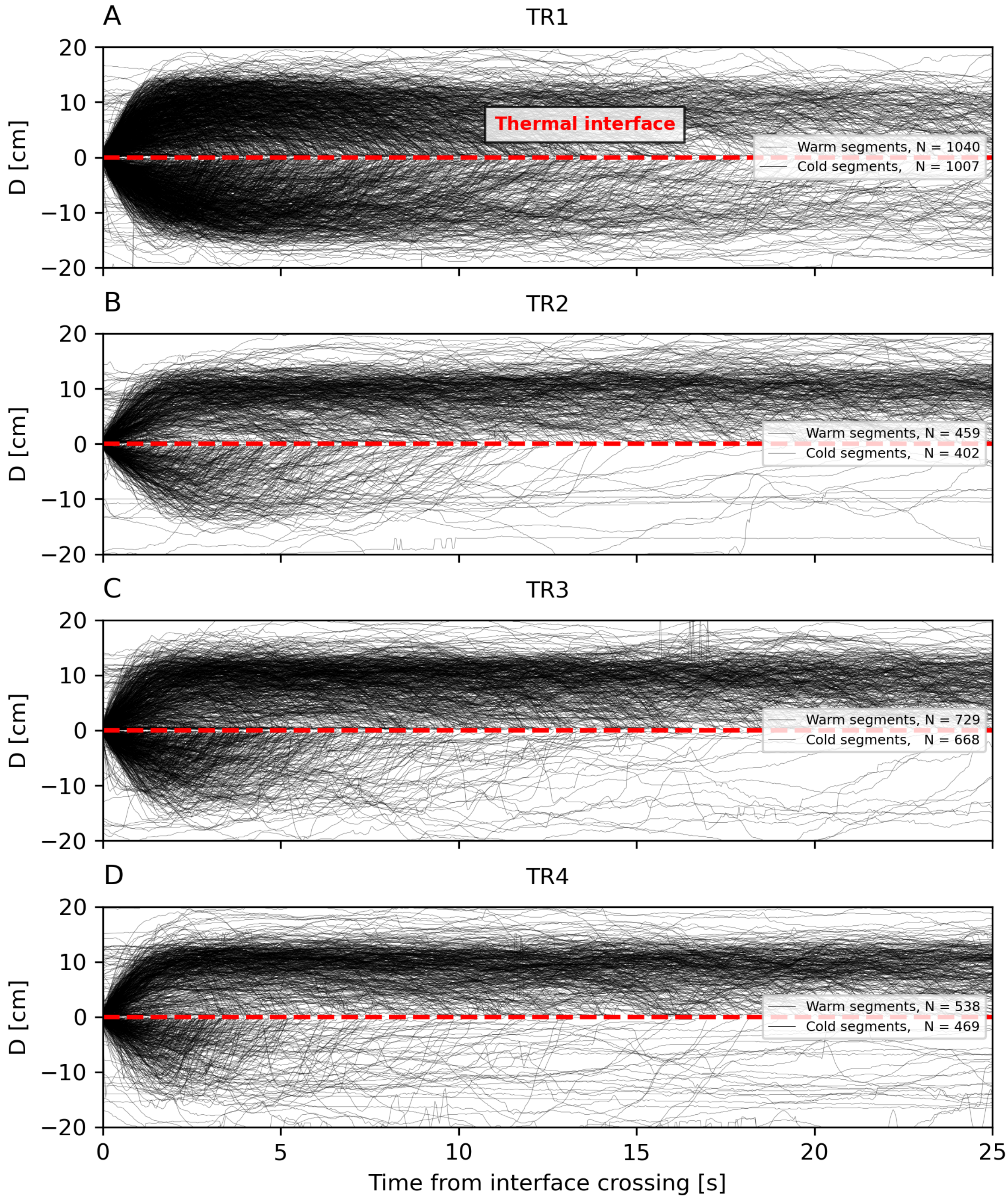


**Fig. 6: Cold treatments, TR1-TR4.** Temporal evolution of the fish's instantaneous vertical distance to the thermal interface D(t) within the first 25 seconds for each cold- and warm-water interval for all cold treatments. At t = 0 fish crosses the thermal interface line upwards (warm segment; upper half) or downwards (cold segment; lower half). The end of an interval occurs when the interface line was crossed again in the opposite direction.

#### Temporal progression of excursions (warm-treatments)

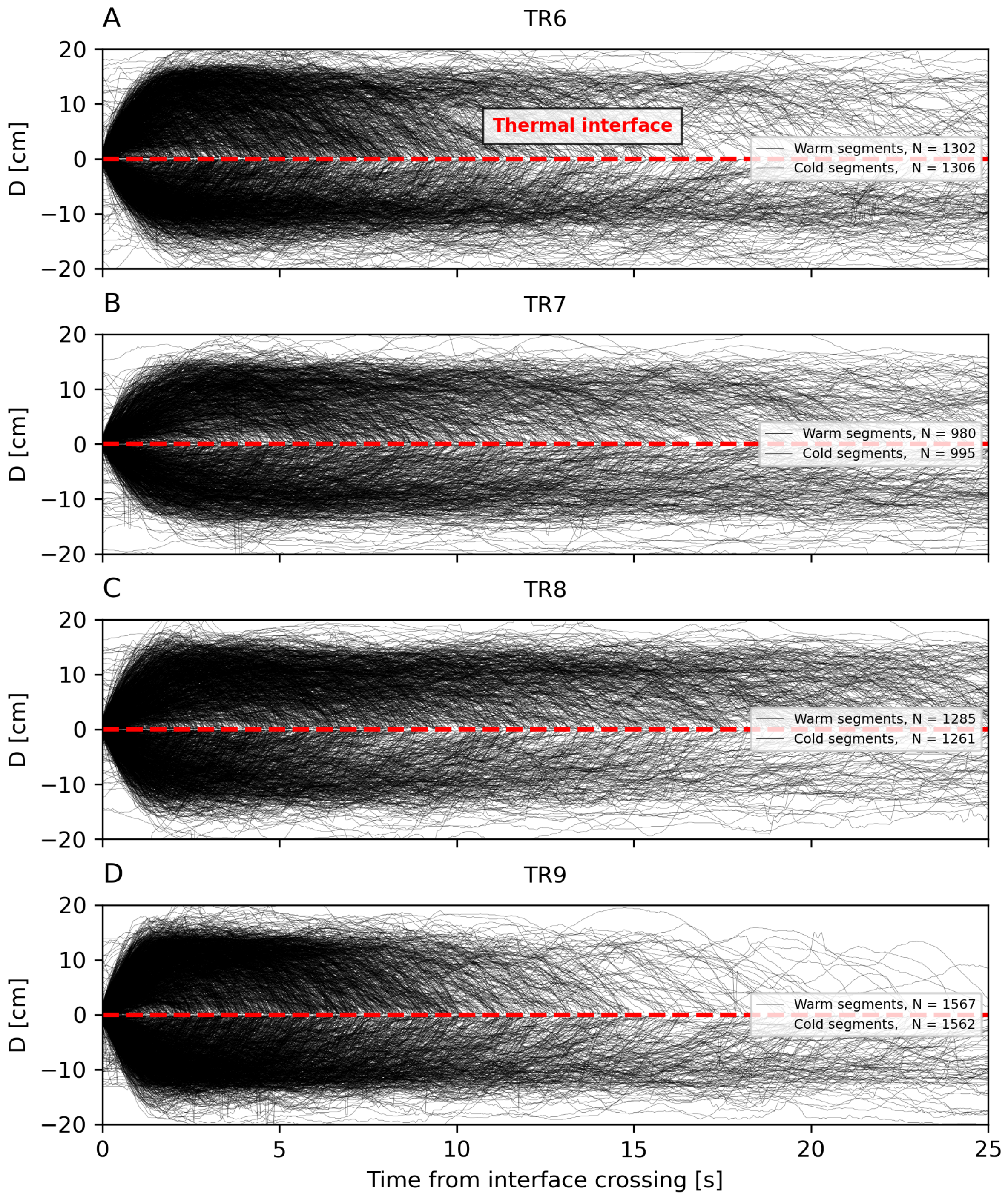


**Fig. 7: Warm treatments TR6-TR9.** Temporal evolution of the fish's instantaneous vertical distance to the thermal interface D(t) within the first 25 seconds for each cold- and warm-water interval for all warm treatments. At t = 0 fish crosses the thermal interface line upwards (upper half) or downwards (lower half). The end of an interval occurs when the interface line was crossed again in the opposite direction.

#### Time fraction in acclimation water

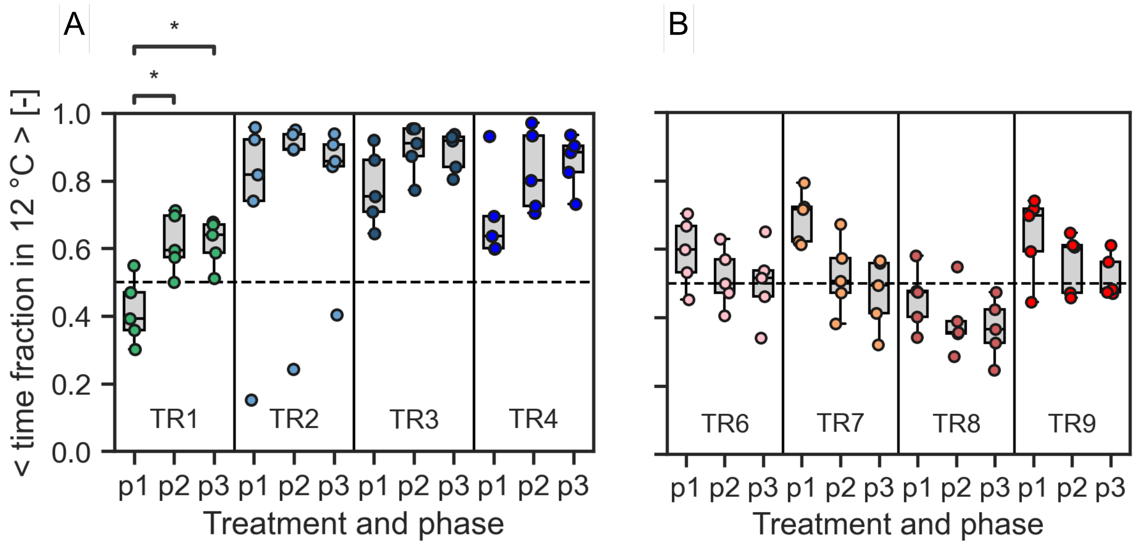


**Fig. 8:** The average time fraction spent in 12 °C was calculated for each fish and phase (p1, p2 and p3) as $T_{cold}$ / 6 minutes for incoming cold water (TR1-TR4, *A*) and incoming warm water (TR6-TR9, *B*). Each dot represents the average of this quantity over four individuals that were tested simultaneously within one experimental replicate. The Friedman-test suggests that there are significant temporal differences within TR1 (p = 0.022). Significance levels are indicated above the plot (Dunn-test: * : 0.01 < p ≤ 0.05).

#### Vertical occupation of the tank

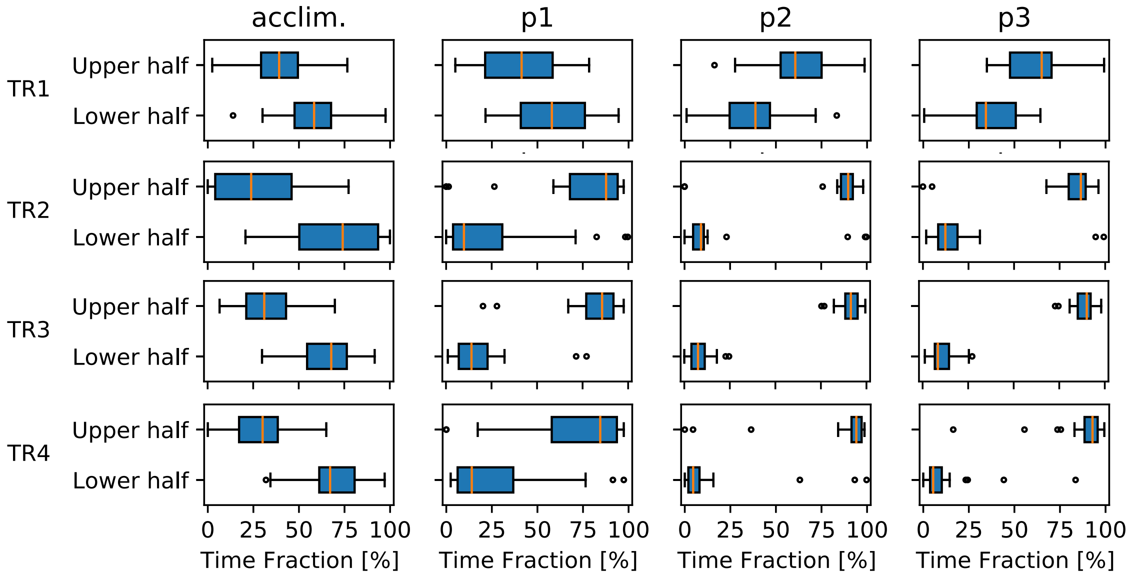


**Fig. 9:** Cold treatments. Boxplots of time fraction spent in upper half ($Y_{norm}$ = 0–0.5) and lower half ($Y_{norm}$ = 0.5–1) of the tank across experimental phases (acclim., p1, p2 and p3) and treatments (TR1, TR2, TR3 and TR4).

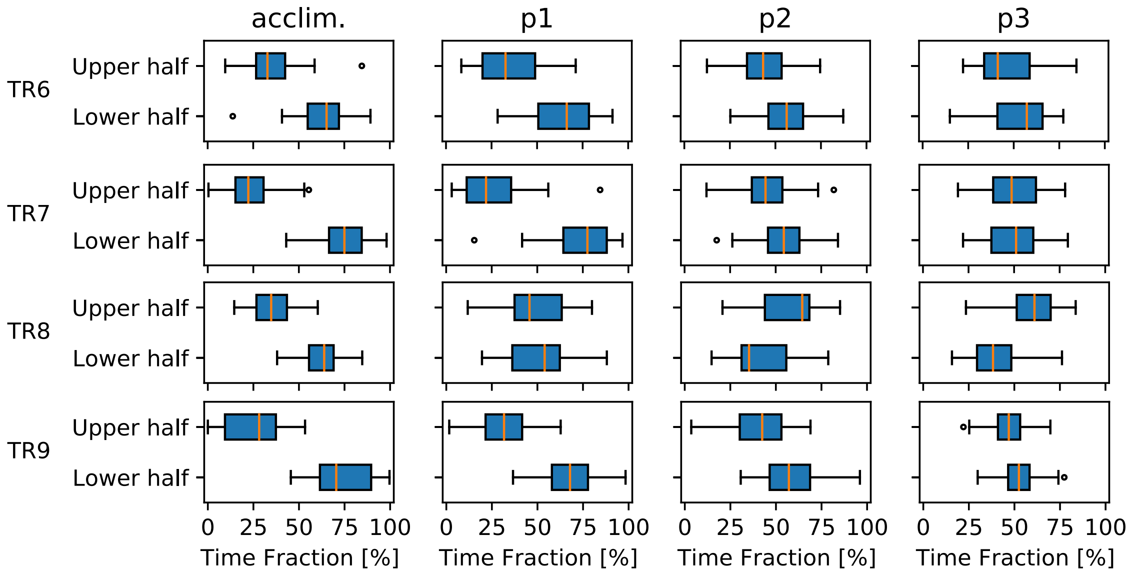


**Fig. 10:** Warm treatments. Boxplots of time fraction spent in upper half ($Y_{norm}$ = 0–0.5) and lower half ($Y_{norm}$ = 0.5–1) of the tank across experimental phases (acclim., p1, p2 and p3) and treatments (TR6, TR7, TR8 and TR9).

#### Detection of vertical turning points

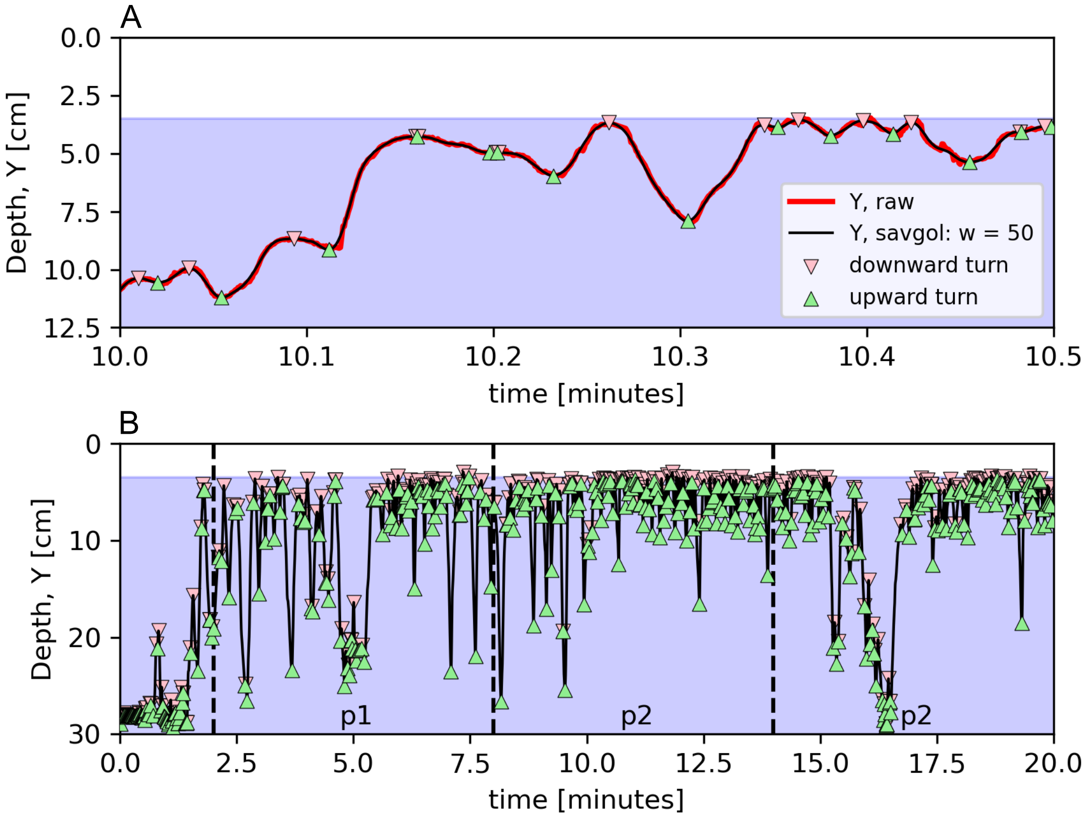


**Fig. 11:** Derivation of vertical turning points. (*A*) To smoothen the fish's vertical position signal (red line), a fifth order polynomial was fitted to a moving window of 50 consecutive time points (black line). This is a standard practice in particle tracking to filter out noise (e.g. Lüthi et al. JFM 2005). Therefore the SAVGOL function (python: scipy.signal.savgol_filter) was applied on the raw output (black line) from the tracking software. Upward turning (green triangles) were detected as sign change (from negative to positive) of the derivative of the smoothed signal. Downward turning (red triangles) were detected via sign change (from positive to negative). (*B*) Occurrence of turning points after gate removal (at t = 0) for an individual fish in cold treatment (TR3).

#### Large turns cold treatments (TR1 – TR4)

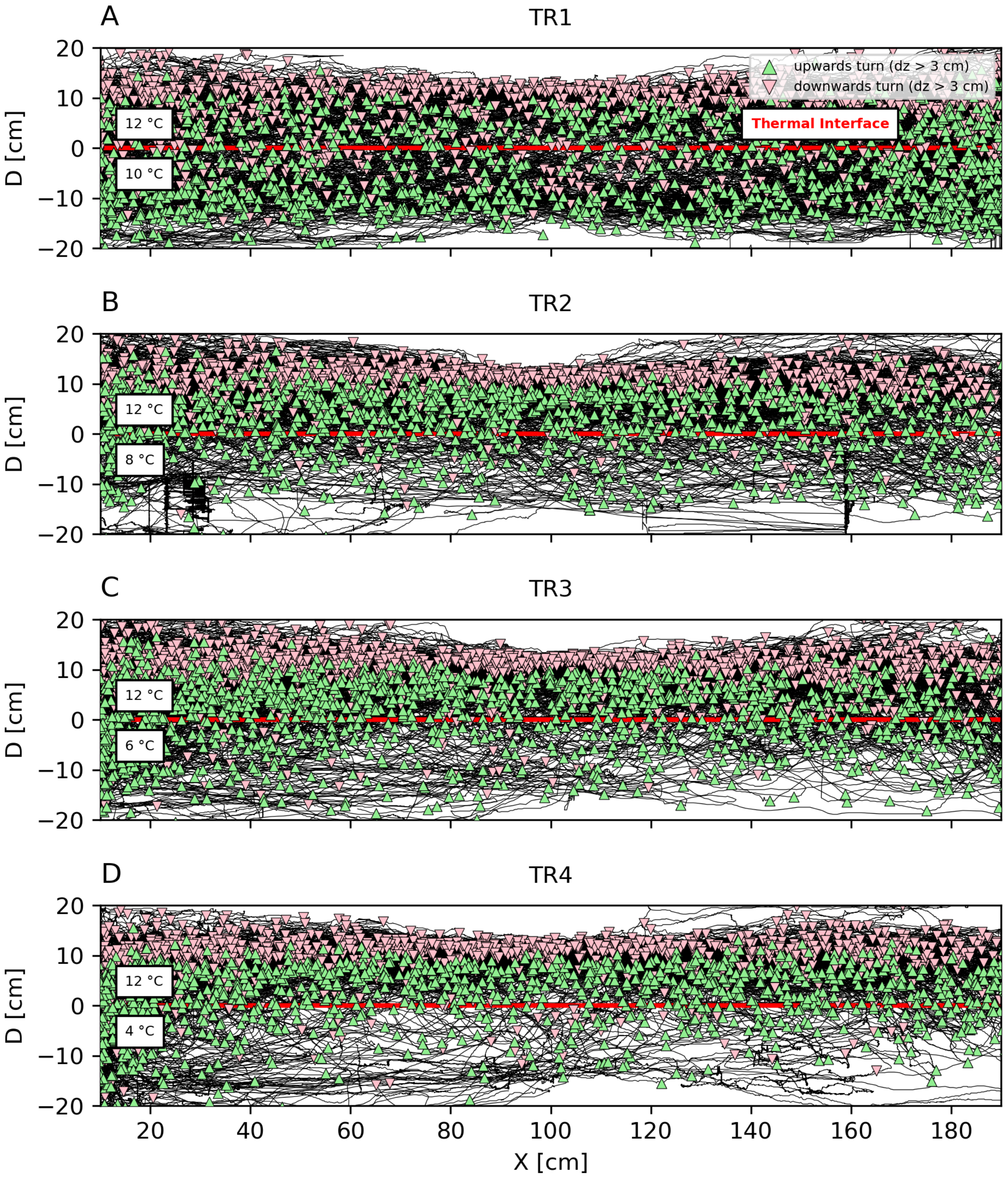


**Fig. 12:** Cold treatments. Swimming trajectories (black lines) relative to the thermal interface (red line) for all fish across cold treatments (TR1 – TR4, *A-D*). The horizontal axis depicts the horizontal position in centimetres; the vertical axis depicts the instantaneous distance from the thermal interface. For D > 0 fish is located at a distance D [cm] above the interface and for D < 0 the fish is located below the temperature interface. Red and green triangles depict large downward and upward turns, respectively (see Supplementary Fig. 11 and Fig. **14** for derivation).

**Large turns warm treatments (TR6 – TR9)**

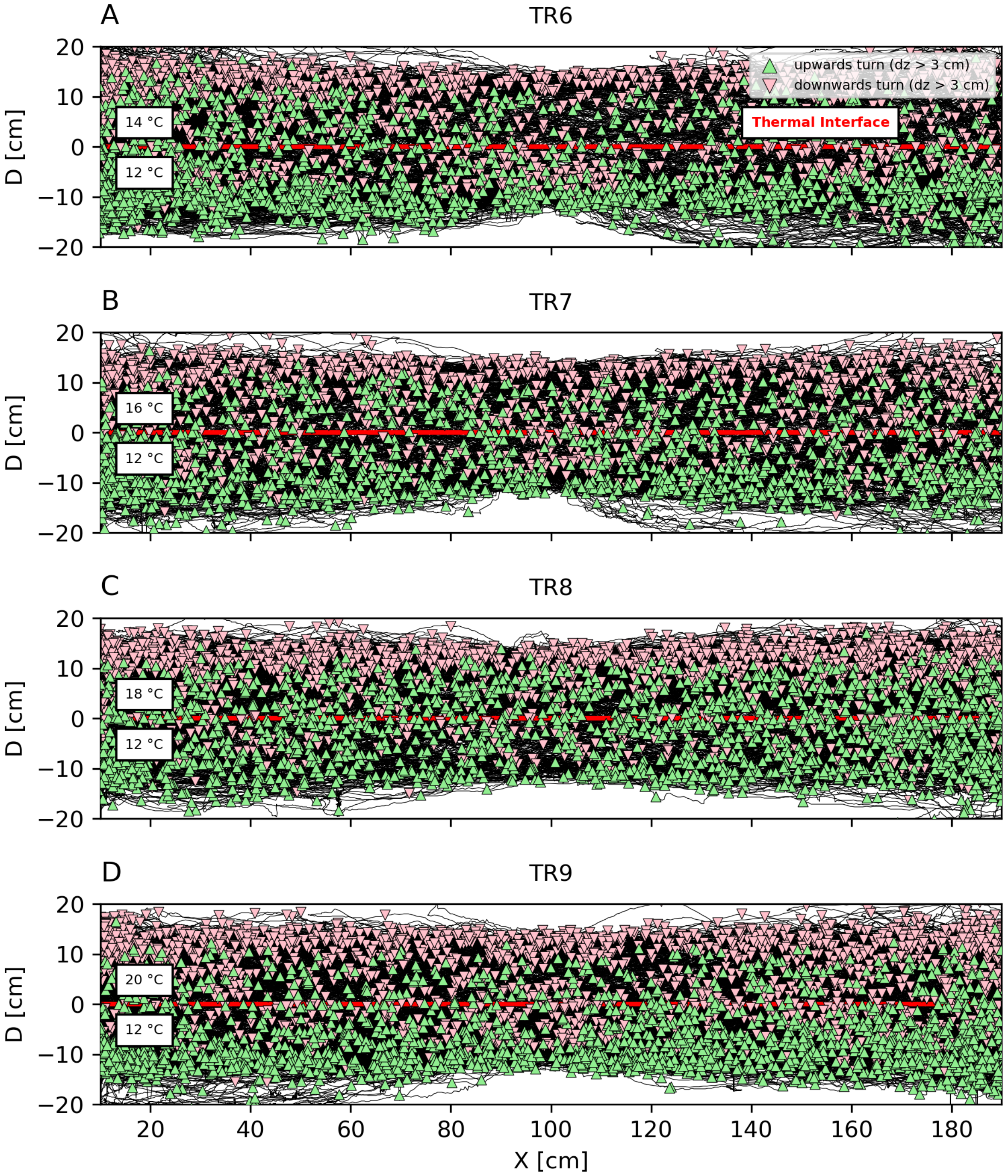


**Fig. 13:** Warm treatments. Swimming trajectories (black lines) relative to the thermal interface (red line) for all fish across cold treatments (TR6 – TR9, *A-D*). The horizontal axis depicts the horizontal position in centimetres; the vertical axis depicts the instantaneous distance from the thermal interface. For D > 0 fish is located at a distance D [cm] above the interface and for D < 0 the fish is located below the temperature interface. Red and green triangles depict large downward and upward turns, respectively (see Supplementary Fig. 11 and 14 for derivation).

#### Extracting large turning points

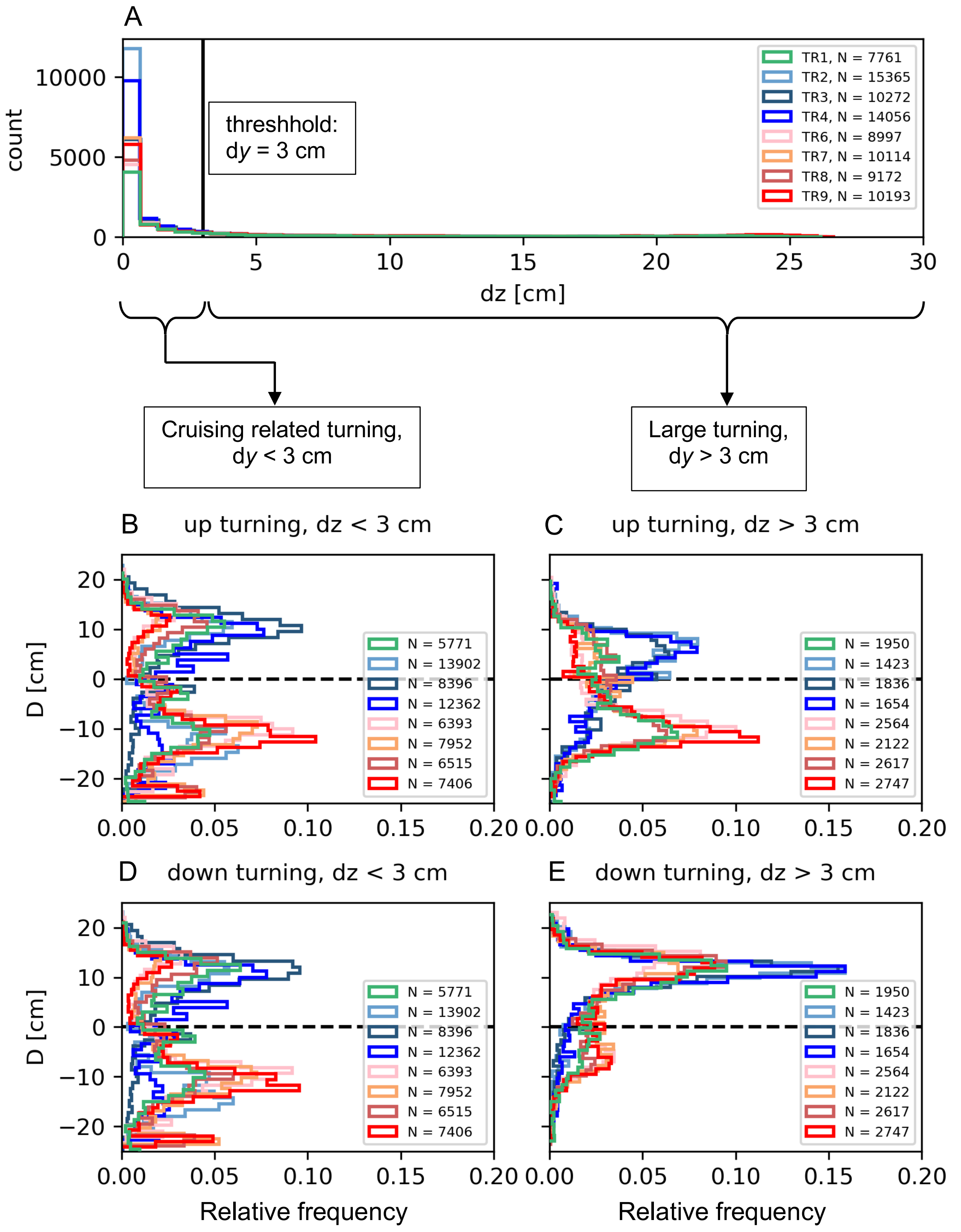


**Fig. 14:** (*A*) Normalized histogram of the vertical distance dz between all consecutive turning points for all treatments (depicted by colors). A threshold of dz = 3 cm was used to differentiate turning events that result in small, cruising related vertical displacements (*B*,*D*) from vertical turns that resulted in large vertical displacement (dz > 3 cm) (*C*, *E*).

**Number of cold and warm excursions (i.e. interface crossings)**

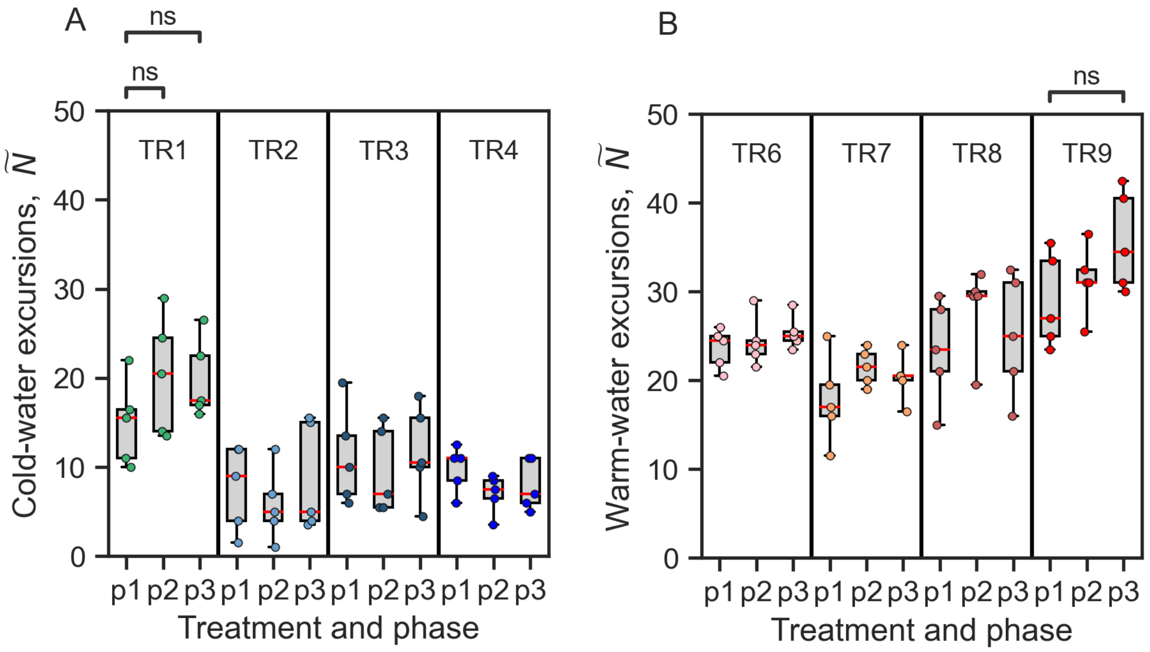


**Fig. 15**: Number of performed cold-water excursions (*A*) and warm-water excursions (*B*) over time (p1 – p3). Boxplots show the median, whiskers extend to the full data range and the boxes are limited by 25$^{th}$ and 75$^{th}$ percentiles. Each dot represents the median across 4 individuals, tested simultaneously.

#### Duration of excursions

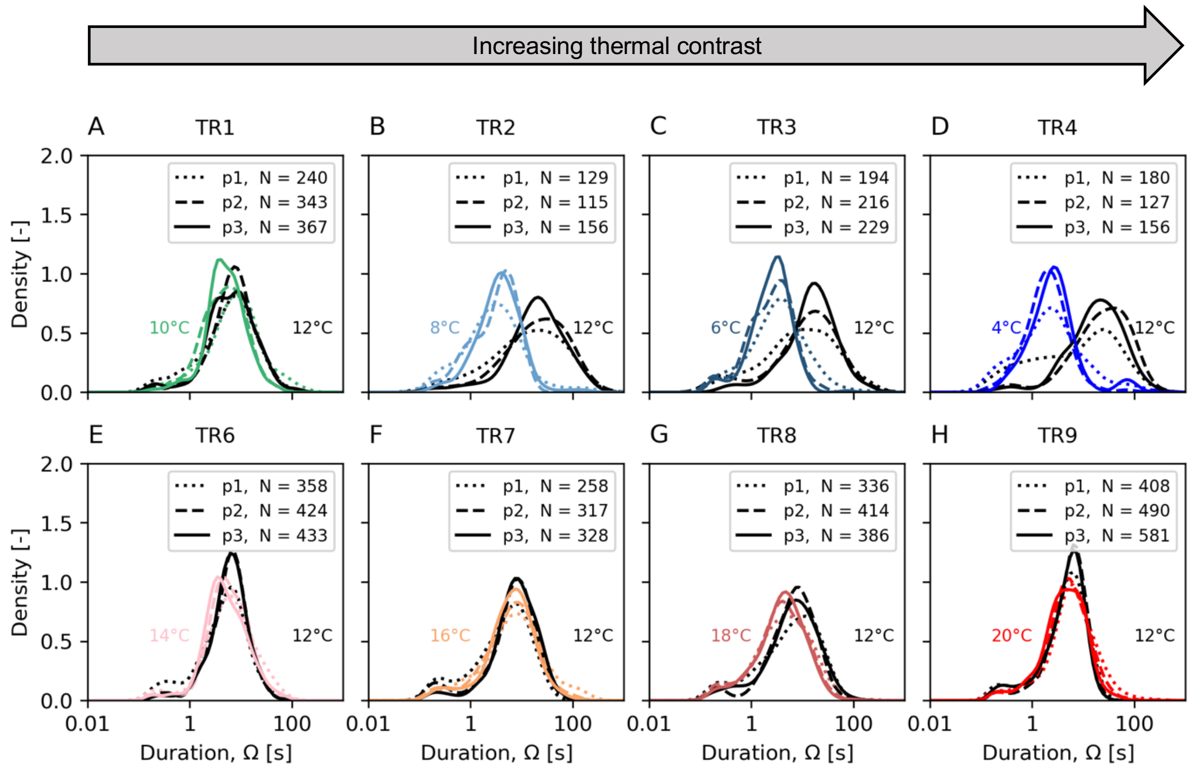


**Fig. 16:** Probability density function of the duration of warm- and cold-water excursions for cold (*A-D*) and warm treatments (*E-H*) and across treatment phases (p1, p2 and p3). As a reference for comparison, black lines depict trajectory excursions in 12°C. Coloured lines show respective distributions in treatment water temperatures below (*A-D*) or above the thermal interface (*E-H*).

#### Excursions in warm treatments

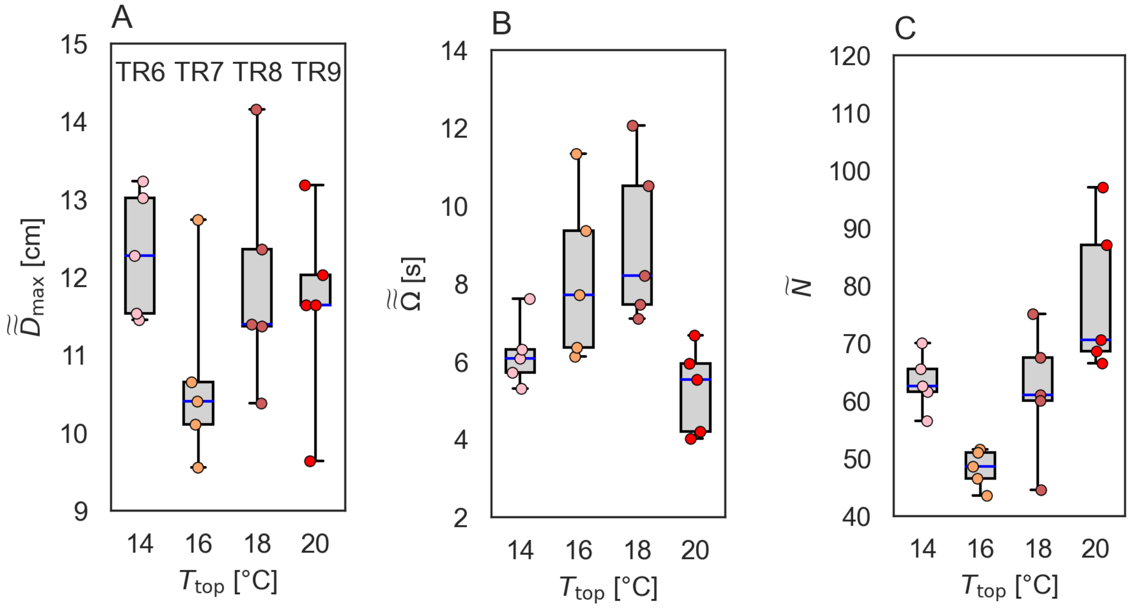


**Fig. 17:** Warm-water excursions during warm treatments (TR6-TR9). For each quantity, the median of 4 individuals (tested simultaneously) was considered; hence each dot represents a single experiment. (*A*) Boxplots of the maximum (upwards) distance to the interface of warm-water excursions. The median of $D_{max}$ was applied to all excursions performed by an individual fish and for all phases (p1, p2 and p3). (*B*) Boxplots of the median duration of cold-water excursions for fish. The median of $D_{max}$ was applied to all excursions performed by an individual fish and for all phases (p1, p2 and p3). Each dot represents a single experiment. (*C*) Boxplot of the median number of warm-water excursions (upwards) for fish groups across warm treatments and experimental phases (p1, p2 and p3). Each dot represents the median response of four fish in a single experiment. Blue lines indicate the median, boxplots are limited by the 25$^{th}$ and 75$^{th}$ percentiles. Whiskers extent to the full range of observations.

#### Maximum distance to interface

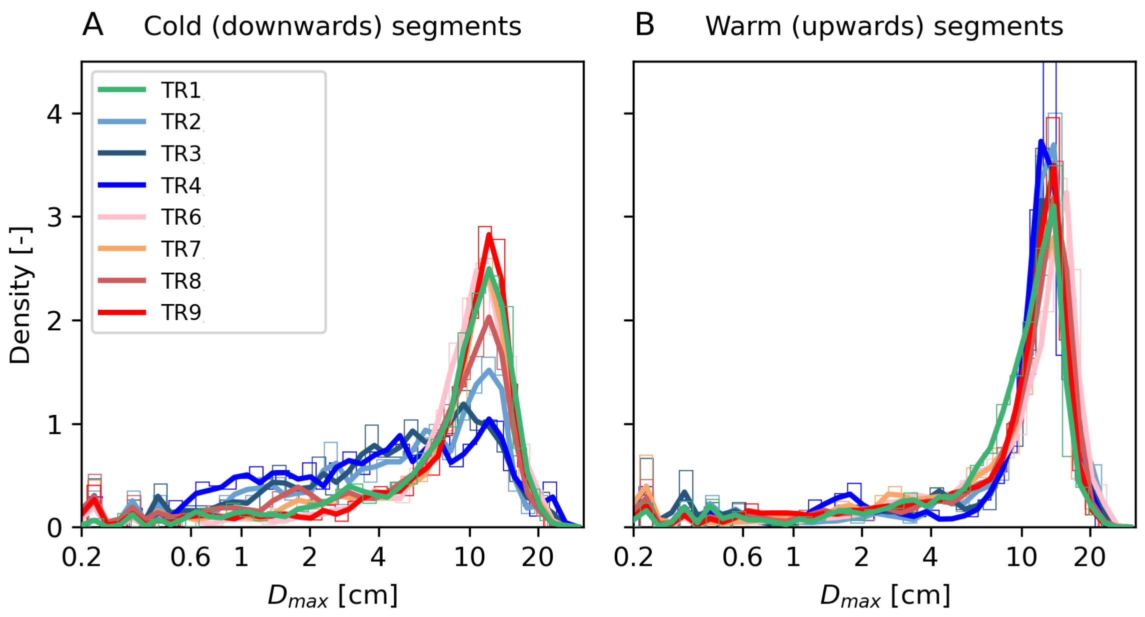


**Fig. 18:** Probability density function of maximum distance to interface line $D_{max}$ for cold (left) and warm (right) excursions for all treatments. Underlying normalized histograms are shown in thin colored lines.

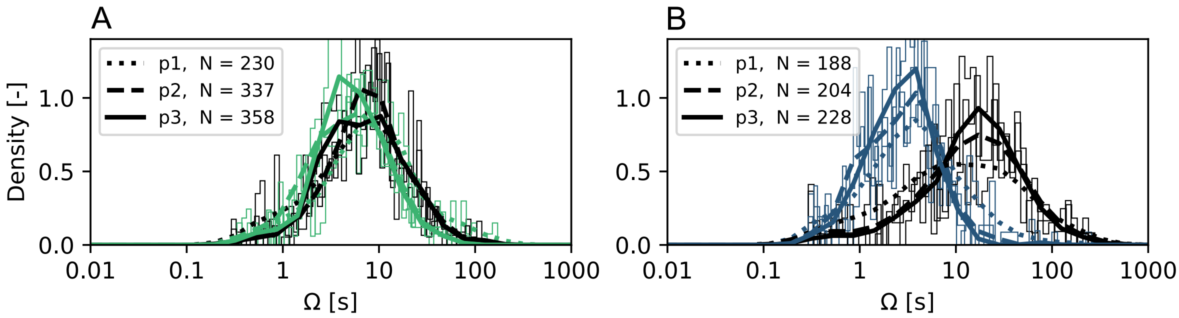


**Fig. 19**: Underlying probability density distributions of the durations *Ω* of warm- and cold-water excursions for TR1 (*A*) and TR3 (*B*), respectively. The distributions of the warm- and cold-water excursion durations become increasingly distinct over exposure time for TR3 but not TR1. Excursions at 12 °C (above the interface) are shown in black, while excursions in the lower, colder water are depicted in the respective color (green: $T_{\text{bottom}}$ = 10 °C; blue: $T_{\text{bottom}}$ = 6 °C. Treatment phases are indicated by line type (p1–p3). Values of *N* indicate the total number of excursions of each type within that phase for 20 fish (as the excursions alternate, *N* is equal for cold and warm excursions). Underlying distributions are shown as histograms.

#### Absolute Swimming speed

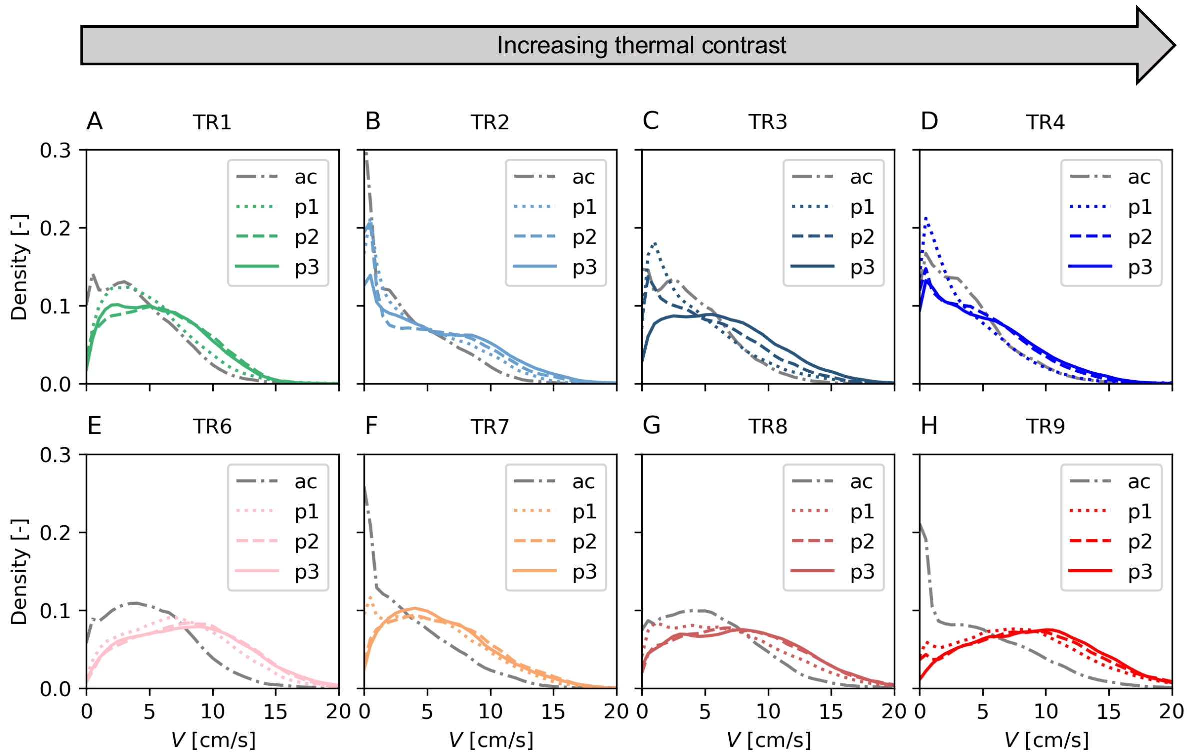


**Fig. 20:** Probability density of absolute swimming speeds *V* per experimental phase (ac: acclimation; p1, p2 and p3) and treatment. Cold treatments (A-D) and warm treatments (E-H). Each lines indicates the normalized distributions of N = 6 min x 60 s x 24 fps x 20 fish = 172’800 observations.

**Burst swimming in cold-water treatments**

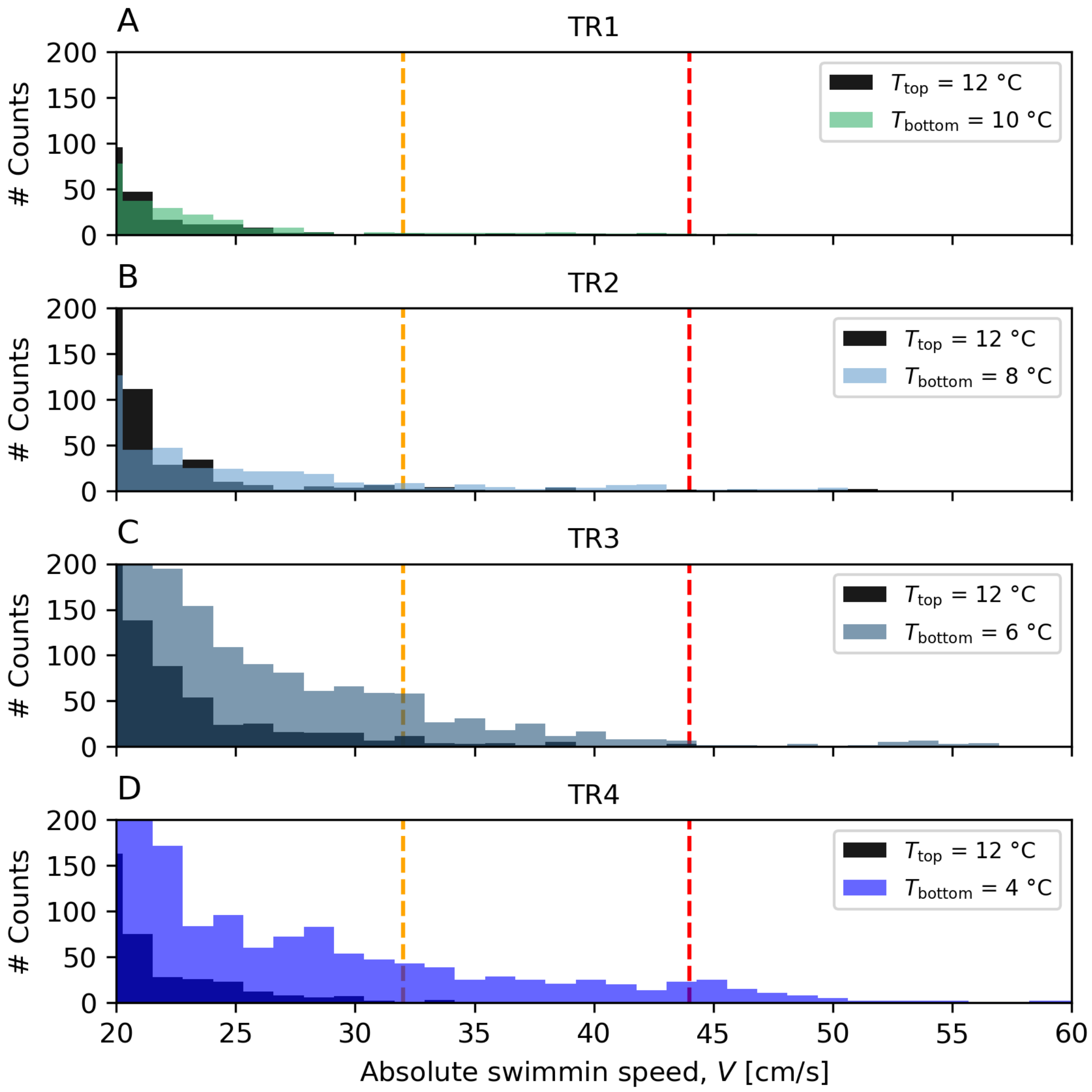


**Fig. 21**: Extreme swimming speeds above $V$ = 20 cm/s for TR1-TR4 (*A*-*D*). Black histograms show all observations above the thermal interface ($T_{top}$ = 12 °C). Colored histograms display all observations below the thermal interface (in colder water). Dashed vertical lines depict literature thresholds for burst swimming speeds of rheophilic fish based on temperature (orange line: $T$ = 12°; red line: $T$ = 4 °C), body length (3.6 cm), and swimming durations (5 seconds) as proposed by Ebel, 2014 (Supplementary Text 4). High swimming speeds were most frequent during cold-water occupation in colder treatments (TR2-TR4).

#### Normed swimming speed in warm

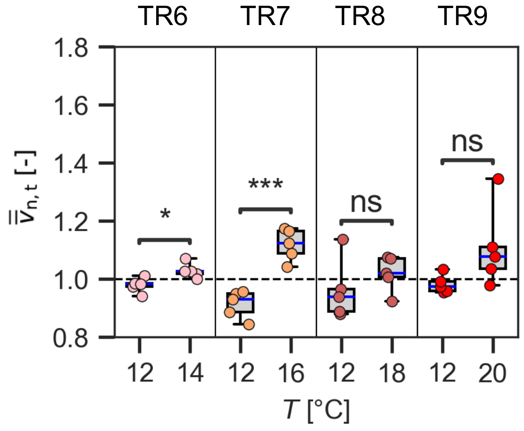


**Fig. 22:** Mean swimming speed when above (in warmer water) and below (in colder water) the thermal interface. Speed was standardized by the average swimming speed of each individual during the treatment phases (p1–p3). Data was aggregated at the replicate level; hence $\overline{\overline{v}}_{n,t}$ represents the average response of the fish group (four fish) within one replicate. Symbols represent the results of *t*-tests (ns, $p > 0.05$; *, $0.01 < p \leq 0.05$; ***, $0.0001 < p \leq 0.001$, see also Table S17). Whiskers extent to the full range of observations and red lined indicate the median.

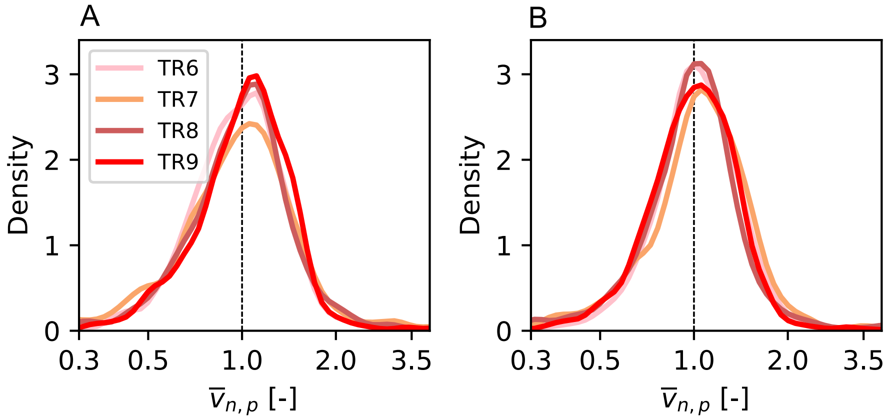


**Fig. 23**: Averaged phase normalized swimming speed during warm water excursions (*A*) and cold-water excursions (*B*) in warm treatments (TR1, TR2, TR3 and TR4). Data has been transformed to log10 and a Kernel density estimator was fitted to the normalized histogram.

#### Averaged normalized speed vs. duration of excursions

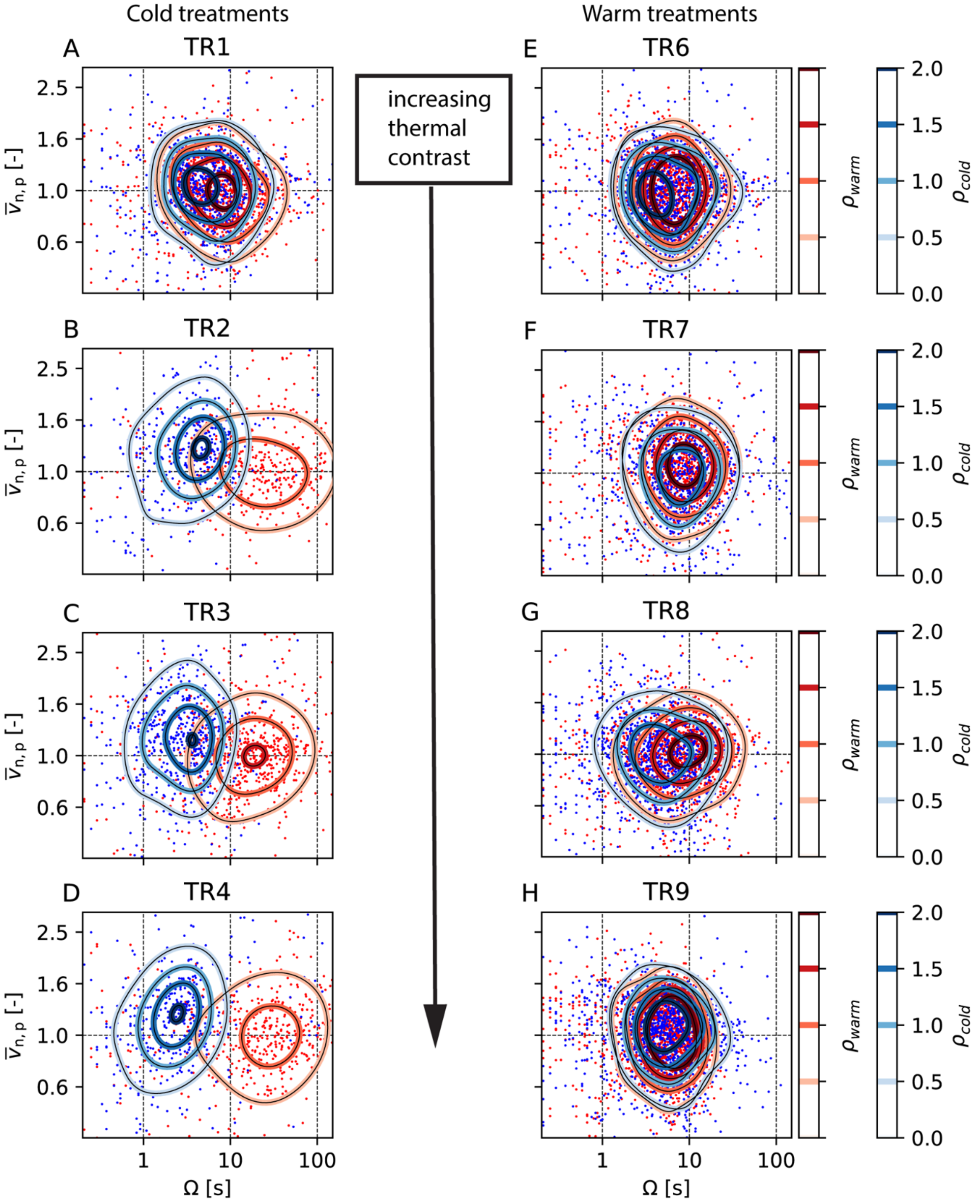


**Fig. 24:** Normalized swimming speed $\overline{v}_{n,p}$ averaged over excursions plotted as a function of the duration $\Omega$ of each cold (blue points) and warm (red points) water excursion for all cold (*A* - *D*) and all warm treatments (*E* - *H*). To control for temporal trends, speed was standardized by the average swimming speed of each individual within each phase. Fitting a 2D Kernel density estimator to the point clouds (red and blue lines) reveals for cold treatments with $T_{\text{Bottom}} \leq 8\ °C$ (TR2-TR4, *B*-*D*), clusters separate with respect to swimming speed (y-axis) and duration (x-axis).

#### Body cooling

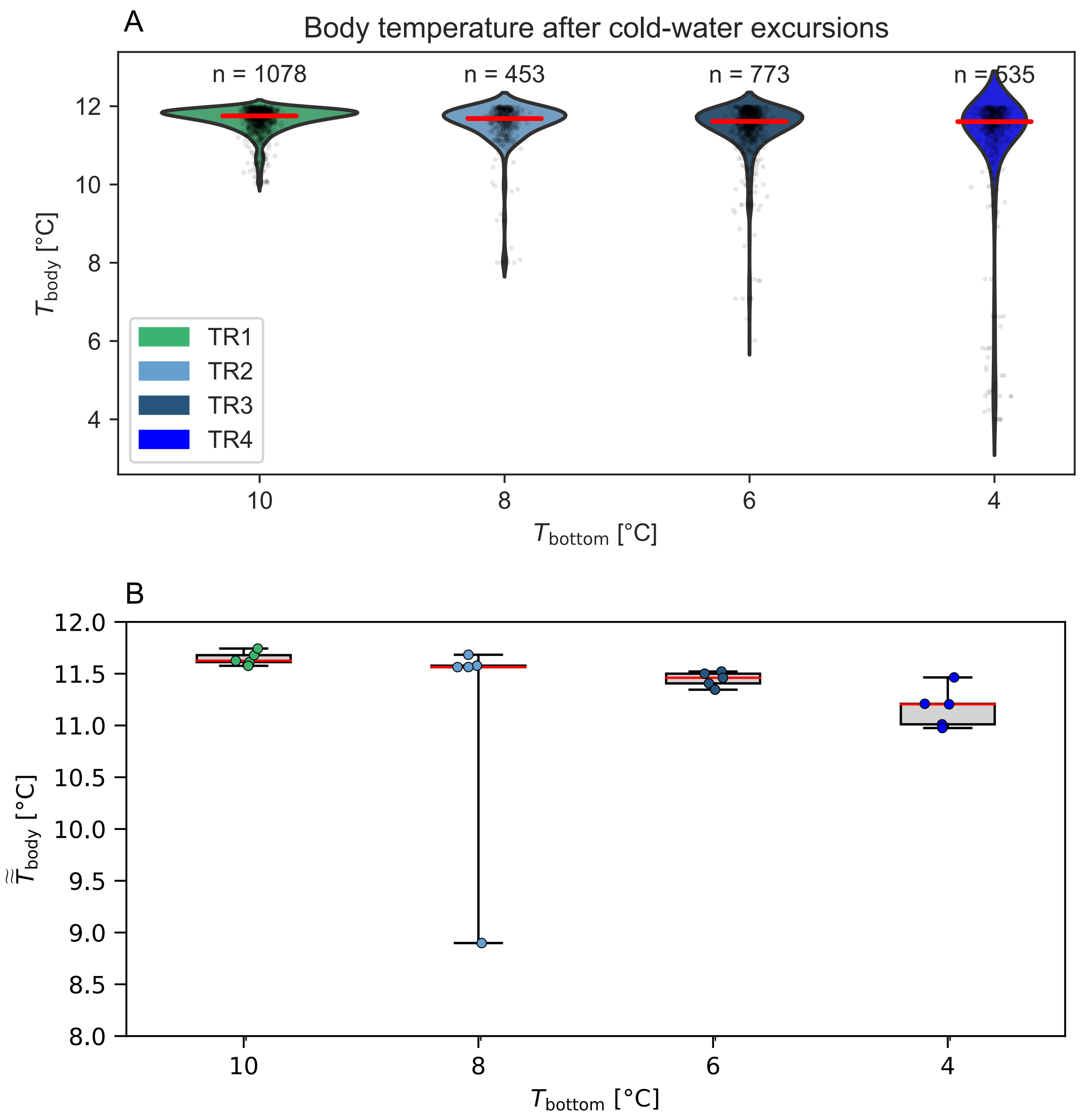


**Fig. 25:** (*A*) Violinplots of simulated body temperature $T_{body}$ after each performed cold-water excursion for cold-water treatments (TR1-TR4). Each dot is one excursion, red line indicates the median and *n* is the number of performed excursions by 5x4 = 20 fish within each treatment. The model (see SI Text) assumed fish body mass of m = 4 gramm, initial body temperature of $T_0$ = 12°C and a cooling rate coefficient of k = 1.373 min-1. (*B*) Boxplot of the median body temperature ($\tilde{\tilde{T}}_{body}$) after cold-water excursions for each experimental replicate. Red line indicates the median across experimental replicates.

#### Time spent

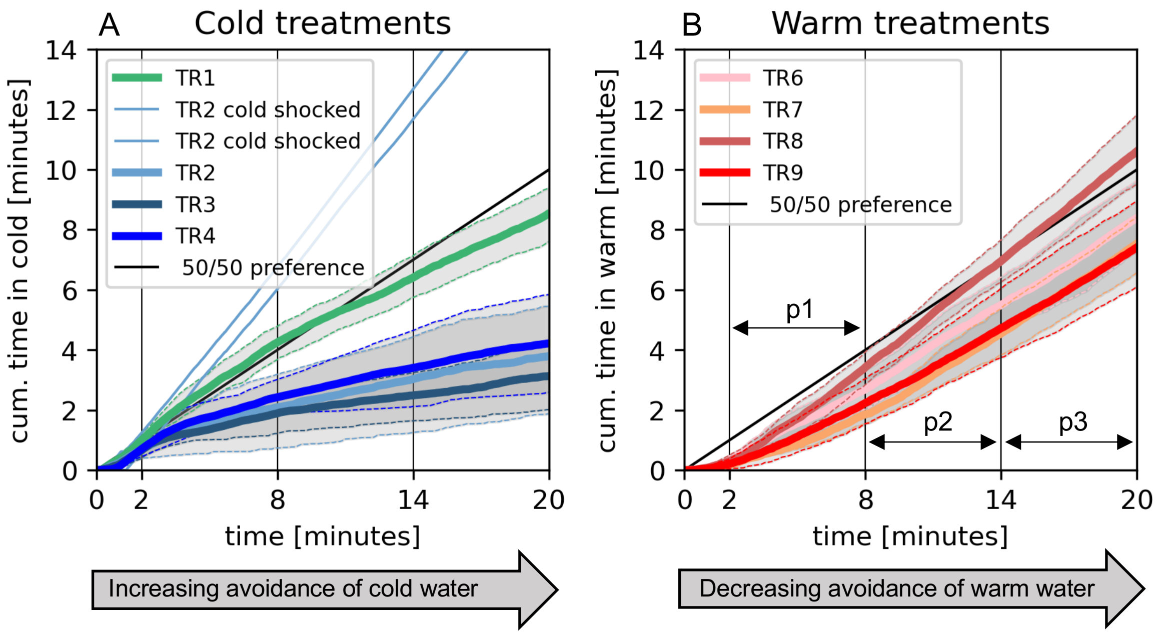


**Fig. 26:** Cumulative time spent in incoming cold water (*A*) and warm water (*B*). Thick lines represent the averaged cumulative time spent in cold water on the replicate level. Thin lines depict the 84$^{th}$and 16$^{th}$ percentile across experimental replicates. Two cold shocked individuals (in TR2) were excluded from the average as outliers. Fish displayed opposing responses to incoming warm and cold water over time. Namely, fish decreased the occupancy of cold water, while in warm treatments fish enhanced occupancy of warm water (see also Fig. 8).

#### Water temperature and dissolved oxygen

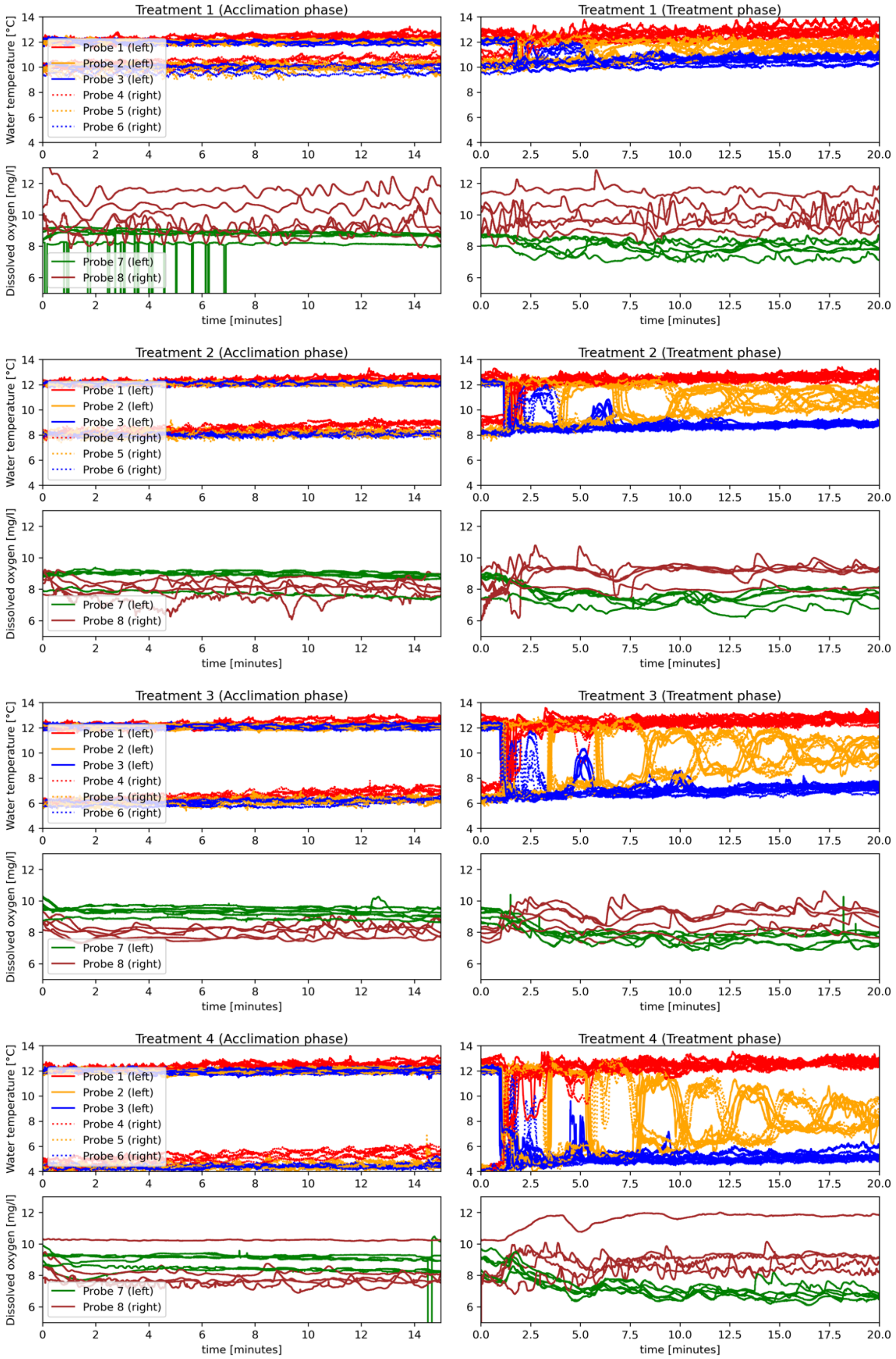


**Fig. 27: Cold treatments.** Time traces of temperature [°C] and dissolved oxygen concentration [mg/L] in experiments during acclimation (left) and treatment phase (right). Temperature (Probe 1-6) was measured at three different depths (red, orange, blue) at both side walls of the experimental tank for all experiments (upper plots). Dissolved oxygen was measured on both sides of the tank. All measurements were continuously obtained at a frequency of 5 Hz.

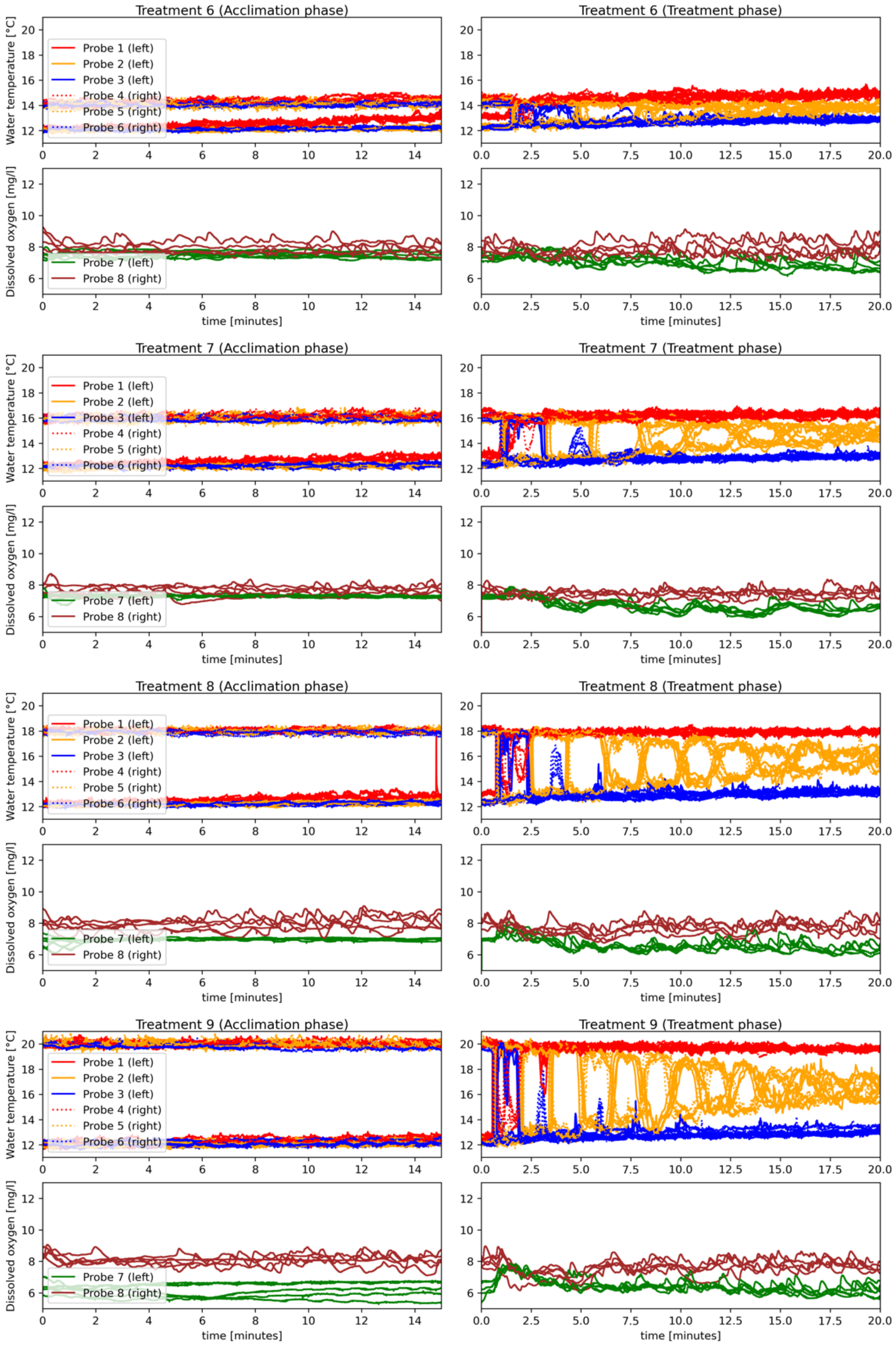


**Fig. 28: Warm treatments.** Time traces of temperature [°C] and dissolved oxygen concentration [mg/L] in experiments during acclimation (left) and treatment phase (right). Temperature (Probe 1-6) was measured at three different depths (red, orange, blue) at both side walls of the experimental tank for all experiments (upper plots). Dissolved oxygen was measured on both sides of the tank. All measurements were continuously obtained at a frequency of 5 Hz.

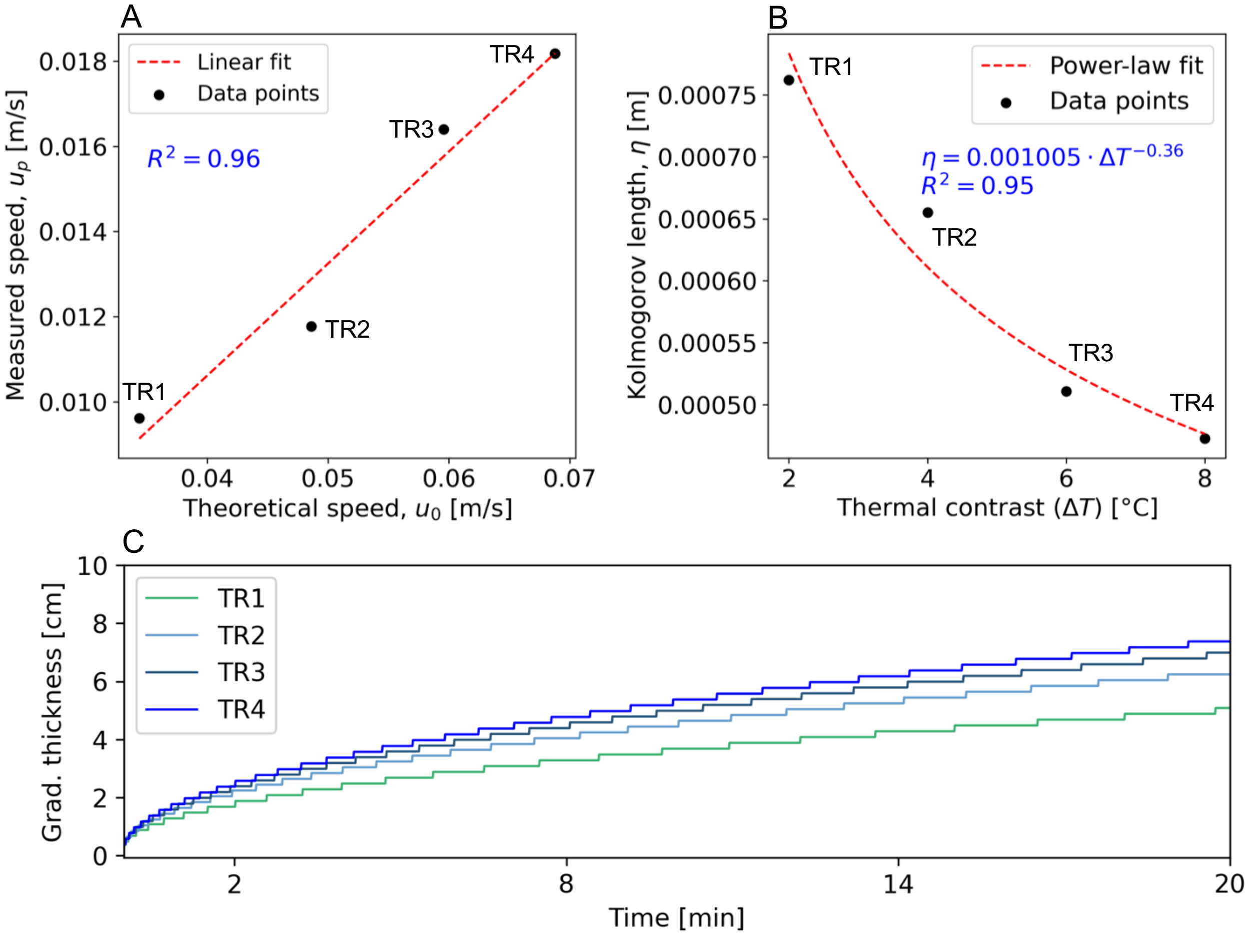


**Fig. 29:** The theoretical large-scale gravitational velocity $u_0$ plotted against the cold current propagation speed $u_p$ obtained from experimental videos of cold-water treatments (TR1–TR4). A linear regression fit (red dashed line) is shown. (B) The Kolmogorov length $\eta$ as a function of the temperature contrast $\Delta T$. The black dots represent experimental data, while the red dashed line indicates a power-law fit. (C)
Temporal evolution of the thermal interface thickness, influenced by turbulence and diffusion. A 2D numerical simulation was conducted over a 20-minute period, using an initial interface thickness estimated as $\delta_\kappa \backsimeq 3.7\ \eta$ (see Methods). The spatial domain was discretized into 1 $mm^2$ grid cells (dx = dy = 0.001 m) and a constant thermal diffusivity of D = 1.43e-07 $m^2/s$ was assumed. Results indicate that, over the course of the experiment, the gradient thickness $\delta_\kappa$increased over time, rising to values larger than the length the individual fish.

#### Trajectories

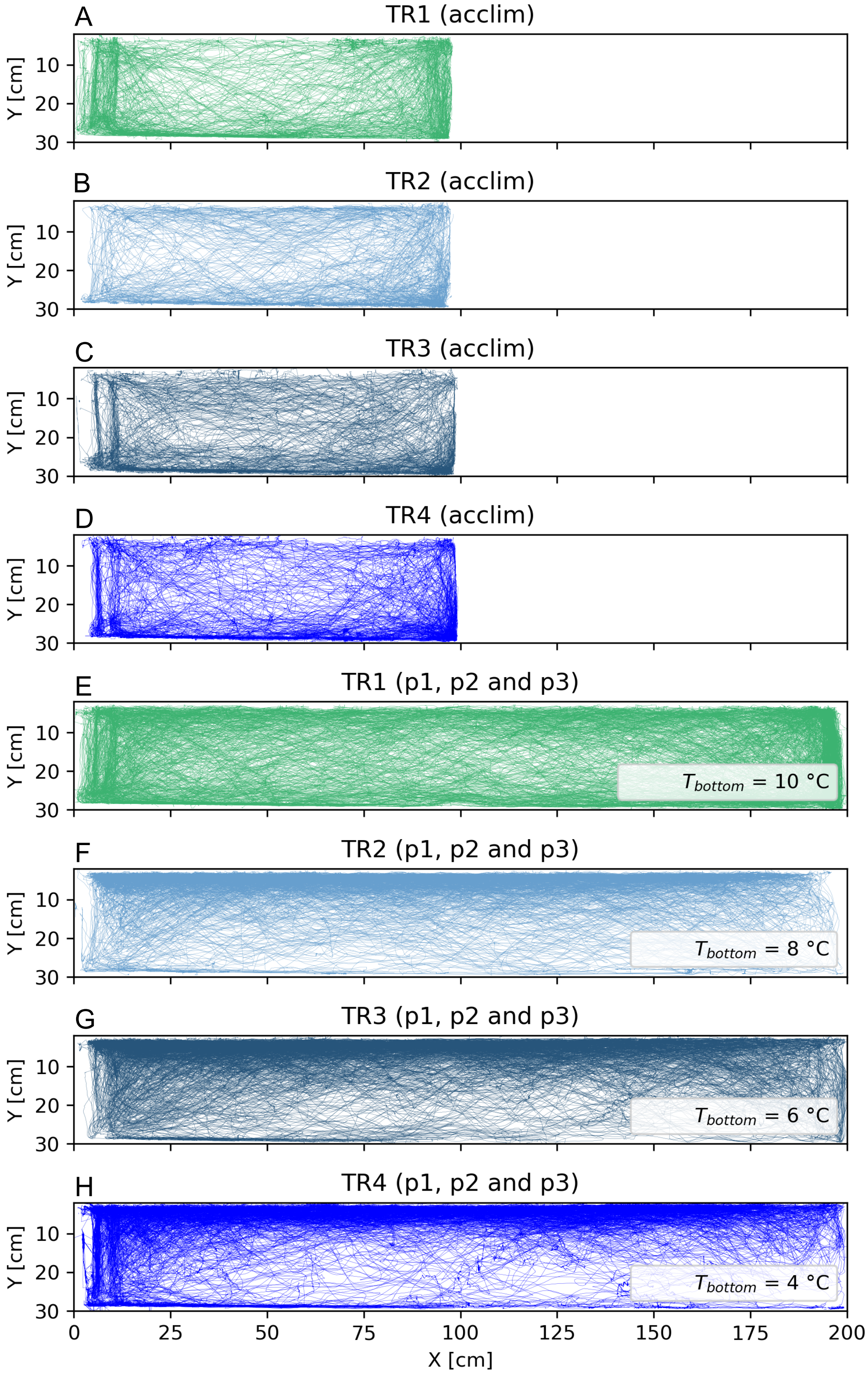


**Fig. 30:** Cold treatments. Trajectories for all fish during acclimation phase (A - D) und treatment phases (E - F).

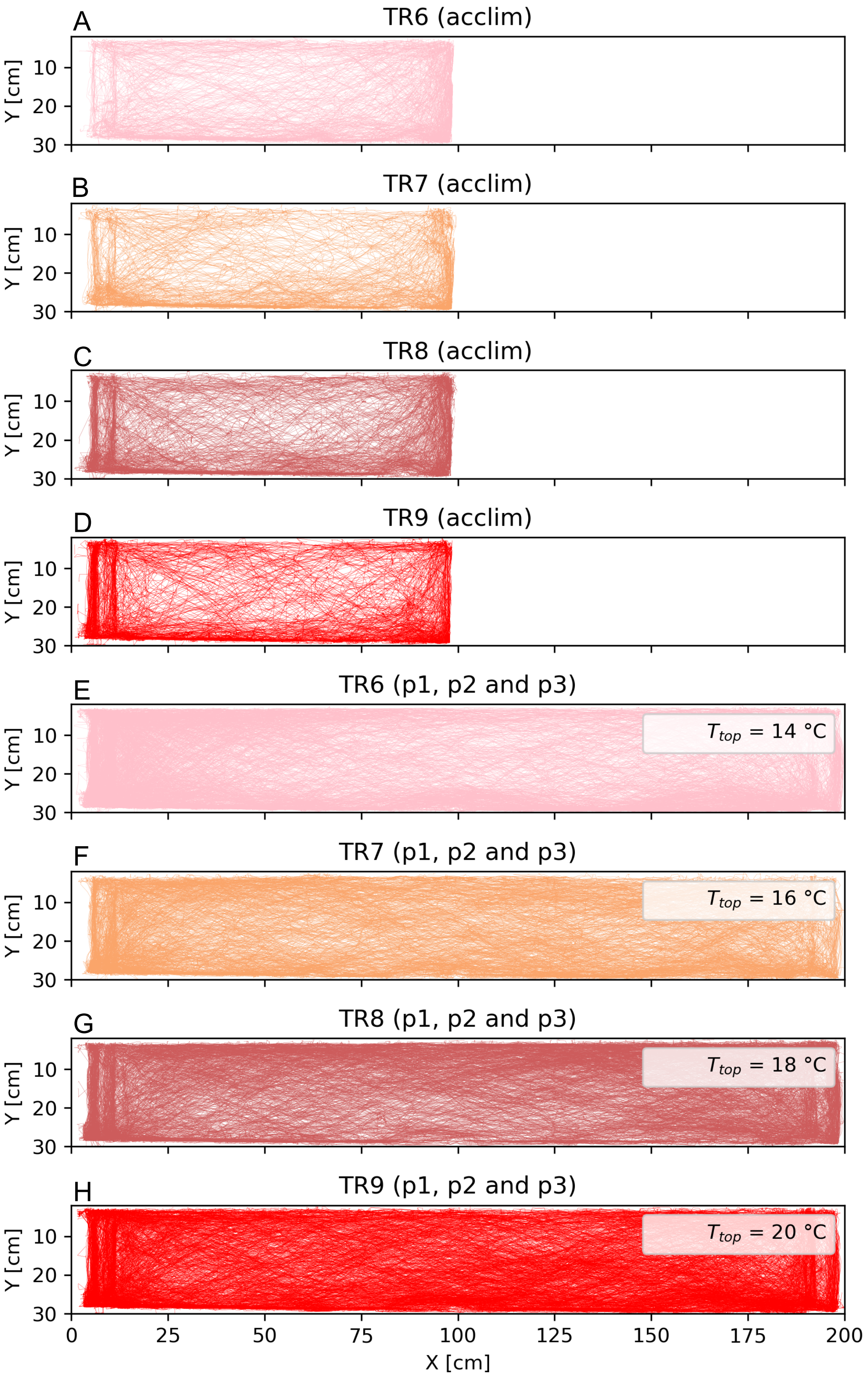


**Fig. 31:** Warm treatments. Trajectories for all fish during acclimation phase (*A* - *D*) und treatment phases (p1, p2 and p3, *E* - *F*).

#### Fish-tracking

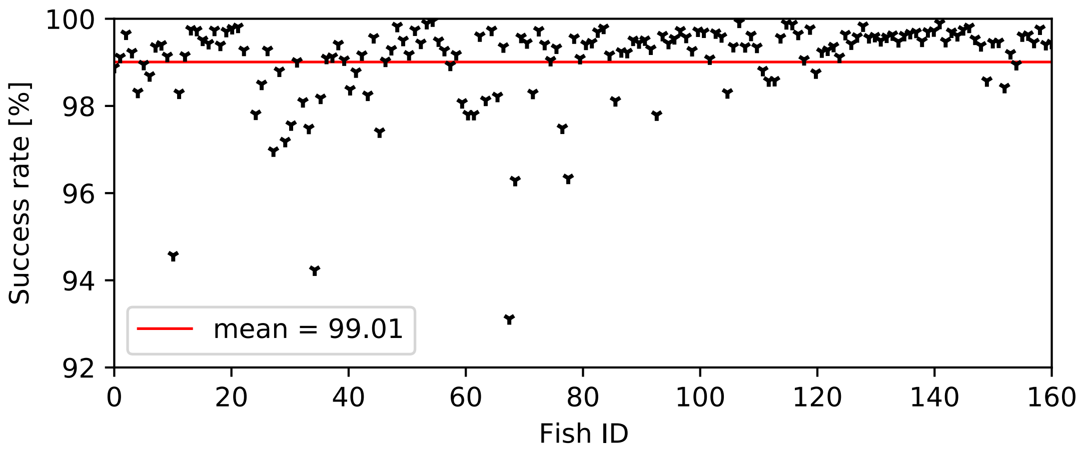


**Fig. 32:** Tracking success rate for each individual during experimental phases (p1, p2 and p3). We captured the trajectory for on average 99% of the time. Low values are explained by fish occluding over longer periods of time (without moving) or being located behind the dissolved oxygen probes. For these cases a second camera perspective was used to resolve the occlusions and data gaps were closed using linear interpolation. The open-source tracking software “TRex” (67) was used to track fish at a temporal resolution of 24 frames per second.

#### Color splitting

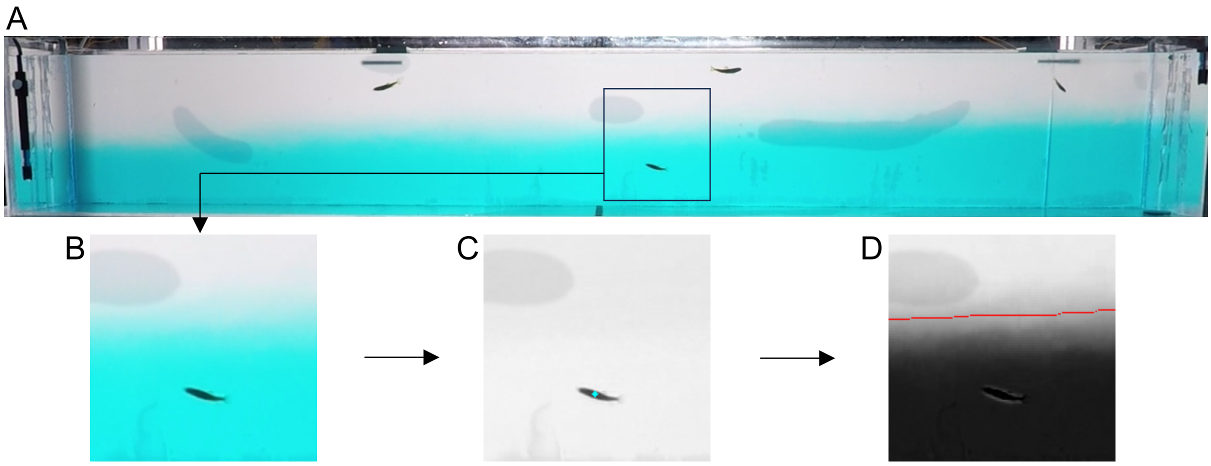


**Fig. 33:** (A) Rotated and cropped RGB image of the side camera (GoPro hero7). (B) Subsection of RGB image. (C) Red channel. The turquoise dot indicates the fish’s center of gravity for this time instance, derived from tracking in TRex. (D) Green channel**.** Red line indicates the thermal interface, derived by custom made python scripts.

#### Control experiment

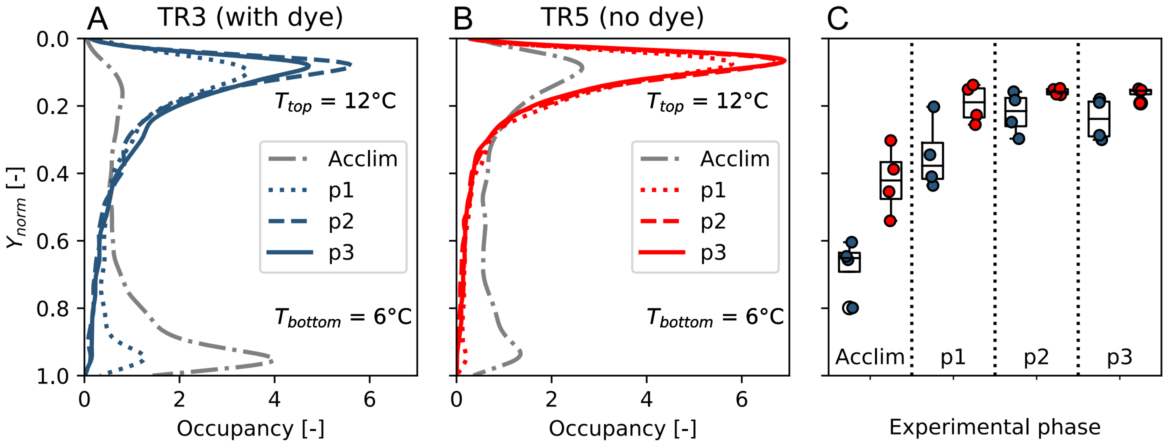


**Fig. 34:** (A *and B*) Probability density functions of fish depth during *experimental phases* (*acclim., p1, p2 and* p3) of the experiments for cold-water treatment 3 (TR3*, A, N* = 16 fish) and its control without the addition of blue dye (TR5, B, *N* = 16 fish). Both cases had identical thermal conditions ($T_{bottom}$ = 6 °C, $T_{top}$ = 12 °C) above and below the thermal interface. The depth was normalized so that $Y_{norm}$ = 0 is the water surface and $Y_{norm}$ = 1 indicates the bottom of the tank. Due to a camera failure one of the control experiments could not be processed, hence the control is based on 4 replicate*s*, while TR3 is based on 5. (C) Boxplots aggregated on the fish group level did not show significant differences with regards to the vertical relocation across experimental phases.

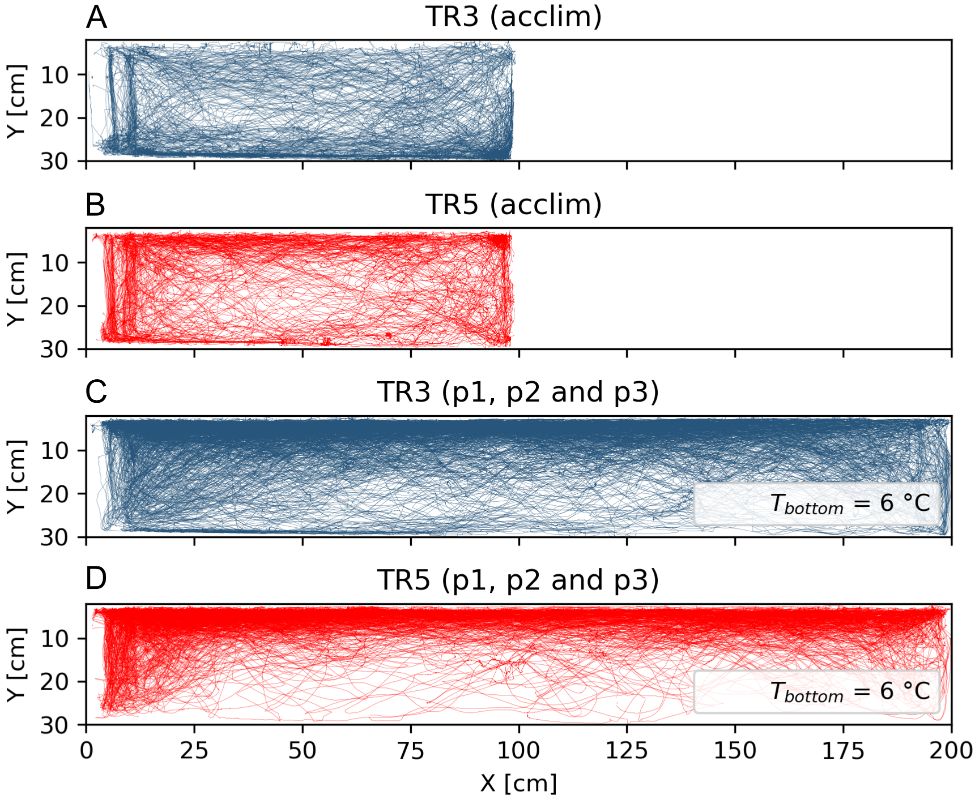


**Fig. 35:** 2D-swimming trajectories for TR3 (*A* and *C*) and its control without blue dye (TR5, *B* and *D*) during 6 min acclimation (*A* and *B*) and later experimental phases (p1, p2 and p3, *C* and *D*). Both treatments had identical temperature conditions ($T_{left}$ = 12 °C; $T_{right}$ = 6 °C).

#### Movement of thermal interface

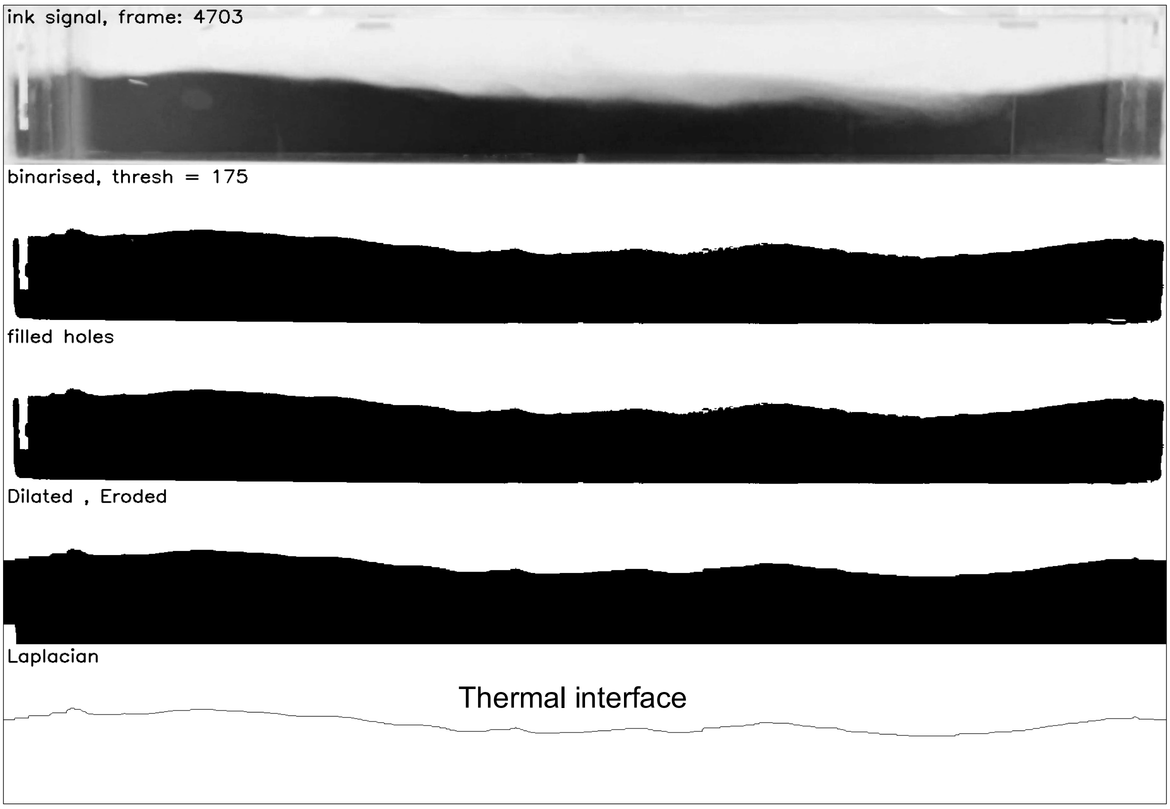


**Fig. 36:** Image operations in python (opencv) to derive the instantaneous position of the thermal interface. As the lock exchange flow is primarily a 2D phenomenon and the width of the tank is narrow (20 cm) as compared to its length (200 cm), the temperature interface was approximated as a 2D line ($X$(t), $Y$(t)). The interface coordinates were derived for each time frame using a custom-made image analysis pipeline (python, opencv).

#### Binarization of Trajectory

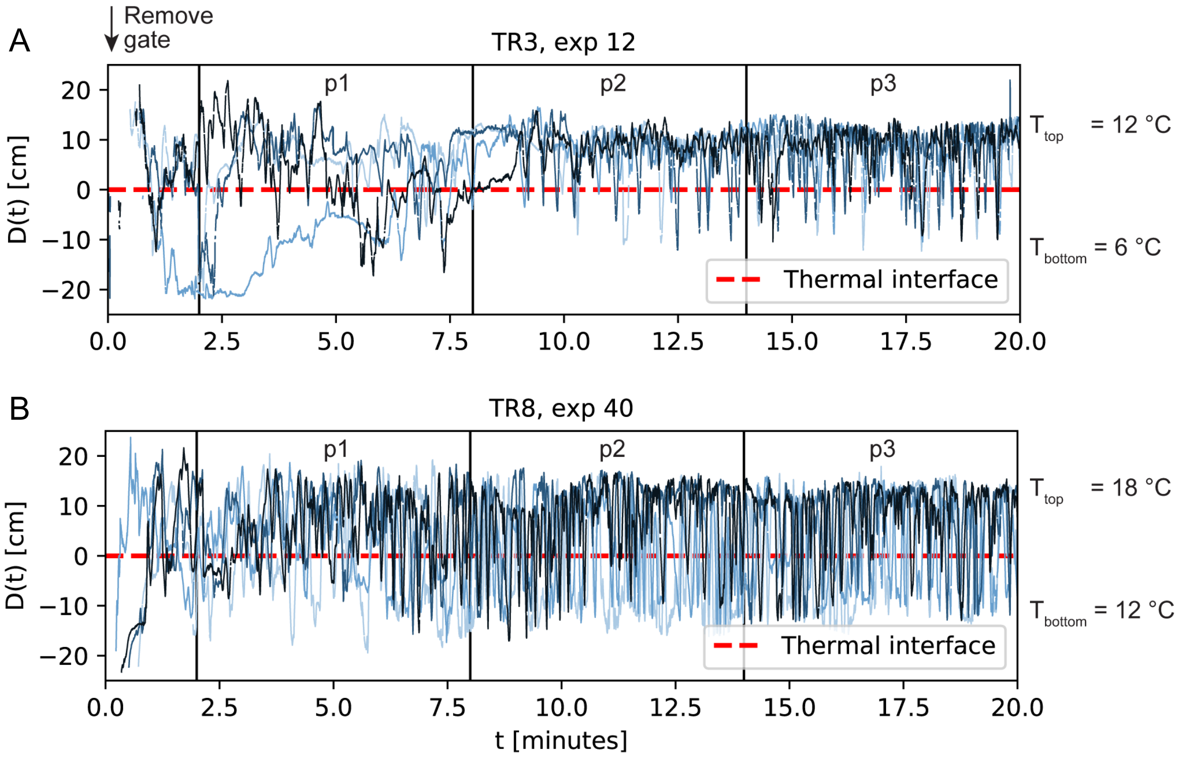


**Fig. 37: Vertical distance to thermal interface line.** The temporal progression of the vertical distance to the thermal interface line, D(t), is shown for 4 individuals in an experiment of cold treatment 3 (TR3) and warm treatment 8 (TR8). The vertical gate was removed at time $t = 0$. Negative values of D(t) depict times when the center of gravity of the tracked fish was below the thermal interface hence in the colder water and vice versa. Each sign change of D(t) depicts an interface crossing. Data gaps within the first two minutes arise from the fact that the gravity current did not yet fully span the entire horizontal dimension of the tank (i.e., in this period was excluded from our analysis, fish may neither be located below nor above but laterally displaced from the thermal interface). Shortly after the current reaches the wall, maximum distances $|D(t)|$ of up to 20 cm are possible (see also Movie 1). After the vertical stratification becomes stationary ($t \approx 10$ minutes), the interface is approximately located at half the tank depth ($h$ = 14 cm) and values for $|D(t)|$ are below 14 cm.

#### Velocity smoothing

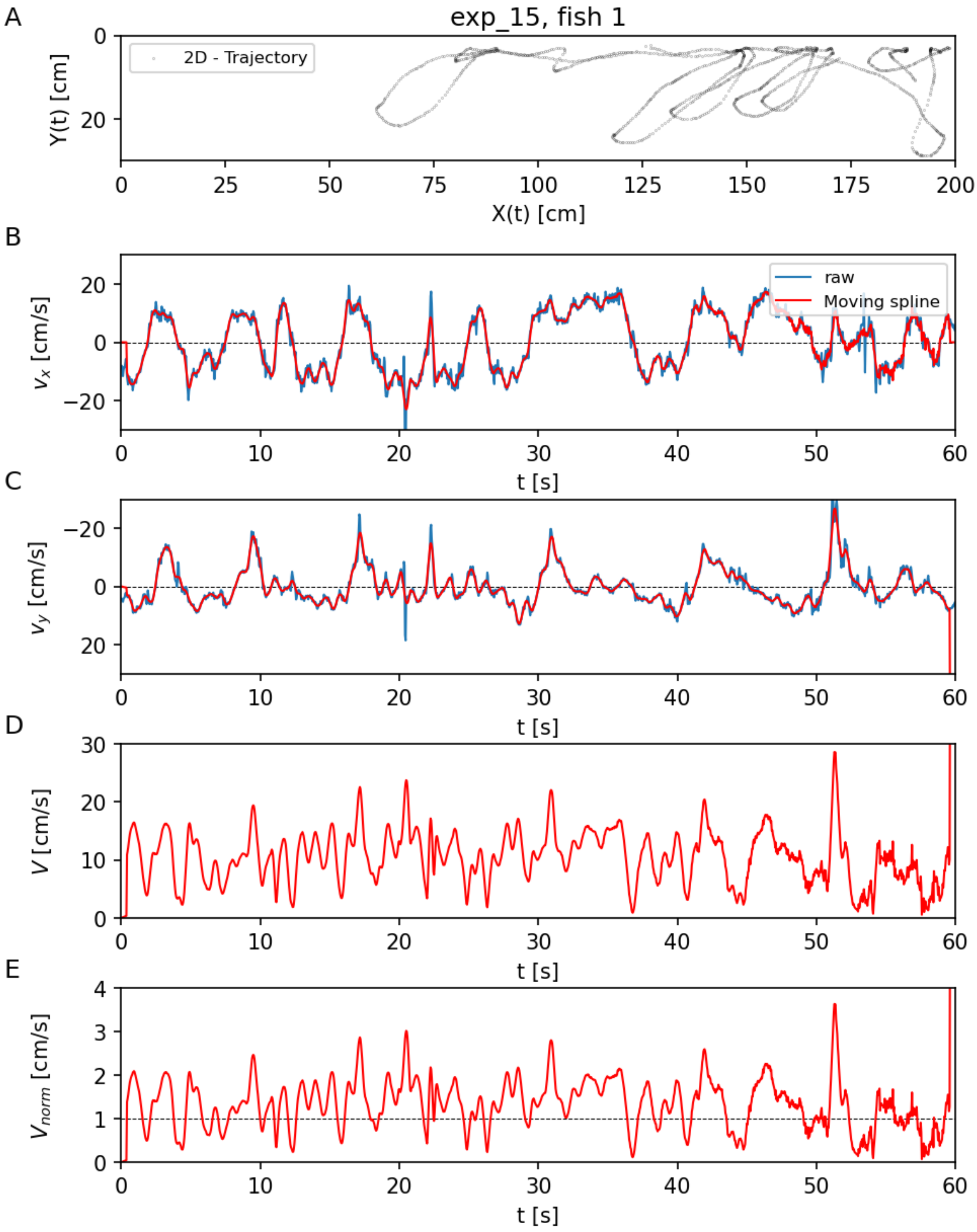


**Fig. 38:** Example time traces illustrating the derivation of fish swimming speed (*V*) and normed swimming speed $V_{\text{norm}}$ from the raw tracking output. (A) Raw trajectory output from tracking software (TRex) for a single fish during 1 minute of treatment 3 ($T_{bottom}$ = 6°C). (B,C) Horizontal and vertical velocity component ($v_x$, $v_y$) derived from raw output (blue line) and after applying the spline smoothing (Lüthi et al. 2005). (D) Absolute 2D-swimming speed *V* based on smoothed velocity components. (E) Normalized swimming speed $V_{norm}$ speed derived as $V_{norm} = V(t) / V_{mean}$. The dashed line indicates the average swimming speed during the entire treatment duration (p1, p2 and p3 combined).

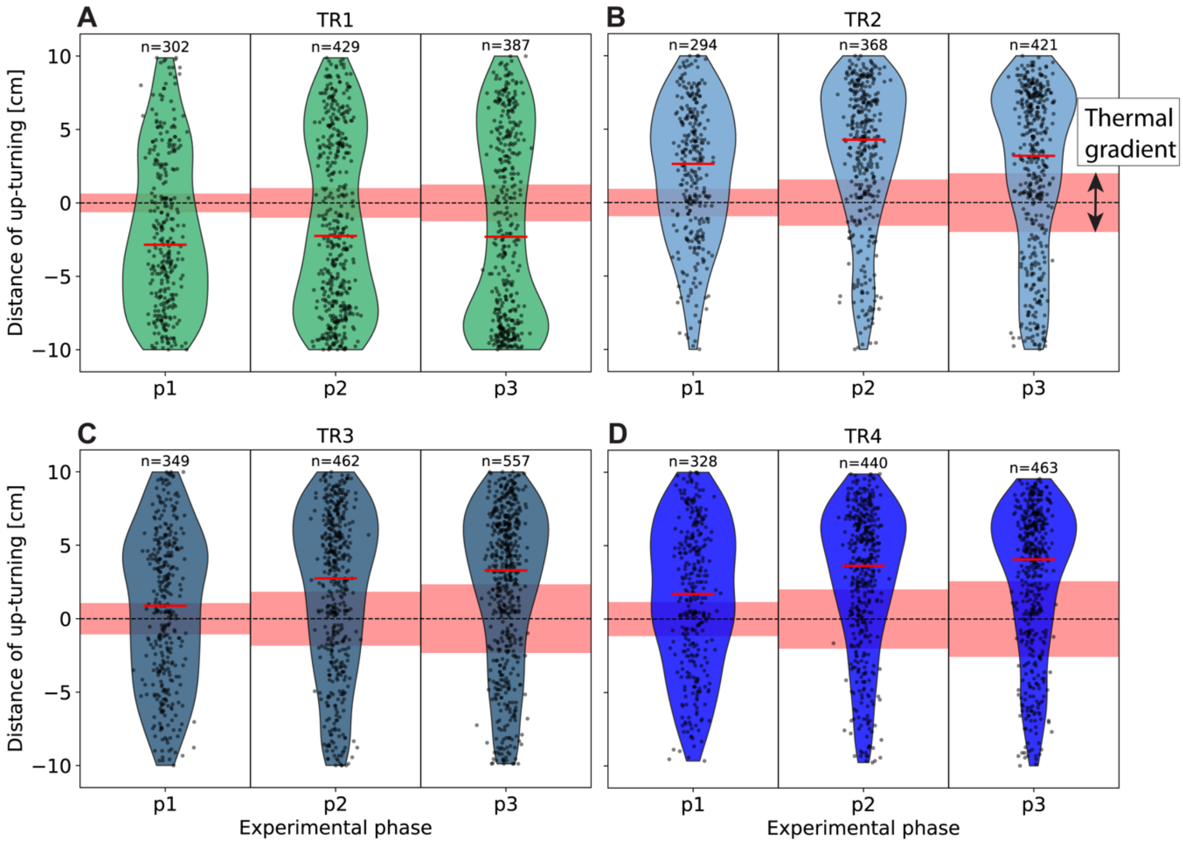


**Fig. 39**: Violin plots showing the vertical distance from the thermal interface at which up-turning events occurred during cold-water treatments (TR1–TR4; panels A–D). Each black dot represents a single up-turning event, and the red line indicates the median distance. Data are grouped by experimental phase (p1–p3). The dashed line denotes the position of the thermal interface (0 cm), and the red shaded area indicates the estimated thickness of the vertical thermal gradient (see Methods).

#### Supplementary Tables

**Table 1:** Definition of treatments. We contrasted water at the acclimation temperature (12 °C) with four cold treatments (TR1-TR4) and four warm treatments (TR6- TR9) hence we varied the incoming water temperature in both directions (Table S1). Initial water temperature ($T$) and density ($\rho$) in the left (subscript: $_{left}$) and right (subscript: $_{right}$) compartments of the tank for each treatment. Treatment 5 (TR5) was the control treatment, conducted without the dye (see Supplementary Figs. 33 and 34).

| Treatment | $T_{left}$ [°C] | $T_{right}$ [°C] | $\rho_{left}$ [kg / m$^3$] | $\rho_{right}$ [kg / m$^3$] | Density ratio: $\gamma$ [-] |
|---|---|---|---|---|---|
| TR1 | 12 | 10 | 999.5 | 999.7026 | 0.99979734 |
| TR2 | 12 | 8 | 999.5 | 999.8512 | 0.99964875 |
| TR3 | 12 | 6 | 999.5 | 999.9432 | 0.99955677 |
| TR4 | 12 | 4 | 999.5 | 999.9749 | 0.99952509 |
| TR5 (control) | 12 | 6 | 999.5 | 999.9432 | 0.99955677 |
| TR6 | 12 | 14 | 999.5 | 999.2474 | 1.00025279 |
| TR7 | 12 | 16 | 999.5 | 998.946 | 1.00055458 |
| TR8 | 12 | 18 | 999.5 | 998.5986 | 1.00090266 |
| TR9 | 12 | 20 | 999.5 | 998.2071 | 1.00129522 |

**Table 2:** Tracking parameters used for fish tracking in TRex software. Detailed description of parameter can be found here: https://trex.run/docs/parameters_trex.html

| TGrabs parameters | TRex parameters |
|---|---|
| enable_difference = true | blob_size_ranges = [[0.1,6]] |
| reset_average = true | track_max_reassign_time = 5 |
| use_adaptive_threshold = true | speed_extrapolation = 3 |
| adaptive_threshold_scale = 2 | frame_rate = 24 |
| dilation_size = -3 | gui_focus_group = [0,1,2,3] |
| track_max_individuals = 4 | heatmap_ids = [0,1,2,3] |
| average_samples = 3000 | track_max_reassign_time = 0.5 |
| averaging_method = mean | heatmap_resolution = 9 |
| threshhold = 150 | huge_timestamp_ends_segment = false |
| meta_real_width= 202 | midline_start_with_head = true |
| blob_size_ranges = [[0.1,6]] | midline_stiff_percentage = 0.4 |
| approximate_length_minutes = 15 | outline_approximate = 15 |
| cam_framerate = 24 | output_csv_decimals = 2 |
| | output_posture_data = true |
| | output_heatmaps = true |
| | posture_closing_steps = 1 |
| | track_end_segment_for_speed = false |
| | track_max_speed = 40 |
| | track_max_individuals = 4 |
| | track_max_reassign_time = 5 |
| | blob_split_max_shrink = 0.2 |
| | output_invalid_value = nan |
| | track_ignore = [[[0,0],[0,16],[1840,0],[1840,16]]] |
| | auto_quit = true |
| | gui_auto_scale_focus_one = true |

**Table 3**: Percentage of all captured cold and warm excursions shorter than 10 seconds for all phases (p1, p2 and p3) of each treatment.

| Treatment | Short cold excursions ($\Omega$ < 10 s) [%] | Short warm excursions ($\Omega$ < 10 s) [%] |
|---|---|---|
| TR1 | 75.4 | 69 |
| **TR2** | **91.2** | **38.1** |
| **TR3** | **94** | **49.1** |
| **TR4** | **92.2** | **42.7** |
| TR6 | 81 | 76.3 |
| TR7 | 73.1 | 71.4 |
| TR8 | 72.5 | 82.7 |
| TR9 | 88.2 | 81.6 |

**Table 4:** Input to body-cooling model. Summary of fish body temperature at beginning of cold-water excursion ($T_b$) and ambient water temperature ($T_a$) during cold-water excursions.

| Treatment | $T_b$ [°C] | $T_a$ [°C] |
|---|---|---|
| 1 | 12 | 10 |
| 2 | 12 | 8 |
| 3 | 12 | 6 |
| 4 | 12 | 4 |

**Table 5**: Mann-Kendall test. Input data: Median distance $\tilde{\tilde{D}}_{max}$ (in centimeters) and median durations $\tilde{\tilde{\Omega}}$ (in seconds) across all treatments and experimental replicates.

| Treatment | $T_{bottom}$ [°C] | $\tilde{\tilde{D}}_{max}$ [cm] | $Med(\tilde{\tilde{D}}_{max})$ | $\tilde{\tilde{\Omega}}$ [s] | $Med(\tilde{\tilde{\Omega}})$ |
|---|---|---|---|---|---|
| TR1 | 10 | 10.87 | | 7.02 | |
| | 10 | 9.83 | | 5.45 | |
| | 10 | 9.50 | | 6.04 | |
| | 10 | 10.37 | | 4.10 | |
| | 10 | 9.91 | 9.91 | 6.94 | 6.04 |
| TR2 | 8 | 14.52 | | 276.22 | |
| | 8 | 5.79 | | 4.04 | |
| | 8 | 7.38 | | 3.94 | |
| | 8 | 6.31 | | 4.30 | |
| | 8 | 4.97 | 6.31 | 2.88 | 4.04 |
| TR3 | 6 | 4.86 | | 3.25 | |
| | 6 | 4.97 | | 3.23 | |
| | 6 | 5.98 | | 3.48 | |
| | 6 | 3.90 | | 2.56 | |
| | 6 | 5.00 | 4.97 | 2.79 | 3.23 |
| TR4 | 4 | 4.14 | | 2.84 | |
| | 4 | 2.94 | | 2.19 | |
| | 4 | 5.27 | | 2.63 | |
| | 4 | 5.24 | | 2.59 | |
| | 4 | 4.28 | 4.28 | 1.95 | 2.59 |

**Table 6:** Mann-Kendall test result. A p-value lower than some significance level (common choices are 0.10, 0.05, and 0.01), indicates statistically significant evidence that a trend is present in the data. We found in both cases $p < 0.001$.

| Test results | $(T_{bottom}, \tilde{\bar{\Omega}})$ | $(T_{bottom}, \tilde{\bar{D}}_{max})$ |
|---|---|---|
| trend | decreasing | decreasing |
| **p-value** | 5.71E-07 | 4.27E-06 |
| Test Statistic | -292 | -263 |
| z-value | -5 | -4.60 |
| tau-value | -0.77 | -0.69 |
| s-value | -292 | -263.00 |
| slope | -0.30 | -0.36 |
| intercept | 7.85 | 9.55 |

**Table 7:** Friedman test within each treatment, < time-fraction spent in 12°C >, $T_r$ depicts the treatment temperature initially in the right compartment of the tank.

| Treatment | $T_r$ [°C] | Source | W | ddof1 | Q | p-unc |
|---|---|---|---|---|---|---|
| **TR1** | 10 | phase | 0.76 | 2 | 7.6 | **0.022*** |
| TR2 | 8 | phase | 0.04 | 2 | 0.4 | 0.819 |
| **TR3** | 6 | phase | 0.76 | 2 | 7.6 | **0.022*** |
| TR4 | 4 | phase | 0.48 | 2 | 4.8 | 0.091 |
| TR6 | 14 | phase | 0.76 | 2 | 7.6 | 0.094 |
| **TR7** | 16 | phase | 0.84 | 2 | 8.4 | **0.015*** |
| TR8 | 18 | phase | 0.52 | 2 | 5.2 | 0.074 |
| TR9 | 20 | phase | 0.28 | 2 | 2.8 | 0.247 |

**Table 8:** TR1, Dunn-test

| phase | 1 | 2 | 3 |
|---|---|---|---|
| 1 | 1.0 | **0.04*** | **0.049*** |
| 2 | | 1.0 | 1.0 |
| 3 | | | 1.0 |

**Table 9:** TR3, Dunn-test

| phase | 1 | 2 | 3 |
|---|---|---|---|
| 1 | 1.0 | 0.774 | 0.688 |
| 2 | | 1.0 | 1.0 |
| 3 | | | 1.0 |

**Table 10:** TR7, Dunn-test

| phase | 1 | 2 | 3 |
|---|---|---|---|
| 1 | 1.0 | 0.688 | 0.537 |
| 2 | | 1.0 | 1.0 |
| 3 | | | 1.0 |

**Number of cold-water excursions (Friedman and Dunn test)**

No significant reduction in the number of cold-water excursions were detected over time (see also *Supplementary* Fig. 15).

**Table 11:** Friedman test, result, median number of excursions over time for all all-treatments.

| Treatment | $T_r$ [°C] | Source | W | ddof1 | Q | p-unc |
|---|---|---|---|---|---|---|
| **TR1** | 10 | phase | 0.760 | 2 | 7.600 | **0.022** |
| TR2 | 8 | phase | 0.443 | 2 | 4.429 | 0.109 |
| TR3 | 6 | phase | 0.200 | 2 | 2.000 | 0.368 |
| TR4 | 4 | phase | 0.178 | 2 | 1.778 | 0.411 |
| TR6 | 14 | phase | 0.074 | 2 | 0.737 | 0.692 |
| TR7 | 16 | phase | 0.040 | 2 | 0.400 | 0.819 |
| TR8 | 18 | phase | 0.520 | 2 | 5.200 | 0.074 |
| **TR9** | 20 | phase | 0.640 | 2 | 6.400 | **0.041** |

**Table 12:** TR1, Dunn-test (post-hoc)

| phase | 1 | 2 | 3 |
|---|---|---|---|
| 1 | 1.0 | 0.537 | 0.312 |
| 2 | | 1.0 | 1.0 |
| 3 | | | 1.0 |

**Table 13:** TR9, Dunn-test (post-hoc)

| phase | 1 | 2 | 3 |
|---|---|---|---|
| 1 | 1.0 | 1.0 | 0.308 |
| 2 | | 1.0 | 1.0 |
| 3 | | | 1.0 |

#### Normed swimming speed (Shapiro-Wilk and t-test)

Out of 20 fish, 2 fish in TR2 were basically 'cold shocked', hence they spent very little time in the warm. One of them only spent 2 seconds above the interface. For TR2 the data was thus not normally distributed ($p > 0.05$).

**Table 14:** Shapiro-Wilk test for cold treatments. Group averaged and normed swimming speeds $\overline{\overline{v}}_{n,t}$ for different temperatures (above and below interface).

| | TR1 | | TR2 | | TR3 | | TR4 | |
|---|---|---|---|---|---|---|---|---|
| Temperature [°C] | 12 | 10 | 12 | 8 | 12 | 6 | 12 | 4 |
| Replicate | | | | | | | | |
| 1 | 1.0726 | 0.9593 | 1.4859 | 1.1885 | 0.8875 | 1.4206 | 0.9857 | 1.0257 |
| 2 | 1.0231 | 0.9907 | 0.9691 | 1.1946 | 0.9896 | 1.2103 | 0.928 | 1.489 |
| 3 | 0.9545 | 1.1524 | 0.9309 | 1.4012 | 0.9202 | 1.1159 | 1.0319 | 1.0756 |
| 4 | 1.0015 | 0.9802 | 0.981 | 1.2883 | 0.9433 | 1.3381 | 0.9652 | 1.279 |
| 5 | 0.9436 | 1.0808 | 0.9747 | 1.1839 | 0.9841 | 1.3986 | 0.917 | 1.3978 |
| p-value | 0.714 | 0.297 | **0.002** | 0.079 | 0.612 | 0.465 | 0.743 | 0.576 |
| test_stat | 0.947 | 0.877 | 0.63 | 0.799 | 0.932 | 0.909 | 0.951 | 0.927 |

**Table 15:** Shapiro-Wilk test for warm treatments. Group averaged and normed swimming speeds for different temperatures (above and below interface).

| | TR6 | | TR7 | | TR8 | | TR9 | |
|---|---|---|---|---|---|---|---|---|
| Temperature [°C] | 12 | 14 | 12 | 16 | 12 | 18 | 12 | 20 |
| Replicate | | | | | | | | |
| 1 | 1.0107 | 1.0181 | 0.9482 | 1.0876 | 0.9654 | 1.0185 | 0.9535 | 1.1097 |
| 2 | 0.9723 | 1.027 | 0.8855 | 1.1742 | 1.1352 | 0.9216 | 0.959 | 1.0755 |
| 3 | 0.985 | 1.0255 | 0.9288 | 1.1218 | 0.8776 | 1.0737 | 0.9735 | 1.3444 |
| 4 | 0.9409 | 1.0696 | 0.9546 | 1.0416 | 0.8878 | 1.0693 | 0.9914 | 1.0356 |
| 5 | 0.9811 | 0.9994 | 0.8443 | 1.1643 | 0.9375 | 1.0068 | 1.0333 | 0.9775 |
| p-value | 0.839 | 0.396 | 0.409 | 0.678 | 0.15 | 0.361 | 0.387 | 0.247 |
| test_stat | 0.965 | 0.897 | 0.9 | 0.942 | 0.834 | 0.891 | 0.896 | 0.865 |

**Table 16:** t-test results for cold treatments. ns: not significant; * : $0.01 < p \leq 0.05$; *** : $0.0001 < p \leq 0.001$

| Treatment | $T_{top}$ [°C] | $T_{bottom}$ [°C] | p-value | significance |
|---|---|---|---|---|
| TR1 | 12 | 10 | 0.4603 | ns |
| TR2 | 12 | 8 | 0.1438 | ns |
| **TR3** | 12 | 6 | 0.0004 | *** |
| **TR4** | 12 | 4 | 0.014 | * |

**Table 17:** t-test results for warm treatments.

| Treatment | $T_{top}$ [°C] | $T_{bottom}$ [°C] | p-value | significance |
|---|---|---|---|---|
| TR6 | 14 | 12 | 0.0147 | * |
| TR7 | 16 | 12 | 0.0002 | *** |
| TR8 | 18 | 12 | 0.3199 | ns |
| TR9 | 20 | 12 | 0.0859 | ns |

**Number of cold-water excursions (Shapiro-Wilk, ANOVA, t-test)**

To test whether the median number of cold-water excursions $\tilde{N}$ is normally distributed, we conducted a Shapiro-Wilk test for each of our treatments. The data of each treatment was confirmed to be normally distributed (Table S18) (i.e. p-value > 0.05) and thus, a one-way ANOVA (python: scipy.stats.f_oneway) was applied to test for significant difference between the treatments (Table S19). To find specific differences, we then applied a t-test to see which treatments are significantly different (Table 20).

**Table 18:** Median number of cold-water excursions $\tilde{N}$ of 4 fish tested simultaneously within each experimental replicate. Shapiro-Wilk-test results.

| Replicate | TR1 | TR2 | TR3 | TR4 | TR6 | TR7 | TR8 | TR9 |
|---|---|---|---|---|---|---|---|---|
| 1 | 40.5 | 2.5 | 18.0 | 21.5 | 56.5 | 48.5 | 61.0 | 70.5 |
| 2 | 61.0 | 20.5 | 24.5 | 24.5 | 61.5 | 46.5 | 44.5 | 66.5 |
| 3 | 50.0 | 31.0 | 34.5 | 28.0 | 65.5 | 51.5 | 60.0 | 68.5 |
| 4 | 69.0 | 11.5 | 43.0 | 24.0 | 62.5 | 51.0 | 75.0 | 97.0 |
| 5 | 34.5 | 32.5 | 27.5 | 16.0 | 70.0 | 43.5 | 67.5 | 87.0 |
| test_stat | 0.96 | 0.93 | 0.98 | 0.95 | 0.99 | 0.93 | 0.96 | 0.85 |
| p-value | 0.82 | 0.58 | 0.95 | 0.77 | 0.98 | 0.64 | 0.78 | 0.19 |

**Table 19:** One-way ANOVA test result

| f-statistic | p-value |
|---|---|
| 21.5 | 2.02e-10 |

**Table 20:** Pairwise t-test result

| pairs | p-value |
|---|---|
| TR1, TR2 | **0.0063** |
| TR1, TR3 | **0.0230** |
| TR1, TR4 | **0.0029** |

#### Supplementary Text

**Text 1: Gravity currents**

The difference in specific weight between warmer and colder fluid provided the driving force for the gravity currents that unfolded during the first 2 min after gate removal. After impounding at the wall, the thermal interface separated the cold water in the bottom from the warmer water in the top of the tank across the entire length $L = 2$ m of the tank. However, the thermal interface was not stationary in time. Continuously dissipating the initial energy of the system, the interface was swaying between left and right side of the tank with diminishing movement over time (Fig. 2 and Fig. 3). For all treatments, its vertical position varied more strongly across $X$ and in time during p1 and decreased for p2 and p3. The maximum movement velocity was small ($v_{interface} \approx 0.1$ cm/s) when compared to the typical swimming speed of the tested fish ($v_{Fish} \approx 2 - 8$ cm/s, Fig. 20). The effect of the flow-field was therefore neglected.

$$x_i(t) = c_{i,0} + c_{i,1}t + c_{i,2}t^2 + c_{i,3}t^2$$

are determined as

$$c_i = (\boldsymbol{A}^T x_i)(\boldsymbol{A}^T \boldsymbol{A})^{-1} ,$$

where

$$\boldsymbol{A} = \begin{bmatrix} 1 & (t-10\cdot\Delta\mathrm{t}) & (t-10\cdot\Delta\mathrm{t})^2 & (t-10\cdot\Delta\mathrm{t})^3 \\ 1 & (t-9\cdot\Delta\mathrm{t}) & (t-9\cdot\Delta\mathrm{t})^2 & (t-9\cdot\Delta\mathrm{t})^3 \\ \dots & \dots & \dots & \dots \\ 1 & (t+10\cdot\Delta\mathrm{t}) & (t+10\cdot\Delta\mathrm{t})^2 & (t+10\cdot\Delta\mathrm{t})^3 \end{bmatrix}$$

and

$$x_i = \begin{bmatrix} x_i(t-10\cdot\Delta\mathrm{t}) \\ x_i(t-9\cdot\Delta\mathrm{t}) \\ \dots \\ x_i(t+10\cdot\Delta\mathrm{t}) \end{bmatrix}$$

The filtered position, velocity and acceleration, $\hat{x}_i(t), \hat{u}_i(t)\ and\ \hat{a}_i(t)$, where then obtained per component as:

$$\hat{x}_i(t) = c_{i,0} + c_{i,1}(t) + c_{i,2}(t^2) + c_{i,3}(t^3),$$

$$\hat{u}_i(t) = c_{i,1} + 2c_{i,2}t + 3c_{i,3}t^2$$

$$\hat{a}_i(t) = 2c_{i,2} + 6c_{i,3}t$$

the swimming speed $\hat{v}(t)$ was then derived as:

$$\hat{v}(t) = \sqrt{\left(\hat{u}_1(t)\right)^2 + \left(\hat{u}_2(t)\right)^2}$$

Finally, an individuals' normalized swimming speed $v_{norm}(t)$ is calculated by division through the individuals' temporally averaged speed during the treatment phase, $\bar{\hat{v}}$:

$$v_{norm}(t) = \frac{\hat{v}(t)}{\bar{\hat{v}}}$$

The effect of the moving spline on the raw velocity components and the derivation of $\hat{v}(t)$ and $v_{norm}(t)$ is visualized in Supplementary Fig. 38 A-E and in Movie 5.

#### Text 4: Body cooling during cold-water excursions

Parameters and equation from [39].

a = 2.267
b = -0.329
$m_{body}$ = 4 [gramm]

Heat transfer rate:

$$k = b * m_{body}^{\ b} = 1.373 \ [1/minute]$$

Fish body temperature change per time:

$$\frac{T_b}{dt} = k * (T_a - T_b)$$

Fish body temperature after cold water excursion with duration Ω:

$$T_b(\Omega) = (T_0 - T_a)\, e^{-k\Omega} + T_a$$

#### Text 4: Burst swimming model

We compared our observed maximum swimming speeds to a multivariate swimming speed model for rheophilic fish species [77]. Thereby fish burst swimming speed ($u$) was calculated for a duration of $t$ = 5 seconds, body length of $BL$ = 0.036 m and temperatures of $T$ = 12 °C and $T$ = 6 °C (Supplementary Fig. 21).

$$\log(u) = 0{,}5460 + 0{,}7937 \cdot \log(BL) - 0{,}0902 \cdot \log(t) + 0{,}2813 \cdot \log(T)$$

The direct comparison of our observed absolute swimming speeds to the model is however difficult because:

1. The model is based on swimming speed derived from respirometers where a steady flow is present. The absence of flow in our experiments does not compare to this and fish inherently displays lower swimming speeds.
2. The confined space and the presence of walls forces fish to decelerate frequently, preventing it from reaching very high swimming speeds.
3. We are looking at the 2D-projection of the swimming trajectory, thus our observations neglect the 3rd velocity component.

#### Supplementary Movies

All movies are played at 20x real speed. Original framerate was 24 frames per second.

**Movie 1: Tracking of 1 fish in cold treatment (TR2).**
Side-view of the tank with individual fish swimming-trajectory and gravity current for single experimental replicate (TR2, cold-water in light blue). Individual fish movement trajectory is shown in red and its instantaneous vertical distance to the thermal interface *D(t)* in the lower plot for the progression of the entire treatment (p1, p2 and p3) immediately after gate-removal. Playback is 20x real speed.

**Movie 2: Tracking of 4 fish in cold treatment (TR2).**
Same for all 4 individuals.

**Movie 3: Tracking of 1 fish in warm treatment (TR7)**
Side-view of the tank with individual fish swimming-trajectory and gravity current for single experimental replicate (TR7, warmer water in light blue). Individual fish movement trajectory is shown in red and its instantaneous vertical distance to the thermal interface *D(t)* in the lower plot for the progression of the entire treatment (p1, p2 and p3) immediately after gate-removal. Playback is 20x real speed.

**Movie 4: Tracking of 4 fish in warm treatment (TR7).**
Same for all 4 individuals.

**Movie 5: Derivation of normed swimming speed**
Derivation of normed swimming speed for trajectory segment of 60 seconds. (*A*) Single fish trajectory. (*B*) and (*C*) horizontal and vertical velocity components. (*D*) Absolute swimming speed. (*E*) Normed swimming speed.
